# Focused Ultrasound Modulation of Hepatic Neural Plexus Restores Glucose Homeostasis in Diabetes

**DOI:** 10.1101/2021.04.16.440207

**Authors:** Victoria Cotero, Hiromi Miwa, Zall Hirschstein, Khaled Qanud, Tomás S. Huerta, Ningwen Tai, Yuyan Ding, Kevin Jimenez-Cowell, Jacquelyn-Nicole Tomaio, Weiguo Song, Alex Devarajan, Tea Tsaava, John Graf, Radhika Madhavan, Kirk Wallace, Evelina Loghin, Christine Morton, Ying Fan, Tzu-Jen Kao, Kainat Akhtar, Meghana Damaraju, Linda Barenboim, Teresa Maietta, Jeffrey Ashe, Kevin J. Tracey, Thomas R. Coleman, Dino Di Carlo, Damian Shin, Stavros Zanos, Sangeeta S. Chavan, Raimund I. Herzog, Chris Puleo

**Affiliations:** General Electric (GE) Research, 1 Research Circle, Niskayuna, NY, 12309; Albany Medical College, Albany, NY 12208; Feinstein Institutes for Medical Research, Manhasset, NY, 11030; Yale School of Medicine, New Haven, CT, 06519; University of California, Los Angeles, Los Angeles, CA, 90095

## Abstract

While peripheral glucose sensors are known to relay signals of substrate availability to integrative nuclei in the brain, the importance of these pathways in maintaining energy homeostasis and their contribution to disease remain unknown. Herein, we demonstrate that selective activation of the hepatoportal neural plexus via transient peripheral focused ultrasound (pFUS) induces glucose homeostasis in models of well-established insulin resistant diabetes. pFUS modulates sensory projections to the hindbrain and alters hypothalamic concentrations of neurotransmitters that regulate metabolism, resulting in potentiation of hypothalamic insulin signaling, leptin-independent inhibition of the orexigenic neuropeptide Y system, and therapeutic alteration in autonomic output to peripheral effector organs. Multiomic profiling confirms pFUS-induced modifications of key metabolic functions in liver, pancreas, muscle, adipose, kidney, and intestines. Activation of the hepatic nutrient sensing pathway not only restores nervous system coordination of peripheral metabolism in three different species but does so across these organ systems; several of which are current targets of antidiabetic drug classes. These results demonstrate the potential of hepatic pFUS as a novel/non-pharmacologic therapeutic modality to restore glucose homeostasis in metabolic diseases, including type II diabetes.

**One Sentence Summary:** We utilize a non-invasive ultrasound technique to activate a liver-brain sensory pathway and demonstrate its potential to induce durable normalization of glucose homeostasis in models of well-established insulin resistant diabetes.

## Ultrasound Stimulation of the Hepatic Neural Plexus Improves Insulin Resistance

We utilized intrahepatic focused pulsed ultrasound stimulation (pFUS)^1^ to modulate the liver-brain neurometabolic pathway^2–12^ in several animal models of hyperglycemia and diabetes (Fig. 1A and B; Supplemental Fig. S1-S3). The pFUS stimuli^1,13–15^ were non-invasively applied to the porta hepatis using ultrasound image-targeting^1^ (Supplemental Fig. S2). The stimulus parameters were based on the dose (i.e., ultrasound frequency, pulse repetition period, pulse length and magnitude, and treatment duration) associated with maximal reduction of blood glucose in an acute rat model of hyperglycemia^1^ (Supplemental Fig. S3). Application of the ultrasound stimulus in both genetic and diet-induced type II diabetes (T2D) models resulted in significantly lower plasma glucose levels (i.e., area under the curve, AUC) compared to sham treatment controls in glucose tolerance tests (GTTs; Fig. 1C and D). A single ultrasound stimulation of three minutes improved both circulating glucose (1C i. and ii.) and insulin concentrations, resulting in improved homeostatic model assessment of insulin resistance (HOMA_IR) scores in the Zucker diabetic fatty rat model (ZDF; 1C iii. and Supplemental Fig. S4). In a modified western diet-induced obesity (WD) mouse model, this effect was shown to be sustainable and resulted in maintenance of significantly lower fasting insulin levels and HOMA IR score after 8 weeks of daily ultrasound stimulation compared to sham controls (Fig. 1D and Supp. Fig. S5).

**Figure 1.**
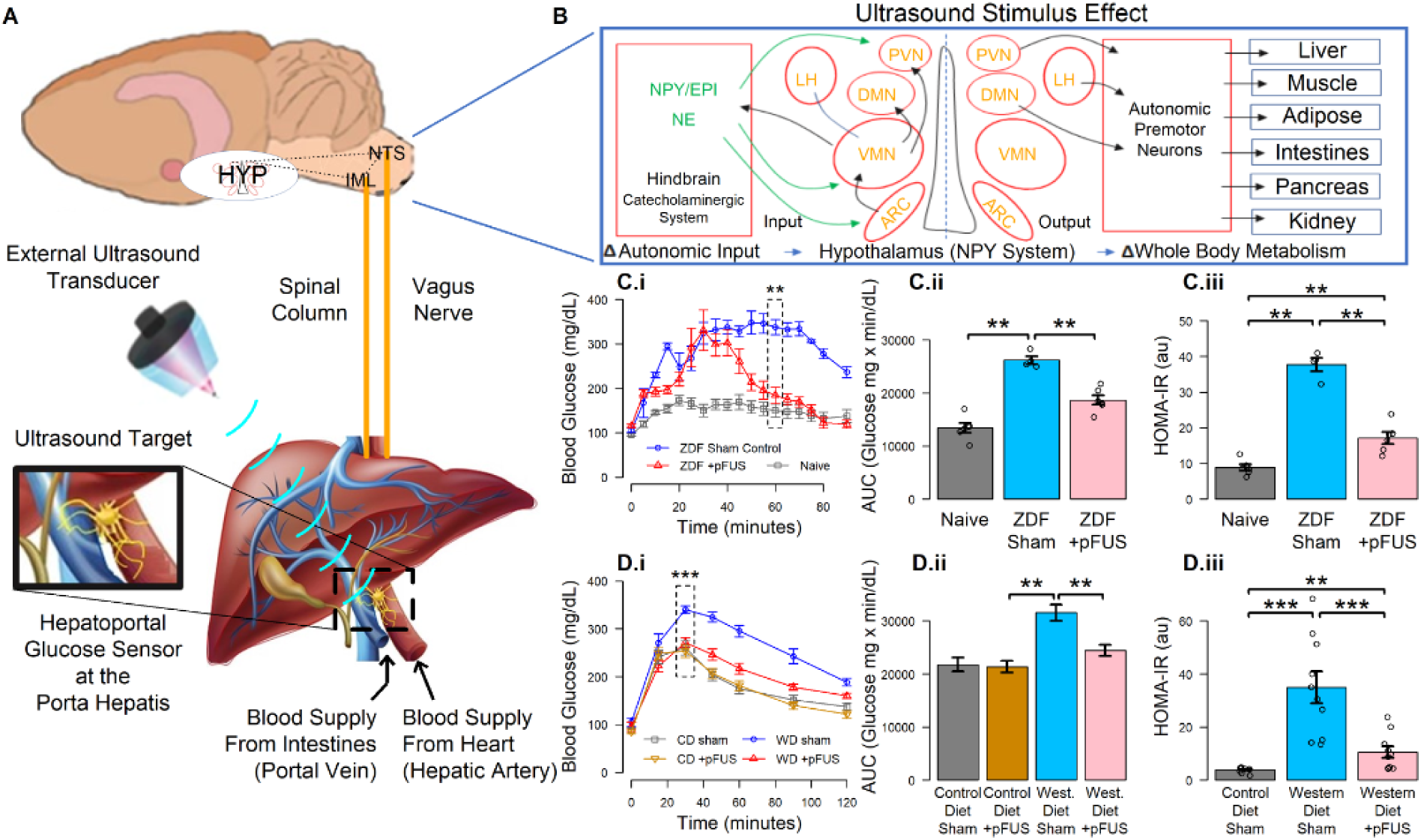
Ultrasound stimulation of the liver-brain neural pathway and effects on glucose metabolism. **A.** A schematic of the ultrasound stimulus target at the porta hepatis, known to contain afferent sensory neurons associated with the hepatoportal glucose sensing system.^1–12,16–22^ These neurons are known to communicate through IML (intermediolateral nucleus) and NTS (nucleus tractus solitaris), modulate specific hypothalamic (HYP) metabolic nuclei, and alter out-going autonomic signaling important in maintaining energy homeostasis. B. Previous studies utilizing invasive hepatoportal glucose clamps have provided evidence that communication between neurons in the porta hepatis and the hypothalamus is mediated by the hindbrain catecholaminergic system^19,43–45^, and affects the output of hypothalamic nuclei via the NPY system.^43,46–48^ **C and D.** Using non-invasive ultrasound for peripheral nerve modulation^1^ enabled activation of these pathways in the genetic (ZDF) and western diet-fed (WD) models of type 2 diabetes (T2D). **C.i.** A single administration of hepatic pFUS improved glucose tolerance in the genetic obese, diabetic ZDF rat (60 min. ** p<0.01, nonparametric Wilcoxon rank-sum test, n=6 per group). **C.ii**. Calculation of incremental areas under the curve (AUC) showed a significant reduction in glucose as compared to Sham stimulated controls (** p<0.01, nonparametric Wilcoxon rank-sum test). **C.iii**. pFUS also showed a significant improvement in insulin sensitivity after a single treatment in the ZDF rat model of T2D (** p<0.01, nonparametric Wilcoxon rank-sum test) as determine by HOMA-IR scoring. **D.** Daily hepatic pFUS improved glucose tolerance and attenuated insulin resistance in WD mice. **D.i**. Hepatic pFUS significantly reduced peak glucose response in obese mice (WD-pFUS) compared to sham stimulated mice (WD-sham; 30 min, *** p < 0.001, nonparametric Wilcoxon rank-sum test, n=10 per group). **D.ii.** A significant reduction in area under the curve was observed in WD-pFUS mice as compared to WD-sham mice (** p < 0.01, nonparametric Wilcoxon rank-sum test). **D.iii.** Hepatic focused ultrasound improves insulin resistance, determined by HOMA-IR. A significant reduction in HOMA-IR score was observed in WD-pFUS group as compared to the WD-sham group (*** p < 0.001, ** p<0.01; nonparametric Wilcoxon rank-sum test).

## pFUS Effect Requires Mechanosensitive Ion Channels in both *In Vitro* and *In Vivo* Models

Hepatoportal glucose sensing is a well-documented phenomenon in which nerve signals report portal vein-hepatic artery glucose gradients during feeding or fasting, that results in neuronal modulation of the metabolic system.^2–4,6–11,16–22^ The sensor comprises at least two glucose sensing mechanisms that utilize GLUT2^9,12,23^ (for low affinity glucose binding) and SGLT3^7,10,11^ (for high affinity glucose binding) and is, therefore, capable of sensing glucose gradients during periods of high or low portal glucose concentration (Supplemental Fig. S1). The neuro-metabolic pathways that are modulated by the sensor are dependent on which of these two molecular sensors are activated^7,10–12,23,24^ and convey signals through multiple vagal and spinal (dorsal root) afferents.^7,11,17,18,25–28^ We have previously shown that low intensity mechanical ultrasound stimuli are capable of site-specific nerve activation in both *in vivo*^1,13–15^ and *in vitro*^14^ models. Herein, the mechanical origin of ultrasound neuromodulation was confirmed by using a purely mechanical stimulus (i.e., replacing the ultrasound transducer with a mechanical piston-based stimulator; Supplemental Fig. S6), and demonstrating the dependence of ultrasound nerve activation on specific families of mechanically sensitive ion channels^29,30^ *in vitro* (Fig. 2 A - C, and Supplemental Fig. S7). Three-dimensional *in vitro* cultures of dorsal root ganglia sensory neurons were activated (as measured by calcium indicator dye) using ultrasound pulse parameters and pressures that were consistent with *in vivo* experiments.^14,31^ Blocking of N-type calcium channels (ω-conotoxin) or voltage-gated sodium channels (tetrodotoxin) did not attenuate the response to pFUS (Fig. 2 D). In contrast, blockade of mechanical/noxious chemical-sensitive piezo (i.e., GxMTx4) and transient receptor potential (TRP; i.e., HC-030031) families inhibited the pFUS effect (Fig. 2 D). Additionally, the glucose lowering effect of hepatic pFUS *in vivo* was abolished by hepatic injection of amiloride, an inhibitor of mechanosensitive cation channels, the degenerin/epithelial sodium channels (DEG/ENAC) and a TRPA1 blocker^32–35^ (Fig. 2 E and F). Several of the ion channels within these families are expressed on afferent neurons, and are required for functional glucose/metabolite sensing.^36–40^ It is known that activation of TRPA1 (target of HC-030031) by allyl isothiocyanate (AITC) improves glucose uptake and insulin signaling in multiple models of T2D^41^, while ablation of TRPV1 and TRPA1 expression (targets of GxMTx4) results in severe insulin or leptin resistance.^38,42^

**Figure 2.**
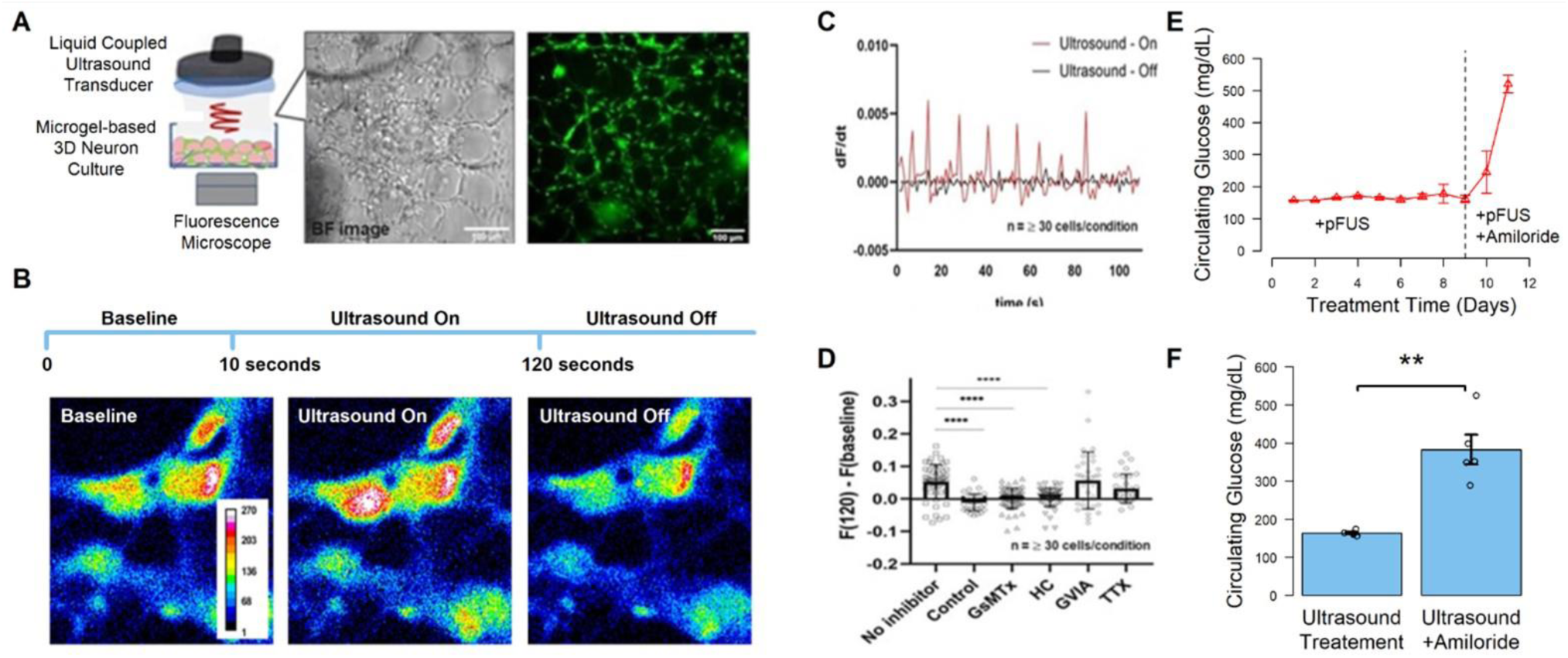
Effects of pFUS on peripheral neurons *in vitro* and *in vivo* with and without specific ion channel blockers. **A.** Schematic of the 3D *in vitro* peripheral neuron culture system, and experimental setup (details provided in methods section) used to capture both bright field and fluorescence images of DRG neuron cells before and after pFUS stimulation. The diameter of hydrogel particles ∼100 μm used for DRG neuron culture has previously been shown^14,29^ to create active axonal networks through the pores formed between hydrogel particles. **B.** The zoom-in fluorescence images show a time lapse of calcium (Ca^2+^) imaging during pFUS stimulation. pFUS stimulation is on at 10 s from the starting point of observation and turned off at 120 s. Ultrasound was then turned off for 2 min (allowing post-ultrasound or off images to be taken) prior to re-starting the stimulus. Calcium concentrations of DRG neuron cells were increased after ultrasound stimulation, and upon pFUS cessation Ca^2+^ concentration in cells returned to baseline levels. **C**. pFUS excitation of DRG neurons led to large changes in Ca^2+^ dependent fluorescence (F). The rate of change, dF/dt, remains small without application of ultrasound stimulus (at 0.83 MPa peak-positive pressure). This is compared to the significant dF/dt increases shown during pFUS stimulation without any ion channel blocker added to the culture (N > 30 cells / each condition). **D.** Observed fluorescence change before/after ultrasound stimulation using multiple ion channel blockers (N > 30 cells / each condition). TTX is a highly specific inhibitor of voltage-gated sodium channels involved in action potential propagation, ω-conotoxin-GVIA is an N-type Ca^2+^ channel inhibitor, and HC030031 is highly sensitive TRP channel inhibitor. When piezo- or TRP-family blockers GsMTx and HC030031 were added to the culture, pFUS-induced activity was significantly suppressed (Dunnet’s multiple test comparison; **** p value < 0.001). **E.** Amiloride (a mechanosensitive cation-selective ion channel, ENAC, and TRPA1 blocker^32–35)^ applied through hepatic injection abolished the effect of hepatic pFUS in ZDF rats (n=5 per group). pFUS was delivered daily for 12 days (day 0 – 11) in the ZDF rats starting on day 60. For the first 10 days (day 0-9), pFUS was applied in the absence of amiloride and resulted in maintenance of an average fasting glucose concentration of 163.86 mg/dL. Injection of a single dose of amiloride on day 10 resulted in immediate increase in circulating glucose, while a second injection on day 11 completely abolished the pFUS effect (i.e. circulating glucose in excess of 500 mg/dL)**. F.** Average circulating fasted glucose in ZDF rats undergoing daily hepatic pFUS before (US Treatment) and after (US + Amiloride) blocking of mechanosensitive ion channels at the porta hepatis using hepatic injections of amiloride (*** p < 0.001; nonparametric Wilcoxon rank-sum test).

## Hepatic pFUS Modulates Metabolism via Afferent Liver-Brain Nerve Pathways

Next, we investigated specific nerve pathways that were activated by hepatoportal pFUS *in vivo* through electrical nerve recordings, hypothalamic neurotransmitter and neuropeptide measurements (with or without site specific chemical nerve lesioning), and expression of the activity-dependent marker cfos (Fig. 3). Hepatoportal pFUS modulated glucose-sensing, but not glucose-insensitive neurons in the paraventricular nucleus (PVN; Fig. 3A and B; and Supplemental Fig. S8 A and B). To study this effect, an intraperitoneal (IP) injection of glucose (20% glucose via inserted catheter) was administered prior to ultrasound, which raised plasma glucose in both stimulated (+pFUS) and unstimulated (-pFUS; Fig. 3C) animals. In the absence of pFUS, the peripheral glucose rise resulted in decreased firing rates of glucose-inhibited (GI) and increased activity of glucose-excited (GE) neurons (Fig. 3 C ii. and iii.). This agrees with previous reports demonstrating experimental activation of these glucosensing neural pathways using a peripheral (i.e., not a portal) glucose source.^49–51^ These pathways have been demonstrated to project to hypothalamic Pomc- and Npy-expressing nerve populations, in a manner that is essential to achieving and maintaining metabolic homeostasis.^47,48,52–58^ Under the same glucose injection conditions, pFUS at the porta hepatis site resulted in attenuation of the glucose-induced firing increase in GE, but not GI neurons (Figure 3 B and C). Glucose-excited pathways have previously been shown to be critical for the initiation and maintenance of signals of energy sufficiency,^59^ with increased activity and firing rates resulting in increased energy expenditure^60–62^ and decreased food seeking behaviors.^60^ In addition, GE neurons have been found to be hyper-responsive in the db/db leptin receptor defect and T2D model.^63–67^ In agreement with these previous findings, the circulating glucose concentrations of the pFUS stimulated group were reduced compared to the unstimulated controls (Fig. 3 C. i.). This suggests that the hepatoportal pFUS stimulus is capable of modulating the ratio of GE-versus GI-neuronal activation after a glucose injection; potentially shifting the hypothalamic response towards a fed-state condition despite high concentrations of peripheral glucose.^5,64,68,69^

**Figure 3.**
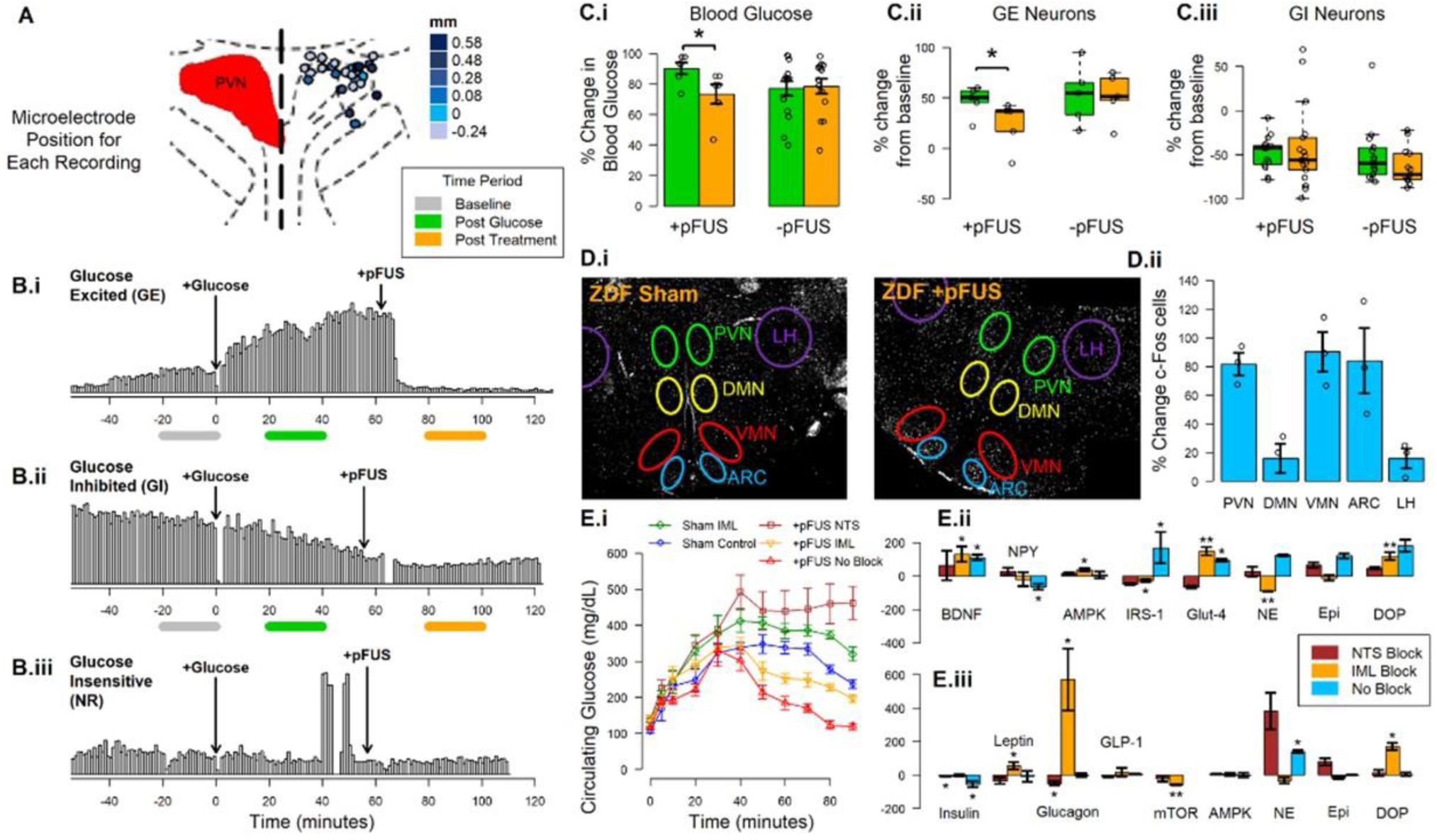
Effects of porta hepatis pFUS stimulation on hypothalamic nerve pathways associated with metabolic control and energy homeostasis. **A.** Schematic position of electrode tips inserted into the paraventricular nucleus (PVN) to measure single neuron firing rates in response to glucose injections versus pFUS stimulation. **B.** Example firing rates from glucose excited (B i.; GE), glucose inhibited (B ii.; GI), or glucose insensitive (B iii.; NR) neurons before glucose injection (baseline), after glucose injection, and after pFUS. **C. i.** Final blood glucose measurements were increased compared to baseline in both pFUS stimulated (+pFUS) and non-stimulated (-pFUS) animals. However, pFUS was associated with a significant decrease in circulating blood glucose compared to -pFUS (*p<0.1, nonparametric Wilcoxon rank-sum test). **C. ii.** GE neurons showed elevated firing rates post-glucose injection compared to baseline, and pFUS resulted in a significant change in firing rate in these neurons (+pFUS; n=5, (*p<0.1, nonparametric Wilcoxon rank-sum test). However, in the absence of pFUS GE neurons maintained an elevated firing rate compared to baseline throughout the experiment (-pFUS; n = 5). **C. iii.** GI neurons showed a significant change from baseline levels post-glucose injection. However, pFUS did not change GI firing rates (n = 16 per condition). **D.** Histochemical analysis of hypothalamic neural pathways associated with response to pFUS after 20 days of daily stimulation in the ZDF model. **D.i.** cFOS immunohistochemistry images show the number of activated neurons in unstimulated (left) versus pFUS stimulated (right) animals. Images were segmented on the paraventricular nucleus (PVN; green), dorsal medial nucleus (DMN; yellow), ventromedial nucleus (VMN; red), arcuate nucleus (ARC; blue), and lateral hypothalamus (LH; purple) Scale bar = 200 microns**. D.ii.** Data showing the percent change in the number of cFos expressing cells with pFUS (n = 6) compared to sham controls (n = 6) in each segmented hypothalamic region (PVN, DMN, VMN, ARC, LH), images in D.i. represent one set of sham versus stimulated paired animals (*p<0.1, nonparametric Wilcoxon rank-sum test). **E.i.** Microinjection of 0.5 microliters of 2% lidocaine hydrochloride (20mg/kg) into the nucleus of the solitary tract (NTS) abolished the effects of pFUS stimulus on circulating blood glucose (dark red circles), as compared to pFUS alone (light red circles). In contrast, microinjection of 0.5 microliters of 2% lidocaine hydrochloride (20mg/kg) into the intermediolateral (IML) nucleus attenuated, but did not completely block, the effects of pFUS (IML-Block+pFUS=yellow circles). Microinjection of lidocaine hydrochloride into the IML in the absence of pFUS (green circles) did not yield any significant change in circulating glucose as compared to standard ZDF Sham pFUS animals (dark blue circles). **E.ii.** Administration of 20mg/kg lidocaine hydrochloride into the NTS abolished the effects pFUS on hypothalamic NPY, IRS, GLUT-4 and NE; however, a significant attenuation of the pFUS effect was only observed with NTS inhibition on Epi and DOP. No significant effect was seen with NTS inhibition on BDNF and AMPK (red bar). In contrast, administration of 20 mg/kg lidocaine hydrochloride in the IML abolished the effects of pFUS on NE and Epi, but attenuated the effects of pFUS on hypothalamic NPY, IRS and DOP. Interestingly, an augmented effect on AMPK and GLUT4 were observed with pFUS stimulus following IML inhibition. No significant effect was seen with IML inhibition on BDNF (yellow bar). **E.iii**. Administration of 20mg/kg lidocaine hydrochloride into the NTS produced a significant increase in circulating NE. In contrast, administration of 20 mg/kg lidocaine hydrochloride into the IML had no effect on NE but produced a significant increase in glucagon release. NTS=nucleus tractus solitarius, NPY=neuropeptide Y, IRS=insulin receptor subunit, GLUT-4=glucose transporter 4, NE=norepinephrine, Epi=epinephrine, DOP=dopamine, BDNF=brain derived neurotrophic factor, AMPK=adenosine 5′ monophosphate-activated protein kinase (*p<0.1, **p<0.001, nonparametric Wilcoxon rank-sum test).

Hepatoportal sensor signaling through the CNS^43^ (via the intermediolateral nucleus (IML) and nucleus tractus solitaris (NTS)) is known to alter NPY/AGRP neuron activity within the arcuate nucleus (ARC)^20^, resulting in reduced inhibition on anorexigenic/catabolic pre-sympathetic neurons in neighboring hypothalamic nuclei.^60,70–72^ These hypothalamic nuclei (including the paraventricular (PVN), ventromedial (VMN), dorsomedial (DMN) nucleus) contain pre-autonomic projections associated with the coordination of whole-body metabolism.^2,5,60,73,74^ In ZDF rodents, pFUS stimulation at the porta hepatis was found to alter cFOS expression across these nuclei (Fig. 3 D i. and ii., and Supplemental Fig. S8), including an increase in cFOS+ neurons within the lateral ARC (i.e., the site of NPY-inhibited glucose excited neurons^75^), and in sites of known anorexigenic/catabolic ARC projections to the VMN and PVN.^75–78^ Chemical blocking at the level of the IML and NTS further validated pFUS modulation of these pathways previously associated with hepatoportal sensor activation.^60,79^ Injection of lidocaine at either site attenuated the glucose lowering pFUS effect in the ZDF model (Fig. 3 E. i.). Blocking of both vagal and spinal mediated pathways at the NTS completely abolished the ultrasound effect on plasma glucose, while blocking spinal (but not vagal) pathways within the IML resulted in partial attenuation (Fig. 3 E.i.). These results again support previous observations demonstrating that hepatoportal sensor signaling occurs through both vagus and IML mediated pathways.^11,17,18,20,23,26^ Our results further demonstrate that in the absence of a chemical block a single pFUS treatment resulted in a significant increase in hypothalamic catecholamine concentrations, reduction in NPY, and increase in IRS-1 and GLUT4 activity (Fig. 3. E ii.). Both IML and NTS block attenuated the catecholamine and NPY response to pFUS. However, the NTS block led to a complete inhibition of the NPY effect, while IML block led to partial attenuation. There was also a differential effect of IML versus NTS block across catecholamines with complete inhibition of the pFUS effect on norepinephrine and epinephrine after IML block, but only partial attenuation of the epinephrine effect after NTS block. The NTS block completely inhibited the pFUS effect on hypothalamic IRS-1 and Glut4 activity, while the IML block had no effect on pFUS-induced GLUT4 modulation. As shown previously, pFUS had no effect on AMPK activity (i.e., a marker of glucose inhibited neuron activity^80^); however, blocking of either the IML or NTS resulted in increased AMPK activity. This supports the previous hypothalamic nerve recording data (Fig. 3 A-C), which shows signaling along the liver-brain pathway through glucose-excited nerves and suggests that activity along this pathway is a key contributor to counter-regulation of glucose-inhibited hypothalamic responses.^58,60,75,81,82^ In blood, the largest effect of the NTS block on pFUS response was on circulating norepinephrine, while IML block resulted in significant changes in circulating glucagon (Fig. 3. E ii.). These results again agree with previous work that utilized portal glucose infusion to activate hepatoportal sensor pathways, which showed that spinal-mediated hepatoportal pathways are essential in modulation of GIP secretion and glucagon output,^11,83^ while vagal-mediated pathways are essential in modulating hepatic glucose-6-phosphatase and glucokinase expression via efferent sympathetic nerve activation^2,17^.

Neurotransmitter/neuropeptide profiling of hypothalamic tissue samples in the ZDF model following 20 days of treatment further validated pFUS activation of hepatoportal sensor associated pathways (Fig. 4A; and Supplemental Figure 9).^19,20^ In the treated animals, pFUS increased hypothalamic concentrations of CNS neurotransmitter NE by 122 %, while NPY concentrations were reduced by nearly 300% compared to sham controls (Fig. 4 A). Alterations in other neurotransmitters with known regulatory or co-modulatory connections to NPY-expressing neurons were also consistent with pFUS induced inhibition of the NPY system. These included increased GABA,^84–86^ decreased glutamate,^87,88^ and increased BDNF^89,90^ concentrations. pFUS also modulated K_ATP_ activity but not AMPK, providing further evidence that these effects are primarily mediated through glucose-excited as opposed to glucose-inhibited hypothalamic pathways.^66,75^ Additional measures of important molecular components of the nutrient sensing pathway (i.e. insulin receptor substrate 1 (IRS1), glucose transporter GLUT4, phosphorylated protein kinase B (pAKT), and glucose-6-phosphate) suggest that these changes are in part due to enhancement of hypothalamic nutrient sensing and insulin signaling.^51,66,67^

**Figure 4.**
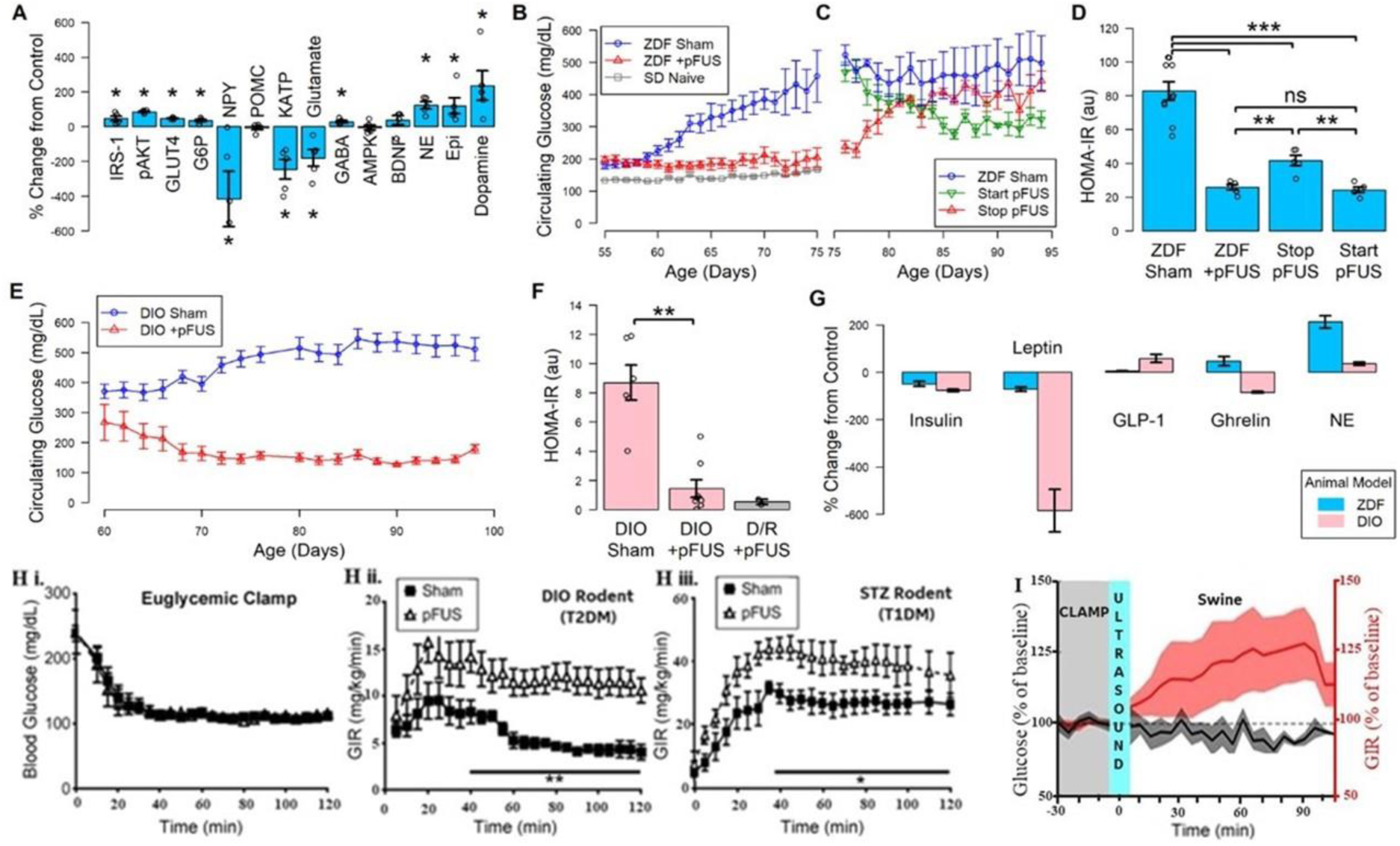
Effects of daily pFUS stimulation in multiple animal models of type II diabetes and species. **A.** The percent change in the concentration of neurotransmitters (NE, norepinephrine; EPI, epinephrine; NPY, neural peptide Y; GABA, gamma-aminobutyric acid; glutamate; BDNF; brain derived neurotrophic factor; and POMC, pro-opiomelanocortin), activity of glucose-sensing associated channels and enzymes (K_ATP_, phosphorylated ATP-sensitive potassium channel; AMPK; phosphorylated AMP-activated kinase), and activity or concentration of insulin signaling associated enzymes and metabolites (phosphorylated IRS-1, insulin receptor substrate-1; pAkt, phosphorylated protein kinase B; GLUT4, phosphorylated glucose transporter 4; and G6P; glucose 6-phosphate) between the stimulated (n=12) and sham control (n=12) ZDF cohorts after 20 days of treatment (i.e. day 55 to 75 of ZDF cohorts; *p<0.1, nonparametric Wilcoxon rank-sum test). **B.** Daily circulating glucose measurements from ZDF animals after daily pFUS (red; n = 12; ZDF +pFUS) or sham control (blue; n = 12; ZDF -pFUS) treatment versus naïve (Sprague Dawley (SD)) controls (gray; n = 8) over twenty days (i.e. day 55 to 75 of ZDF cohorts). **C.** Daily circulating glucose measurements in a cross-over study in which pFUS treatment was stopped at day 75 in the previously stimulated cohort (red; n = 6; Stop pFUS), pFUS treatment was initiated at day 75 in half of the previous sham cohort (green; n = 6; Start pFUS), or sham control treatment was maintained (blue; n = 6; ZDF Sham) for an additional 20 twenty days (i.e. day 76 to 95 of ZDF cohorts). Terminal samples were harvested from 6 animals from the original pFUS treated cohort at day 75 for analysis (ZDF +pFUS). **D.** Terminal (day 95) HOMA-IR scores for the ZDF Sham, ZDF +pFUS, Start pFUS, and Stop pFUS cohorts (**p<0.01,***p<0.001, nonparametric Wilcoxon rank-sum test). **E.** Daily blood glucose measurements in DIO animals undergoing pFUS (DIO +pFUS, red) or sham control (DIO Sham, blue) treatment. **F.** Terminal (day 96) HOMA-IR scores for the DIO Sham, DIO +pFUS, diet resistant (D/R) control +pFUS cohorts (**p<0.01, nonparametric Wilcoxon rank-sum test). **G.** The percent change in the concentration of circulating markers (insulin, leptin, GLP-1, ghrelin and norepinephrine) between stimulated (n=12) and sham control (n=12) ZDF rats and stimulated (n=5) and sham control (n=5) DIO rats (**p<0.01, nonparametric Wilcoxon rank-sum test). **H.i.** Plasma glucose levels during hyperinsulinemic-euglycemic clamp (HEC) in pFUS and sham treated STZ-induced diabetic DIO rats and the corresponding glucose infusion rate (GIR; 2-way ANOVA, n=7 per group, p = 0.0019) **(H.ii)** as well as GIR of during HEC in lean STZ-diabetic animals (**H.iii,** 2-way ANOVA, n=6 per group, p = 0.013). **I**. Time-course of acute effects of pFUS treatment during HEC in healthy swine on plasma glucose (black trace) and GIR (red trace) (mean +/−SEM; relative changes from pre-FUS baseline (n=5 pFUS experiments in 2 animals).

## pFUS Restores Glucose Homeostasis in Multiple Rodent Diabetes Models

The NPY system, a potent orexigenic pathway, interacts with the leptin and insulin systems on several levels.^54,82,84,89^ While recent evidence suggests an important role of this system in human diabetes^82,91,92^, the redundancy and pleotropic effects of central versus peripheral NPY signaling makes it difficult to study or target pharmacologically.^48,54,93^ We tested the non-invasive pFUS inhibition of the hypothalamic NPY system as a therapy in the ZDF and DIO (diet-induced obesity) models of T2D. Daily pFUS for 3 minutes prevented the onset of hyperglycemia compared to sham treated controls and maintained circulating glucose at levels comparable to non-diabetic animals in the ZDF rats (Fig. 4B, n=12; Sprague Dawley (SD) Naive). Following 20 days of treatment, a sub-set of treated and sham cohorts were interchanged (Fig. 4C; crossover experiment). pFUS decreased circulating glucose in the hyperglycemic animals (previously sham controls), while cessation of the stimulus in treated animals resulted in onset of hyperglycemia within three days. No adverse pFUS-induced deviations of serum biomarkers were observed, rather circulating levels of glucose, T4, cholesterol, and amylase all stayed in the normal range compared to sham controls (Supplemental Fig. S9 D). In addition, daily stimulation in a lean Zucker (Fa/fa) control cohort did not result in glycemic deviation or evidence of hypoglycemia in non-diabetic controls (Supplemental Fig. S9 C).

## Hepatic pFUS Modulates Glucose Utilization and Insulin Sensitivity in Multiple Species

Insulin measurements and HOMA-IR scores demonstrated significant improvement in insulin sensitivity in all treated animals (Fig. 4D and Supplemental Fig. S9). Similar results were observed in the DIO model (Fig. 4 E and F) where treatment of hyperglycemic animals reduced circulating blood glucose within days of treatment. This glycemic improvement was significant (393 mg/mL reduction at study termination), sustained for 5 weeks, coincident with reduced circulating insulin, leptin and ghrelin, and improved HOMA-IR scores indicating attenuation of insulin resistance (Figure 4F and G). To specifically quantify the effect of pFUS on glucose utilization we went on to perform hyperinsulinemic-euglycemic clamps in both DIO-STZ diabetic and lean STZ-diabetic SD-rats to test its effectiveness in models of insulin resistant (T2D) and lean insulinopenic (T1D) diabetes, respectively (Figure 4 H. i-iii; Supplemental Figures 10 and 11). Again, our study confirmed that three days of pFUS prior to the clamp resulted in a significant increase in glucose utilization as measured by increased GIR consistent with the restoration of a more insulin sensitive phenotype. Analysis of catecholamine, corticosterone and glucagon responses during the clamp revealed significantly lower glucagon concentrations in both pFUS stimulated models (Supplemental Figures 10 and 11). This supports the notion that reversal of hyperglucagonemia, an important contributor to abnormal glucose homeostasis in diabetes, is one of the mechanisms of pFUS-mediated restoration of glucose.^11,94,95^ It is notable that in the lean STZ-SD cohort of longstanding and profoundly hyperglycemic animals no sustained reduction in hyperglycemia was achieved as we previously observed after a few days of treatment in the ZDF and DIO cohorts (data not shown). These results suggest that, while capable of activating both insulin-dependent and insulin-independent pathways, sustained pFUS reduction in blood glucose might be dependent on an intact insulin signal. This agrees with reports demonstrating that the sustained effects of pharmacological diabetes therapies targeting hypothalamic pathways are dependent on an intact insulin signal.^96,97^

We next investigated whether acute pFUS-induced hepatoportal modulation of metabolism is achievable in a large pre-clinical animal model (Figure 4 I; Supplemental Figures 12). For this purpose, we performed pFUS in non-diabetic swine during hyperinsulinemic-euglycemic clamp conditions. Stimulation was initiated upon achieving a clamped steady state (110 mg/mL glucose) using a 0.5 mU/kg/min continuous insulin infusion (Figure 4 I. and Supplemental Fig. S12). Consistent with our findings in mice and rats, the ultrasound stimulation resulted in an immediate increase of glucose utilization, as measured by the change in glucose infusion rate (GIR), an encouraging step towards further translation of pFUS therapy to humans.

## MultiOmic Profiling Reveals pFUS-induced Metabolic Coordination across Organ Systems

We next used multiomic profiling to establish an atlas of the effects of hepatoportal pathway pFUS stimulation across different metabolic tissues, including liver, intestines, kidney, adipose, muscle, pancreas, hypothalamus and plasma (Fig. 5 and 6; Supplemental Fig. S13-S20). Tissue samples were processed and analyzed by untargeted metabolomics^98^ (plasma and liver) and gene expression profiling^99–104^ (full transcriptome RNA sequencing of all tissues) after 3 days, 4 weeks, and 7 weeks of 3 minute per day pFUS treatment (Supplemental Fig. S13). After the full 7 weeks of pFUS, significant alteration in gene expression was observed in all measured tissues as compared to sham controls (Fig. 5 A and B), with 27.5 % (ZDF model; p-value = 4.1e-4) and 28.8% (DIO; p-value = 1e-10) of these previously identified as diabetes risk genes. The top twenty modified diabetes-related genes show significant differences in pFUS effect within the DIO and ZDF models (Fig. 5 C and D). Genes most broadly affected across the tissues in the Zucker model (Fig. 5C) included the gene encoding bmal1 (arntl; endogenous circadian clock gene associated with suppression of diurnal variation in circulating glucose^105^) and insulin receptor substrate 2 (IRS2). Leptin receptors are present on circadian clock neurons, and deficiencies in leptin signaling (such as those in the ZDF model) are known to effect central clock function.^106–108^ Furthermore, dysregulation of peripheral circadian transcription factor bmal1 is associated with altered expression of carbohydrate regulating proteins (including IRS2^109^) and significant metabolic dysfunction. The most broadly pFUS-impacted genes in the DIO model (Fig. 5D) included DNA-dependent protein kinase catalytic subunit (PrkDC). PrkDC is a potent regulator of AMPK activity, including its role in glucose uptake, fat oxidation, energy production, and mitochondrial biogenesis.^110^ Of the top twenty pFUS altered diabetic genes, changes associated with intestinal tissue were more prominent in the ZDF model, while the DIO model was associated with an increased number of effected genes in liver tissue. The ZDF model is characterized by intestinal enlargement (driven by increased food intake), and significantly elevated biomarkers of insulin resistance within intestinal epithelial cells.^111^ In contrast, the DIO model is characterized by significant fat accumulation, which leads to inadequate fatty acid oxidation, and accumulation of free fatty acids in the liver.^112^ The unique effect of pFUS in the DIO versus ZDF models suggest that activation of hepatoportal pathways and modulation of hypothalamic metabolic coordination holds therapeutic potential across multiple risk and pathogenic factors involved with T2D (Supplemental Fig. S1).

**Figure 5.**
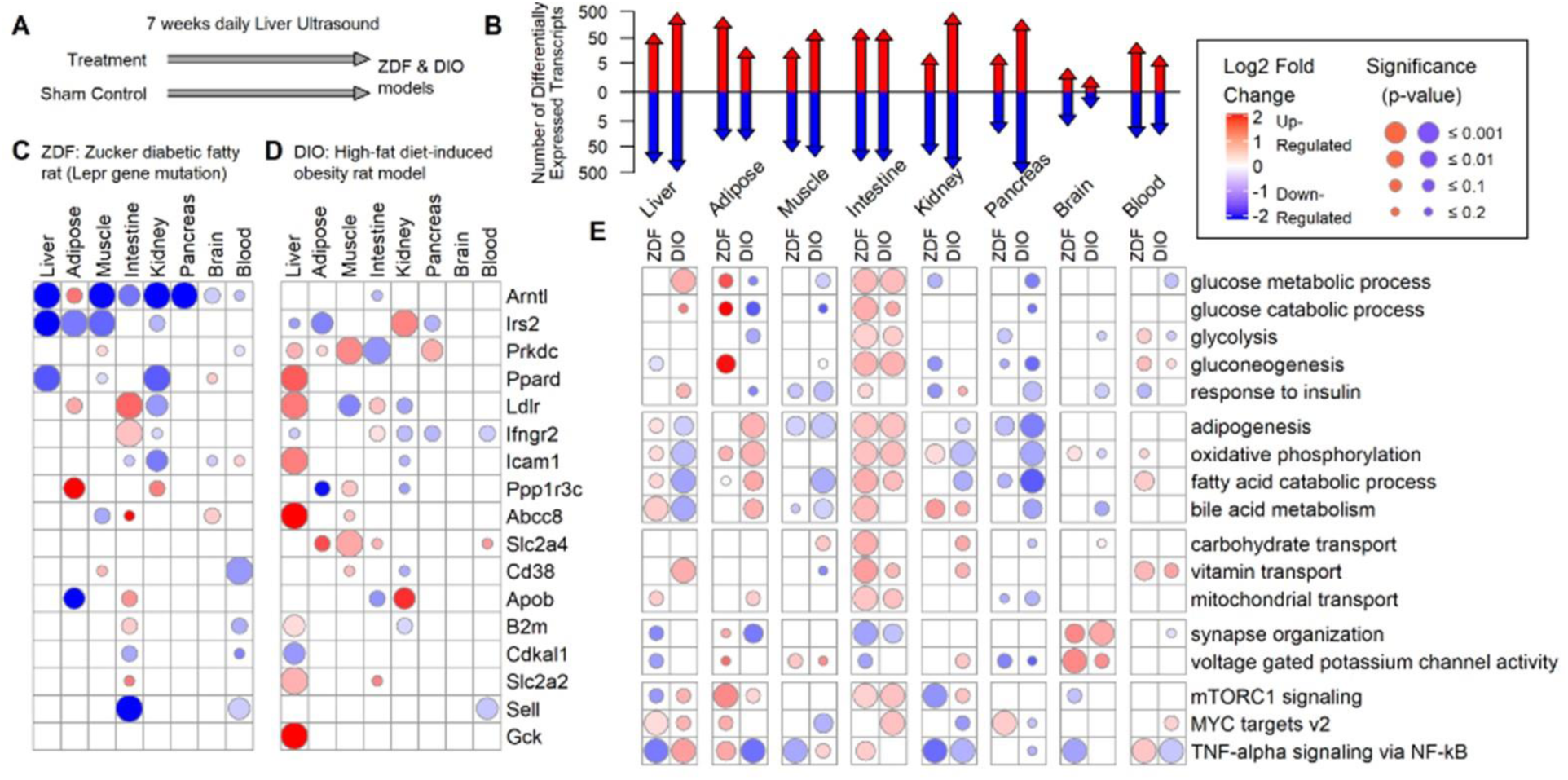
Tissue-specific transcriptomic changes after seven weeks of daily pFUS stimulation of animal models of obese type II diabetes. **A.** Genetic (ZDF) and a DIO animal models of type II diabetes underwent seven weeks of daily pFUS stimulation along with sham treated control animals (n=5 per treatment group). **B.** The number of differential RNA transcripts (adjusted p-value < 0.1) that are upregulated (red arrows) and downregulated (blue arrows) across eight tissues between the pFUS stimulated vs sham controls in both ZDF and DIO animal models. **C-D.** The top diabetic genes that are significantly upregulated (red) or downregulated (blue) across eight tissues in ZDF and DIO animal models respectively. The size of each circle represents the p-value of differentiation between the pFUS stimulated group (n=5) vs sham treated controls (n = 5). **E.** Biological processes found to be significant (Familywise-error rate p-value < 0.1) from gene set enrichment analysis across tissues and the ZDF and DIO animal models. Biological processes are highlighted as being activated (red) or repressed (blue) if the median value of the log2 transformed fold changes for the set of genes was positive or negative respectively. The biological processes includes glucose metabolic process [GO:0006006], glucose catabolic process [GO:0006007], glycolysis [Broad Hallmark], gluconeogenesis [Reactome R-HSA-70263], response to insulin [GO:0032868], adipogenesis [Broad Hallmark], oxidative phosphorylation [Broad Hallmark], fatty acid catabolic process [GO:0009062], bile acid metabolism [Broad Hallmark], carbohydrate transport [GO:0008643], vitamin transport [GO:0051180], mitochondrial transport [GO:0006839], synapse organization [GO:0050808, voltage gated potassium channel activity [GO:0005244], mTORC1 signaling [Broad Hallmark], MYC targets v2 [Broad Hallmark], and TNF-alpha signaling via NF-kB [Broad Hallmark].

**Figure 6.**
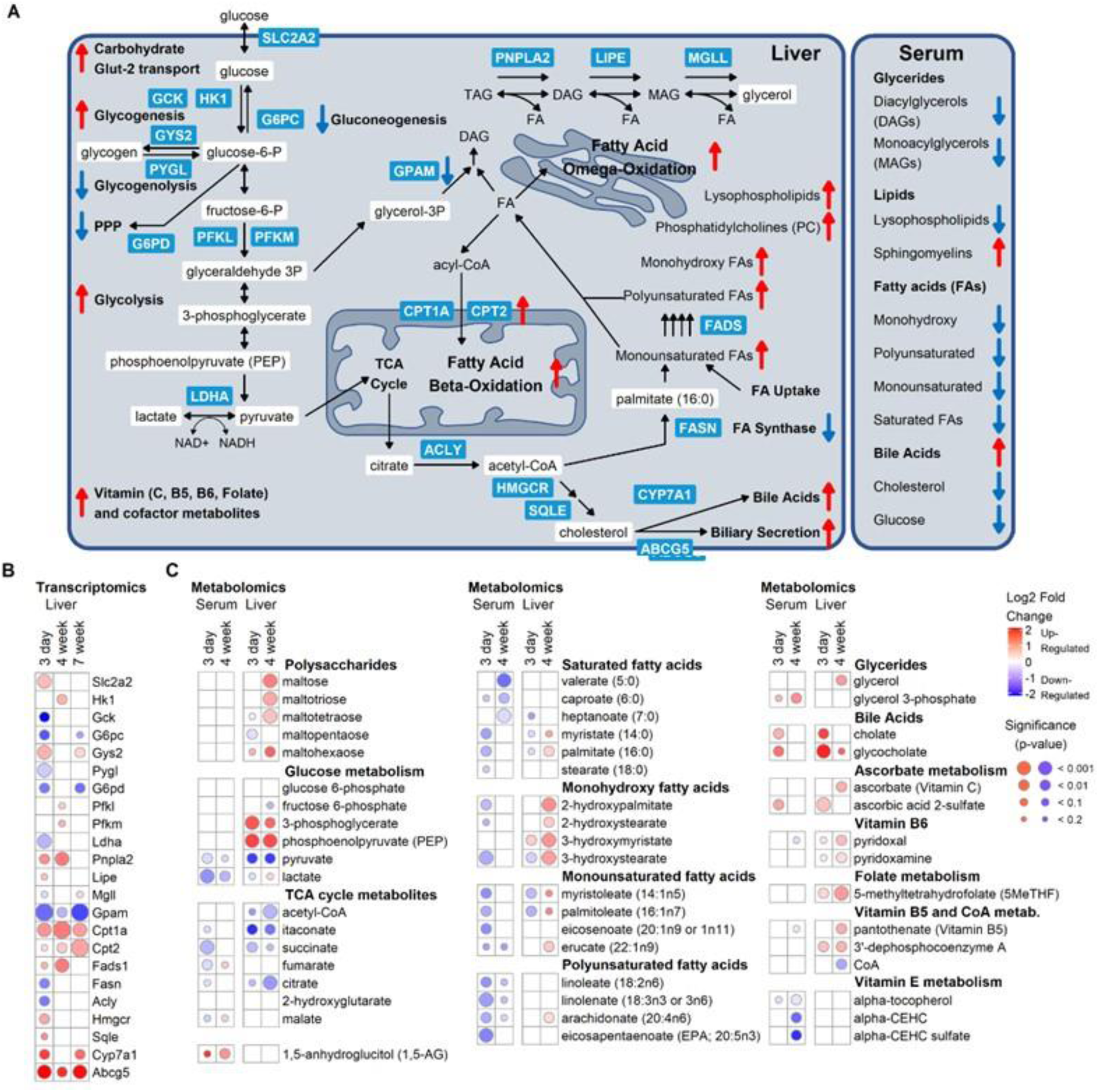
Metabolomic changes in liver and blood serum confirm changes in key metabolic pathways inferred from the liver transcriptomic data. **A.** Schematic of key protein enzymes (blue rectangles), metabolites (white rectangles) and metabolic pathways (e.g., gluconeogenesis, fatty acid and bile acid metabolism, oxidative phosphorylation, mitochondrial transport) that are either upregulated (red upward arrows) or downregulated (blue downward arrows). **B.** Differential RNA transcripts from liver tissue of ZDF animals undergoing 3 days, 4 weeks, or 7 weeks of daily pFUS stimulation vs sham controls. The protein encoded by these differentially expressed genes are presented in the schematic diagram. **C.** The changes in metabolites in blood serum or liver tissue at 3 days and 4 weeks between ultrasound treated vs controls. The size of each circle represents the p-value of differentiation between the pFUS stimulated group (n=5) vs sham controls (n = 5). The color of the circle represents the degree of being upregulated (red) or downregulated (blue). TAG = triacylglycerol; DAG = diacylglycerol; MAG = monoacylglycerol; FA = fatty acid; PPP = Pentose Phosphate Pathway.

Gene sets (Fig. 5E) enriched in the hypothalamic/brain samples by pFUS included those associated with synaptic reorganization and altered voltage-gated potassium channel activity (in agreement with results shown in Fig. 4A). In liver, gene sets enriched by pFUS included those associated with the MYC pathway (known to activate multiple glucose utilization enzymes^113^), and fatty acid/bile acid metabolism. In the intestines, gene expression hallmarks of improved nutrient/vitamin transport and metabolism were apparent; these included significant changes to glycolytic and gluconeogenic gene sets, and markers of improved mitochondrial function. In adipose tissue, markers of suppressed lipolysis were down regulated by pFUS (e.g., mTORC1 signaling^114^), and reduction in adipogenic gene sets were observed in liver, intestinal, pancreatic, and muscle samples.

By day 3, pFUS treated ZDFs exhibited expression signatures of an increased capacity for hepatic glucose uptake (i.e., Slc2a2) and glycogen storage (i.e., Gys2; Pygl) with a decrease in gluconeogenesis (i.e., G6PC; Fig. 6 A and B, Supplemental Fig. S14). These signatures of improved glucose metabolism were accompanied by reduced expression of enzymes that catalyze glycerolipid biosynthesis, and upregulation of enzymes that catalyze fatty acid oxidation. The signatures of improved fatty acid oxidation continued through the 4- and 7-week timepoints after hyperglycemia was already resolved. Metabolomic signatures within the liver and serum validate these transcriptomic observations. Circulating free fatty acids were reduced within 3 days of treatment, and the improvement in dyslipidemia was apparent across fatty acids of all chain lengths and degree of saturation (Fig. 6 A and C; Supplemental Fig. S15 and S16). Within the liver, TCA cycle metabolites were decreased, further supporting renewed capability to uptake and utilize circulating fatty acids. In addition, circulating 1,5-anhydroglucitol concentrations were increased following treatment (a metabolite whose reabsorption in the renal tubules is inhibited during glucosuria^115^), a signature of re-established glycemic control. These circulating biomarkers were also accompanied by increased hepatic concentrations of gluconeogenic precursors (PEP and 3-phoshpcylerate), further validating the decreased expression of gluconeogenic enzymes in the pFUS treated rodents.

By week 4, additional signatures of the pFUS reversal of metabolic dysfunction were apparent including hepatic and circulating signatures of improved bile acid and vitamin metabolism (Fig. 6C and Supplemental Fig. S17).^116^ Additional details on pFUS-induced transcriptomic and metabolic effects on the liver-adipose^117^ and liver-muscle^23^ metabolic axis are available in Supplemental Fig. S14 and S15. Supplemental Fig. S18 – S19 contain further evidence of broad pFUS-induced modulation of whole-body metabolism, including modulation of circulating sphingomyelin and ceramide concentrations (Supplemental Fig. S18; lipids associated with insulin resistance and pancreatic beta cell function^118^) and reduction in circulating markers of diabetes-induced renal impairment^119^ (Supplemental Fig. S19 and S20; phosphatidylcholine and lysophospholipids).

## Conclusions

Our anatomically precise pFUS neuromodulation method^1^ enabled daily activation of the hepatoportal glucose sensing pathway^6,8–11,23,26^ without the need for invasive portal clamps and infusions. The stimulation was shown to activate neuronal liver-brain pathways utilized by the body to enhance insulin sensitivity, glucose uptake, and energy substrate utilization.^4,5,8,26,68,69,74,117^ The pFUS-induced effect was shown to be dependent on vagus nerve and sympathetic spinal afferent-mediated communication to the hypothalamus, ultimately inhibiting the central NPY system. The resulting hypothalamic modulation of autonomic output to multiple organs resulted in sustained glycemic control in multiple animal models of impaired glucose metabolism and diabetes. pFUS activation of these pathways was validated in three different species and is currently undergoing clinical feasibility testing in both diabetic (NCT04502212) and pre-diabetic (NCT04622683) subjects.

In summary, our findings confirm that liver-brain neural pathways play an important role in maintaining whole body glucose homeostasis and that non-invasive portal focused ultrasound represents an exciting new treatment modality to alter whole body glucose metabolism. We speculate that this new tool could be used as a non-pharmaceutical adjunct or even alternative to current treatments of diabetes.

## Acknowledgments

Gjessing Petter Fosse (University Hospital of Northern Norway), for helpful discussions on large animal experiments.

## Funding

Experiments in the manuscript were partially funded with Federal funds from the Defense Advanced Research Project Agency (United States Department of Defense; DAPRA DoD; DARPA HR0011-18-C-0040). R.H. is supported by the NIH via UL1 TR001863; P30 DK045735; R01 DK101984; R01 DK020495; and DARPA 401126008. The views, opinions and/or findings expressed are those of the authors and should not be interpreted as representing the official views or policies of the Department of Defense or the U.S. Government.

## Author contributions

VC performed chronic stimulation experiments in ZDF and DIO models and data analysis, and short term stimulation experiments involving chemical lesioning and in vivo blocking; HM performed the in vitro stimulation experiments and data analysis; ZH, KA, MD, LB, TM performed in vivo electrical recording experiments and contributed to data analysis; KQ, JNT, WS performed swine model experiments and data analysis; TSH, AD, TT performed western diet model experiments, NT, YD, KJC performed rodent H/E clamp experiments; JG performed transcriptomic and metabolomic analyses and data presentation and statistical analysis across manuscript data; RM performed analysis of electrical nerve recording data; KW, TJK, YF installed, set-up, and calibrated ultrasound equipment and contributed experimental results from the mechanical piston stimulation data; EL, CM assisted in sample collection, storage, and analysis of DIO and ZDF biological samples; JA, KJT, TRC, DDC, DS, SZ, SSC, RH, CP designed research and experiments, performed data analysis, edited and co-wrote sections of the manuscript, CP wrote the manuscript including assembly of sections from collaborating institutions.

## Competing interests

VC, JG, RM, KW, EL, CM, YF, TJK, JA, CP are employees of General Electric and declare that GE has filed US and international patent applications describing methods, devices, and systems for precision organ-based ultrasound neuromodulation. HM, ZH, KQ, TSH, NT, YD, KJC, JT, AD, TT, KA, MD, LB, TM, KJT, TRC, DDC, DS, SZ, SSC, RH have received research funding from GE to investigate the effects of ultrasound on metabolism;

## Data and materials availability

accession numbers to any data will be included, if applicable, in published manuscript

## List of Supplementary Materials

Materials and Methods

Figures S1-S20

## Supplementary Materials for

### Materials and Methods

#### Ultrasound stimulation hardware/parameters/measurements

##### 1.1-MHz Single-element Focused Ultrasound System

The 1.1 MHz single-element focused ultrasound system (Fig 2SA) is comprised of a signal generator (Model 33120A, Agilent Technologies Inc., Santa Clara, CA), a RF power amplifier (Model 350L, Electronics & Innovation Ltd., Rochester, NY) and a 1.1 MHz focused single-element ultrasound transducer (Model H102, Sonic Concepts Inc., Bothell, WA). The transducer is connected to the output of the power amplifier using a matching network (Model H102, Sonic Concepts Inc., Bothell, WA). The transducer element is 64 mm in diameter and has a 63.2 mm radius of curvature with a 20 mm diameter hole in the center into which a small imaging transducer can be inserted for image guidance (Fig S2B). The transducer is acoustically coupled to the animal through a 6-cm tall plastic standoff cone filled with degassed water.

The nominal settings of the waveform (Fig S3B) are listed below:

- 1.1 MHz Carrier
- 135 mV pk-to-pk signal generator Amplitude
- 150 usec Pulse Duration
- 5 Hz Pulse Repetition Frequency

The acoustic performance of the system was characterized at an ISO/IEC 17025:2017 accredited laboratory (Acertara Acoustic Labs, Longmont, CO). Under the above settings three separate systems were characterized, producing a mechanical index, MI of 1.79 mean (+/− 0.10 stdev), a derated peak negative pressure, pr.3 of 1.87 (+/− 0.11) MPa, a derated spatial-peak pulse-average intensity, Isppa.3 of 125.7 (+/− 15.0) W/cm2 and a derated spatial-peak temporal-average intensity, Ispta.3 of 94.3 (+/− 11.2) mW/cm2. Good linearity (0.9947 R^2^) of MI was observed over an amplitude range from 100 mV pk-to-pk to 200 mV pk-to-pk therefore acoustic output can reasonably be linearly extrapolated to other amplitudes, pulse durations and pulse repetition frequencies (Fig S3B). The simulated pressure profile has a full width half maximum amplitude of 1.8 mm laterally and 6 mm in the depth direction using Field II^120,121^ (Fig S3A). The −6dB diameters at the peak were measured to be 1.45 (+/− 0.02) mm in the X scan axis and 1.46 (+/− 0.02) mm in the Y scan axis at the accredited laboratory.

##### Ultrasound System and Parameters used during Electrophysiology and Nerve Recording Experiments

The focused ultrasound system used for recording experiments was comprised of a signal generator (Model 33120A, Agilent Technologies, Santa Clara, CA), a RF power amplifier (Model 350L, Electronics & Innovation Ltd., Rochester, NY) and a custom made 2.5 MHz focused single-element ultrasound transducer. The transducer element is 19 mm in diameter and has a 25.4 mm radius of curvature. The transducer is connected directly to the output of the power amplifier without a matching network. The transducer is acoustically coupled to animal using ultrasound gel without a standoff cone.

The nominal settings of the waveform for this system are listed below:

- 2.5 MHz Carrier
- 300 mV pk-to-pk signal generator Amplitude
- 120 usec Pulse Duration (300 carrier cycles)
- 5 Hz Pulse Repetition Frequency

##### Image Target Identification Methods (Figs S2C, S2D)

The portal vein of the rat liver was identified using an ultrasound imager. Two methods were employed to identify the target and position the focused ultrasound transducer. The first method involved placing an ultrasound transducer directly on the skin, identifying the location of the portal vein and removing the ultrasound imaging probe before placing the focused ultrasound transducer in the same location. In this manner, a high-frequency probe (Model L8-18i linear probe, GE Healthcare, Milwaukee WI) was used to provide high-resolution imagery consisting of B-mode imaging where the target was identified by changes in contrast (Fig S2C left) as well as color Doppler imaging where the target was identified by the pulsation of blood flow in the portal vein (Fig S2C right). The second method involved placing an ultrasound transducer through the opening in the center of the focused ultrasound transducer while stimulation was being applied. In this manner, a lower frequency probe (Model M3S phased-array probe, GE Healthcare, Milwaukee WI) was used to obtain images at depth through the through 6-cm tall plastic standoff cone water coupling cone and target identification was obtained by observing changes in contrast.

### Animal Model of Type 2 Diabetes

#### Zucker diabetic fatty (ZDF) rat model

Adult male ZDF rats (Charles River, Kingston, NY USA) were ordered for arrival prior to 8 weeks of age. All animals were maintained on a high caloric rodent chow (Purina 5008), provided, with water, provided ad libitum. All rats were housed at 25 °C on a 12-h light/dark cycle and acclimatized for 1 week, with handling, before experiments were conducted to minimize potential confounding glucose measures due to stress response. All procedures performed in accordance with the National Institutes of Health (NIH) Guidelines under protocols approved by the Institutional Animal Care and Use Committee (IACUC) of GE Global Research.

ZDF animals were acclimated by daily handling to prevent stress-induced changes in circulating glucose. After 8 weeks, ZDF rodents begin to exhibit rapid development of the diabetic phenotype and are then separated into either Sham-CTRL or pFUS treatment groups for acute (Figure 1C. Figure 2E and F) or chronic (Figure 2D, Figure 3A-D, Figure 3G) ultrasound stimulation.

#### Diet Induced Obese rat model

Adult male Sprague-Dawley rats (Charles River, Kingston, NY USA) were ordered to arrive at 8 weeks of age. Upon arrival animals were placed on a high fat diet (45% kcal as fat; Research Diet, D12451; New Brunswick, NJ) and maintained consistently for 8 weeks. Following 8-weeks of a high fat diet, rats were treated with streptozotocin (STZ; 30mg/kg in 0.1M citric acid buffer, pH=4.5 i.p.). Diabetes was confirmed 96 hours following STZ injection by evalulated blood glucose levels using a handheld glucometer Freestyle Freedom Lite (Abbot Diabetes Inc., Alameda, CA, US). Rats having blood glucose level of ≤280 mg/dl (11.1 mM) or greater were considered to be diabetic. Those animals which failed to develop hyperglycemia were labeled as diet resistant (DR) and kept on study as a control.

##### Peripheral focused ultrasound stimulation in ZDF and DIO models

All rats were anesthetized at 1-4% isoflurane at 1 L/min O_2_. Rats were then placed on a water circulating warming pad, with a rectal thermometer probe to maintain body temperature.

The area above the stimulation target was shaved and hair was fully removed with Nair. The porta hepatis was localized using a custom ultrasound imaging device (Vivid E9; GE Healthcare). The location was marked with a permanent marker and a pFUS stimulation probe was placed on the target area (represented in Figure 1A). The device then delivered 3 minutes of stimulation (1.1 MHz, 200mV per pulse, 150 burst cycles, 500 µs burst period). Blood glucose levels of the rats were monitored on a daily basis for chronic studies (Figure 2D, Figure 3A-D and Figure 3F) and at 5 minute intervals for acute and OGTT studies (Figure 1C and Figure 2E-F). In chronic studies (Figure 2D, 3A-D and 3F) additional blood samples were collected on a weekly basis for the analysis of circulating markers. A final terminal blood sample was collected at the time of euthanization and used for the evaluation of circulating insulin levels.

##### Blood glucose determination

Blood samples obtained from the tail vein were used to assess daily glucose (for chronic studies) and 5-minute glucose intervals (for acute studies) using a Freestyle Freedom Lite (Abbott Diabetes Inc., Alameda, CA, US). The Freestyle Freedom Light meter using a small blood volume (0.3uL) and as such no additional fluids were needed to recover total volume following blood sampling.

##### Glucose Tolerance Tests

Rats were fasted overnight (12-16 h) prior to anesthetization and placement of a tail vein catheter and collection of baseline blood samples. Immediately following the collection of the baseline sample, a single round of pFUS stimulation was applied and the animal allowed to recover from anesthesia. Upon recover, the animal was given a single oral dose of 2g/kg glucose solution by oral syringe feeding. Following glucose administration blood samples were collected on a 5-minute basis from the tail vein by catheter collection. All procedures were performed following Institutional Animal Care and Use Committee (IACUC) of GE Global Research. The area under the curve determined by glucose levels at baseline and 120 minutes after glucose overload was considered for calculation of AUC-OGTT.

##### Insulin resistance evaluation

Glucose and insulin levels were utilized to determine insulin resistance by applying the homeostatic model assessment-insulin resistance (HOMA-IR) formula. The HOMA-IR is defined as fasting insulin (microU/L) x fasting glucose (nmol/L)/ 22.5 (REF).

##### Blood Chemistry Analysis

Whole blood samples collected from rats was used to assess the overall health of the rodents using a Abbaxis Vetscan blood analyzer and the associated Comprehensive Metabolic Panel rotors. A detailed protocol for the Comprehensive Metabolic Panel is provided by the respective supplier of the product: https://www.abaxis.com/sites/default/files/resource-packages/VetScan%20Comprehensive%20Diagnostic%20Profile%20PI.pdf

##### Tissue Harvesting for Protein and Metabolomic Analysis

An incision was made starting at the base of the peritoneal cavity extending up and through to the pleural cavity. Tissues collected included the liver, muscle (gastrocnemius), visceral white fat (abdominal), intestine (jejunum), brain (hypothalamus punch), kidney, pancreas, spleen, and blood.

##### Tissue Preparation for ELISA analysis

Organ samples (e.g. liver, muscle, adipose, brain) were rapidly removed and homogenized in a solution of phosphate-buffered saline (PBS), containing phosphatase (0.2-mM phenylmethylsulfonyl fluoride, 5-µg/mL aprotinin, 1-mM benzamidine, 1-mM sodium orthovanadate, and 2-µM cantharidin) and protease (1-µL to 20 mg of tissue as per Roche Diagnostics) inhibitors. A targeted final concentration of 0.2-g tissue per mL PBS solution was applied in all samples. Blood samples were stored with the anti-coagulant disodium (ethylenedinitrilo)tetraacetic acid (EDTA) to prevent coagulation of samples. Samples were then stored at −80 °C until analysis.

##### Tissue Preparation for Transcriptomic analysis

Organ samples (e.g. liver, muscle, adipose, brain) were rapidly removed and cut into sections ≤ 2mm in thickness and transferred into pre-filled tubes containing RNA later (Applied Biosystems; Foster City, CA) and stored in at 4-5°C for 24 hours. Following 24-hour incubations organ samples were transferred to a clean centrifuge tube and stored at −20°C prior to analysis.

##### Tissue Preparation for Metabolomic analysis

Organ samples of approximately 20mg was snap frozen in liquid nitrogen and stored at −80C prior to metabolomic analysis. All samples were prepared at Metabolon using their automated MicroLab STAR® system from Hamilton Company. Briefly, small molecules are extracted with methanol under vigorous shaking for 2 min (Glen Mills GenoGrinder 2000) followed by centrifugation. The extract is divided into five fractions: two for analysis by two separate reverse phase (RP)/UPLC-MS/MS methods using positive ion mode electrospray ionization (ESI), one for analysis by RP/UPLC-MS/MS using negative ion mode ESI, one for analysis by HILIC/UPLC-MS/MS using negative ion mode ESI, and one reserved for backup. Further methods on untargeted metabolomic analysis are found below.

##### HPLC analyses

Serum samples were injected directly into the machine with no pre-treatment. Tissue homogenates were initially homogenized with 0.1-M perchloric acid and centrifuged for 15 min, after which the supernatant was separated, and the sample injected into the HPLC. Catecholamines norepinephrine and epinephrine were analyzed by HPLC with inline ultraviolet detector. The test column used in this analysis was a Supelco Discovery C18 (15-cm × 4.6-mm inside diameter, 5-µm particle size). A biphasic mobile phase comprised of [A] acetonitrile: [B] 50 = mM KH_2_PO_4_, set to pH 3 (with phosphoric acid). The solution was then buffered with 100-mg/L EDTA and 200-mg/L 1-octane-sulfonic acid. Final concentration of mobile phase mixture was set to 5:95, A:B. A flow rate of 1 mL/min was used to improve overall peak resolution while the column was held to a consistent 20 °C to minimize pressure compaction of the column resulting from the viscosity of the utilized mobile phase. The UV detector was maintained at a 254-nm wavelength, which is known to capture the absorption for catecholamines including norepinephrine, epinephrine, and dopamine.

##### Immunohistochemistry and histology protocols

Tissue extraction and paraffin block conversion performed as follows. Put tissue (rat brain) into fixative immediately and fix ∼24 h in 10% formalin at 4 °C. Process tissue with the following protocol (with vacuum and pressure during each incubation): 70% ethanol, 37 °C, 40 min, 80% ethanol, 37 °C, 40 min, 95% ethanol, 37 °C, 40 min, 95% ethanol, 37 °C, 40 min, 100% ethanol, 37 C, 40 min, 100% ethanol, 37 °C, 40 min, xylene, 37 °C, 40 min, xylene, 37 °C, 40 min, paraffin, 65 °C, 40 min, paraffin, 65 °C, 40 min, paraffin, 65 °C, 40 min. Sample is then left in this paraffin until ready for embedding (not to exceed ∼12–18 h).

Embed into Paraffin block for sectioning, allow block to cool/harden before sectioning. Section 5-µm thick, float on 50 °C water bath for collection. Use positively charged slides and try to position the tissue in the same orientation for every slide. Air dry slides. Overnight at room temperature seems to be the best for drying but the slides can be placed on a 40 °C slide warmer to speed up the drying process, but do not leave slides more than an hour on the warmer. Store slides at 4 °C.

Formalin-fixed paraffin-embedded (FFPE) tissue samples (rat brains) were baked at 65 °C for 1 h. Slides were deparaffinized with xylene, rehydrated by decreasing ethanol concentration washes, and then processed for antigen retrieval. A two-step antigen retrieval method was developed specifically for multiplexing with FFPE tissues, which allowed for the use of antibodies with different antigen retrieval conditions to be used together on the same samples. Samples were then incubated in PBS with 0.3% Triton X-100 for 10 min at ambient temperature before blocking against nonspecific binding with 10% (wt/vol) donkey serum and 3% (wt/vol) bovine serum albumin (BSA) in 1× PBS for 45 min at room temperature. Primary antibody c-Fos (Santa Cruz-SC52; sc-166940) was diluted to optimized concentration (5 μg/mL) and applied for 1 h at room temperature in PBS/3% (vol/vol) BSA. Samples were then washed sequentially in PBS, PBS-TritonX-100, and then PBS again for 10 min, each with agitation. In the case of secondary antibody detection, samples were incubated with primary antibody species-specific secondary Donkey IgG conjugated to either Cy3 or Cy5. Slides were then washed as above and stained in DAPI (10 μg/mL) for 5 min, rinsed again in PBS, and then mounted with antifade media for image acquisition. Whole-tissue images were acquired on fluorescence microscope (Olympus IX81) at ×10 magnification. Autofluorescence, which is typical of FFPE tissues, needs to be properly characterized and separated from target fluorophore signals. We used autofluorescence removal processes, wherein an image of the unstained sample is acquired in addition to the stained image. The unstained and stained images are normalized with respect to their exposure times and the dark pixel value (pixel intensity value at zero exposure time). Each normalized autofluorescence image is then subtracted from the corresponding normalized stained image. We ensured that the same region in the stimulated and control samples were imaged.

##### cFos analysis and measures of US-induced activation

pFUS stimulated and sham animals were rapidly euthanized, and brains removed and transferred to 10% paraformaldehyde for 24 h, after which they were transferred to a 30% sucrose solution and stored for 4 °C prior to paraffin embedding (detailed in the IHC section above). Coronal section (5–10 µm) were cut by cryostat. Structures were anatomically defined according to an anatomical atlas. Quantification of c-Fos positive cells was counted with a fixed sample window across at least four sections by an experimenter blinded to the treatment conditions associated with each distinct coronal section. Regions of interest were as follows: paraventricular hypothalamic nucleus, ARC, VMN, DMN, LH, and mammillothalamic tract (all structures visible in coronal slices taken between Bregma −2.56 to −3.60 mm). The number of c-Fos positive cells in each group were expressed as a % of cFos+ cells as compared to Sham-stimulated control littermates.

##### Tissue Glycogen Measurements

Tissue samples were washed thoroughly in PBS and transferred into a vial of potassium hydroxide (1mg:4uL; 30% w/v). The suspended tissue was then heated in a boiling water bath for 10 minutes with constant mixing. The sample was then cooled in an ice bath prior to the addition of ethanol (100%; EtOH) for a final concentration of 55% EtOH (v/v). The mixture was then vortexed followed rapidly by 10 minutes of centrifugation at 1700 x g. The supernatant was then decanted, and the remaining pellet resuspended in 2mL of filtered distilled water. This solution was then used to meaure total glycogen for each tissue type as per supplier (Abcam: https://www.abcam.com/Glycogen-Assay-Kit-ab65620.html).

##### Chemical Lesioning at the NTS and IML

For NTS injection, rats were anesthetized with 1-2% isoflurane and then placed into a stereotaxic instrument with the head angled down at approximately 45°. A water circulating heating pad placed immediately below the animal to maintain core body temperature throughout the procedure. An incision was then made at the cisterna magna and the skin retracted to expose the dura mater immediately above the 4^th^ ventricle. A 22-gauge needle was then stereotaxically inserted through the dura mater at the following coordinates (anterior=0.3mm, lateral=±0.15mm and ventral= 0.3mm) from the calamus scriptorius and a 20nL volume of lidocaine hydrochloride (20mg/kg) or phosphate buffer injected over a 30s period. The pipette was removed 2-minutes post injection and the surgical site closed using absorbable suture material for muscle and the skin closed with nylon sutures. Animals were allowed a 30 minutes incubation under 1% isoflurane, prior to pFUS application and OGTT.

For IML injection, rats were placed into a stereotaxic instrument equipped with a rat-spinal unit. The 5^th^ and 12^th^ thoracic vertebrae were identified and then rigidly fixed into the spinal unit. The dorsal surface of the spinal cord was exposed by laminectomy and irrigated with warm (37 C) paraffin oil to prevent drying of the spinal cord. The IML was identified at the following coordinates; 0.5mm lateral to midline and 0.7mm ventral from the dorsal surface of the spine. A micropipetter was then stereotaxically inserted into the IML and a 20 nL volume of lidocaine hydrochloride (20mg/kg) was injected over a 30-s period delivered by a controlled syringe pump. The pipette was removed 2-minutes post injection and the surgical site closed using absorbable suture material for muscle and the skin closed with nylon sutures. Animals were allowed a 30 minutes incubation under 1% isoflurane, prior to pFUS application and OGTT.

##### Hepatic Injection of TRPA1 Blocker Amiloride and Analysis of Effect on Hepatic pFUS

Initial pre-study glucose values were collected using a handheld glucometer Freestyle Freedom Lite (Abbot Diabetes Inc., Alameda, CA, US). Following, glucose measurements amiloride (1mg/kg) was administered to the region immediately adjacent to the porta hepatis via an ultrasound-guided percutaneous injection using the GE Vivid E9 Imaging system (GE Healthcare). In brief, sonography was used to identify the portal vein relative to the hepatic artery and common hepatic duct, which serve as anatomical markers for the porta hepatis. Following identification of the porta hepatis, a single 27 gauge needle was guided into the lower portion of the caudate lobe immediately adjacent to the hepatic artery and portal vein and a small volume (50uL) of amiloride was administered. Caution was taken to not disrupt the artery or portal vein to reduce blood loss. Animals were then observed for signs of internal blood loss before they were allowed to recover from anesthesia.

#### Modified Western Diet Mouse Model

##### Animals

Experiments were performed on male C57BL/6J mice (8 weeks old, Jackson Lab, Bar Harbor, ME, USA). All procedures performed in accordance with the National Institutes of Health (NIH) Guidelines under protocols approved by the Institutional Animal Care and Use Committee (IACUC) of the Feinstein Institutes for Medical Research, Northwell Health.

##### Experimental design

6–8 week old C57BL/6J mice (Jackson Laboratories, Bar Harbor, ME) were fed regular chow for 10 d in a reverse light cycle room, and then switched to a high-fat diet (D12492, 60% kcal from fat), or its corresponding isocaloric low-fat diet (10% kcal from fat) for 16 weeks. Mice in the high-fat WD group received sugar supplemented water (55% fructose, 45% sucrose). After 8 weeks, the WD-fed mice were divided into two groups, either treated with pFUS of the porta hepatis (once daily) or sham stimulation for the following 8 weeks. After 8 weeks, the low-fat control diet mice were treated with either the pFUS or the sham stimulation for the remaining 8 weeks (represented in Figure 1B). A second cohort of mice underwent the same dietary regiment, however only received alternate-day hepatic pFUS during the stimulation period (week 9–16, Figure 1C). Blood glucose levels of the mice were monitored on a weekly basis. At week 9, (pre-stimulation period) and week 16 (post-stimulation period) insulin levels were evaluated. Prior to euthanasia, mice were subjected to a glucose tolerance test.

##### Peripheral focused ultrasound stimulation

Mice were anesthetized at 2% isoflurane at 1 L/min O_2_. Mice were then placed on a water circulating warming pad, with a rectal thermometer probe to maintain body temperature. The area above the stimulation target was shaved and hair was fully removed with Nair. The porta hepatis was localized using a custom ultrasound imaging device (GE Healthcare). The location was marked with a permanent marker and a focused ultrasound stimulation probe (GE Healthcare) was placed on the target area (represented in Figure 1A). The device then delivered 1 min of stimulation (1.1 MHz, 200mV per pulse, 150 burst cycles, 500 s burst period), followed by a 30-s period of rest, then a subsequent 1 min of stimulation.

##### Blood glucose determination

Blood glucose levels were assessed weekly by cheek bleed and using a Freestyle blood glucose monitoring system (Abbott Diabetes Inc., Alameda, CA, USA) with Freestyle blood glucose strips following the manufacturer’s recommendations. Mice were fasted 3 h prior to blood glucose assessment. In order to recover fluid volume after bleeding, mice were given a 100 µL injection of saline IP.

##### Glucose Tolerance Tests

At the end of the 16-week period, mice from the four experimental groups were subjected to a glucose tolerance test. The mice were fasted overnight (18 h), weighed, and injected with glucose (10% D-glucose solution; Sigma, St. Louis, MO, USA; 1 g/kg; IP). Glucose levels were determined at 0, 15, 30, 60, and 120 min after glucose administration in blood from the tail vein by using Freestyle glucose monitoring test strips. Glucose tolerance curves were generated from the continuous data and the area under the curve was calculated per graph.

##### Blood collection

After a morning fast (3–4 hr) blood was collected at week 9 and week 16 using the cheek bleed method. Approximately 300 µL of whole blood was sampled per animal. Blood samples were spun in a centrifuge (10 min at 5000 rpm, then 2 min at 10000 rpm) and the serum was extracted and frozen for further evaluation.

##### Serum blood biochemistry tests

Serum samples were centrifuged from whole blood drawn by cheek bleeding (10 min at 5000 rpm, then 2 min at 10000 rpm). The samples were then analyzed with a Millipore MILLIPLEX panel assay for insulin levels.

##### Insulin resistance evaluation

Glucose and insulin levels were utilized to determine insulin resistance by applying the homeostatic model assessment-insulin resistance (HOMA-IR) formula. The HOMA-IR is defined as fasting insulin (microU/L) x fasting glucose (nmol/L)/ 22.5.

#### STZ-treated Diabetic Rat Model and Hyper-insulinemic-Euglycemic Clamp

<u>Animals:</u> Male Sprague-Dawley rats were purchased from Charles River Laboratories (Wilmington, MA) and housed in the Yale Animal Resource Center in temperature (22-23°C) and humidity-controlled rooms. Animals fed with high fat diet (HFD, DIO model) or standard rat chow (STZ T1D model) and water ad libitum were acclimatized to a 12 h light cycle. All the animal procedures were approved by the Yale University Institutional Animal Care and Use Committee. <u>DIO model</u>: Male rats (150-161 g) were fed with HFD for two weeks (Body weight around 300 g) and then underwent vascular surgery for catheter implantation into the left carotid artery (for blood glucose and hormone sampling) and into the right jugular vein (for insulin and glucose infusions). Four days later, a single intraperitoneal injection of STZ (35 mg/kg dissolved in 0.1M sodium citrate) was administrated followed with daily blood glucose monitoring and subcutaneous saline injection to maintain hydration. A week after steady high blood glucose, rats underwent either pFUS or sham ultrasound treatment (see above for US-parameters) for three consecutive days prior to study via insulin clamp. <u>Lean STZ T1D model</u>: Male rats (250-280 g) were given a single intraperitoneal injection of STZ (65 mg/kg dissolved in normal saline). A second dose of STZ was introduced if the blood glucose of rats fell below 400 mg/dl 24 hours after the first injection. 5 days post STZ injection diabetic rats were given long-acting protamine zinc insulin (PZI) subcutaneously for daily treatment to maintain elevated blood glucose levels (300-550 mg/dl) but to avoid blood glucose extremes and ketosis and the associated loss in body weight. In addition, the rats received daily subcutaneous saline injections, if needed to maintain hydration. After stable diabetes induction and maintenance for 10 days animals were anesthetized and underwent aseptic surgery with vascular catheter implantation into the left carotid artery (for blood glucose and hormone sampling) and into the right jugular vein (for insulin and glucose infusions). Following a 7-day recovery period all animals underwent either pFUS or sham ultrasound treatment (see above for US-parameters) for three consecutive days prior to study via insulin clamp. <u>Hyperinsulinemic Euglycemic Clamp:</u> The catheters of overnight-fasted rats were connected to infusion pumps in the morning of the study and then left undisturbed to minimize handling stress for at least 60 min before baseline sampling. Euglycemia was induced via a bolus infusion of human insulin (0.025 units/kg/min; Human R U-100; Eli Lilly) for 30 minutes (DIO model) or 30-50 minutes (STZ T1D model) followed by a constant infusion (0.015 units/kg/min in DIO model or 0.01 units/kg/min in STZ T1D model) of human insulin together with a variable infusion of 20% glucose for the remainder of the clamp. Plasma glucose was measured every 5 to 10 min throughout the study and levels were gradually lowered to 110 mg/dL and maintained at this target for the remaining 90 min. Blood samples were collected at baseline and at 30, 60, 90 and 120 min of the clamp. Plasma was separated immediately and snap frozen in liquid nitrogen, then stored either at −20°C (For insulin, glucagon and corticosterone measurement) or −80°C (for catecholamine measurement). The animals were sacrificed at the end of clamp procedure for further endpoint tissue analyses as necessary.

#### Swine Model and Hyper-insulinemic-Euglycemic Clamp

All experimental procedures and protocols were reviewed and approved by New York Medical College Animal Care and Use Committee.

Four adult male Yucatan miniature pigs (Exemplar Genetics, Sioux Center, IA) weighing 50-55 kg were examined by attending veterinarian upon arrival to the facility. None of the animal expressed any signs or symptoms of being unhealthy.

Animals went through a 5-10 day acclimation period before the hyperinsulinemic-euglycemic clamp (HEC) experiment took place. Each animal was randomized and assigned to a different experiment. Due to inter-species variability, animals were subjected to initial hyperinsulinemic-euglycemic step clamp procedures where insulin was infused at a fixed different dose (0.3-4 mU/kg/min) to establish the appropriate insulin-response curve. After the insulin-response curve was determined, each animal underwent an additional 2-4 HEC procedures to evaluate the effect of noninvasive focused ultrasound (FUS) on insulin sensitivity. Finally, each swine was subjected to a non-survival procedure, during which porta hepatis (PH) was approached surgically and FUS was delivered directly to it.

##### Sedation and Anesthesia Procedures

Sedation and Anesthesia procedures for: a) Vascular Access Procedures, b) Survival hyperinsulinemic euglycemic clamp (HEC) Procedures, and c) Non-Survival HEC Procedures.

For a 12-hour period before surgery, the animals were maintained on a no food and no fluid regimen. On the day of surgery, the animal was weighed and sedated with Telazol (2-4 mg/kg IM) and transported to the operating room (OR). Once in the OR, 2-4% isoflurane inhalation anesthetic mixed in oxygen was administered by a mask. In a supine position, an endotracheal tube was placed, and surgical anesthesia was maintained on isoflurane and oxygen (2-4% induction, 0.5-3% maintenance). Ventilator settings was adjusted to maintain the end-tidal carbon dioxide at 40±3 mmHg. The depth of anesthesia was monitored by assessing heart rate, blood pressure, respiration, mandibular jaw tone, absence of corneal reflex and absence of withdrawal response to toe pinch. After successful intubation, a catheter was placed in an ear vein and maintenance fluids was administered (0.9% NaCl at 5-10ml/ko/hr). Also, ECG and pulse oximetry were monitored. Body temperature is controlled with a recirculating warming blanket and/or a hot air warmer.

##### Vascular Access Procedures

<u>Infusion port (venous side):</u> With continued surgical level of anesthesia, the skin of the right or left ear was shaved, cleansed, sterilized, and draped with a surgical field. A (20G) intra venous catheter was placed in an ear vein, secured, and flushed with heparinized saline. The port was connected to a dual infusion pump (Kd Scientific) and used to infuse insulin and glucose. <u>Blood samples withdrawals port (arterial side):</u> **Carotid artery catheter placement:** The animals was anesthetized as described above. In a dorsal recumbency position, the neck was shaved and prepared with ethanol and betadine surgical scrub to disinfect the area. A right or left, ventral incision (6-8 cm) was made 1 cm lateral and parallel to midline in the cervical region of the neck. Subcutaneous tissue and muscle layers were bluntly dissected to expose the carotid sheath and incised to expose the carotid artery. The carotid artery was isolated and permanently ligated at the rostral end using 3-0 Silk suture. A tension suture placed around the caudal end to control the bleeding and a small sharp incision was made through the arterial wall and a catheter (polyethylene ID 0.04-0.06 inches) was inserted into the lumen and secured in place using 3-0 Silk sutures. The skin and the underlying layers were closed, and the catheter tunneled to the intrascapular region. At the end of the procedure, the animal was returned to the Animal Facilities Recovery area by a Comparative Medicine technician and was monitored continuously until sufficiently recovered. The animal was monitored at least 3 times/day for 72 hours. Animal was allowed to recover for 3-5 days and then was randomly assigned for one of the protocols experiments. The patency of catheters was maintained by flushing with heparinized saline at least once a week and up to 3 times a week. A locking solution (200 U Heparin / ml saline 0.9 NaCl) was used to prevent formation of clots and was replaced weekly. **Percutaneous puncture technique:** The skin above the puncture site was shaved, cleansed, sterilized, and dressed in a surgical field. Under ultrasonic guidance, a 19G angiographic needle was passed slowly through the skin at 60-degree angle to the target vessel. Once blood backflow is detected (single wall puncture technique), a soft guide wire (0.035 inch) was introduced into the vessel. Under direct vision and firm vessel compression, the needle was removed and exchanged with a 5 Fr introducer sheath over the sliding wire and secured in place. This technique was used in the event of the failure of the carotid artery catheter (due to blockage or was destroyed) during the first clamp procedure and prior to beginning the infusion.

##### Survival Hyperinsulinemic Euglycemic Clamp Procedures

Following the induction of anesthesia, fasting blood glucose level was measured, which was an average of three fasting blood samples taken immediately before the clamp (−30, −15 and −5 minutes). A short acting soluble human Insulin (Humalog U-100) infusion (0.3-4mU/kg/min) was initiated at 0 minutes and glucose (Dextrose 20%) infusion (20-250mL/h of a 200 mg/mL solution) was given 5 minutes afterwards. The glucose infusion rate was regulated based on frequent plasma glucose measurement (5-minute intervals) to keep the pigs within +/− 10 mg/dl of their fasting glucose level. Heparinized 0.3 mL of whole blood for each time point was used to measure blood glucose, using i-stat handheld blood analyzer (Abbott). Samples (1 mL in EDTA tubes) for insulin measurement were taken at 15, 30, 60, 90, 120, 150, 180 and 210 minutes and stored for further analysis

After establishing the HEC, approximately within 90-100 min, a diagnostic ultrasound (U/S) system (GE Vivid E10) was used to visualize the liver to identify the anatomical biomarkers relevant to the target nerve. In the supine position, the lower chest and upper abdomen was shaved and cleaned for a proper imaging. Full sweep of the liver was performed using GE C1-6 Curved Array Probe 1-6 (MHz) at right subcostal region and a full scan was obtained visualizing the gallbladder and the large vessels. Doppler imaging of the portal vein was used to identify the porta hepatis which located approximately 8 cm in depth. Following diagnostic imaging of the liver and identifying the porta hepatis, the same U/S probe was placed above the target site and used to apply U/S stimulation with defined frequency and pulse length parameters, for up to 4 minutes total duration. Mechanical ventilation was interrupted for a period of 30-45 seconds to ensure that the target is not being moved. The ultrasound pulse was applied at full power: maximum intensity of 26895mW/cm2, maximal pulse width of 363.3us, maximum frequency 2.2MHz and maximum pulse length 400 cycles. Glucose levels and glucose infusion rates were recorded again as an indication of insulin sensitivity for additional 90 minutes post FUS. At the end of the experiment, insulin infusion was stopped followed by stopping glucose infusion after 45 minutes. Consecutive blood samples were confirming the normal glucose level (Euglycemia).

At the end of the experiment, the physiological parameters were monitored, and consecutive blood samples were confirming the normal glucose level (Euglycemia). Then the animal was gradually weaned off the ventilator and extubated, provided that all physiological parameters are within normal limits. Adequate perfusion, normal blood sugar, temperature, and physiological signs were evaluated by the Veterinary Staff. The animal was returned to the Animal Facilities Recovery area by a Comparative Medicine technician and was monitored continuously until sufficiently recovered. The animal was monitored at least 3 times/day for 72 hours. Animal was allowed to recover for 3-5 days and then was assigned for the next experiment.

##### Non-Survival Hyperinsulinemic Euglycemic Clamp Procedures

Following the induction of anesthesia and while maintaining the anesthesia with isoflurane (2-3%), using a sterile technique and appropriate surgical attire, a midline surgical incision was made vertically into the skin, extend caudally 10-12 cm from the xiphoid process. Subcutaneous tissue and muscle layer were blunt dissected, and hemostasis was achieved suing diathermy. The peritoneum was identified and carefully opened parallel to the surgical incision. A self-retractor was placed, and the abdominal cavity was exposed. The internal structure/organs were gently mobilized to visualize a clear field of the liver and its anatomical structures. The liver borders were identified, and the inner surface was carefully lifted upward to gain access, with direct visualization, of the site where the major vessels and ducts enter or leave the liver (porta hepatis). Following visualization of porta hepatis, GE C1-6 probe, in a sterile sleeve, was placed and separated from the target site using sterile ultrasound gel pad, approximately 4 cm thickness (Aquaflex). Following the invasive surgical procedure for the US probe placement, animal fasting blood glucose level was measured, which was an average of three fasting blood samples taken immediately before the clamp (−30, −15 and −5 minutes). A short acting soluble human Insulin (Humalog U-100) infusion (0.3-4mU/kg/min) was initiated at 0 minutes and glucose (Dextrose 20%) infusion (20-250mL/h of a 200 mg/mL solution) was given 5 minutes afterwards. The glucose infusion rate was regulated based on frequent plasma glucose measurement (5-minute intervals) to keep the pigs within +/− 10 mg/dl of their fasting glucose level. Heparinized 0.3 mL of whole blood for each time point was used to measure blood glucose, using i-stat handheld blood analyzer (Abbott). Samples (1 mL in EDTA tubes) for insulin measurement were taken at 15, 30, 60, 90, 120, 150, 180 and 210 minutes and stored for further analysis.

After establishing the HEC, approximately within 90-100 min, a diagnostic doppler ultrasound (U/S) system (GE Vivid E10) was used to visualize the portal vein with used to identify the porta hepatis which located approximately 5 cm deep from the US probe. Following diagnostic imaging of the porta hepatis, the same U/S transducer was placed above the target site and used to apply U/S stimulation with defined frequency and pulse length parameters, for up to 4 minutes total duration. Mechanical ventilation was interrupted for a period of 30-45 seconds to ensure that the target is not being moved. The ultrasound pulse was applied at full power: maximum intensity of 26895mW/cm2, maximal pulse width of 363.3us, maximum frequency 2.2MHz and maximum pulse length 400 cycles. Glucose levels and glucose infusion rates were recorded again as an indication of insulin sensitivity for additional 90 minutes post FUS. At the end of the experiment, insulin and glucose infusion were stopped and while maintained the general anesthesia, swine was euthanized with a solution (Euthasol).

At the end of the experiment and while maintained the general anesthesia, swine was euthanized with a solution (Euthasol) (A dosage of 1 ml/10 lb body weight, intravenously [IV]). Euthanasia solution (Euthasol) was injected IV. Death was confirmed using the ECG signal and the arterial blood pressure signal. Absence of a pulse in both signals confirms cardiac death. After euthanasia, tissue samples were harvested for histological and biochemical analysis.

#### Direct Neural Recording in the Paraventricular Nucleus (PVN) in Rats

##### Animals

All animal use in our study complied with the National Institutes of Health and Albany Medical College (AMC) Institutional Animal Care and Use Committee (IACUC) guidelines. Male Sprague Dawley rats weighing 225-300g were purchased from Taconic (Germantown, NY, USA) and had access to food and water *ad labitum* except when fasted for 16-18 hours prior to experiment and craniotomy surgery.

##### Peripheral focused ultrasound stimulation

All animals were anesthetized with an intraperitoneal (IP) injection of urethane (1.2-1.5g/kg in saline). The fur in the abdominal area over the liver removed and ultrasound gel placed on the shaved surface. The hepatic portal vein was located with an ultrasound imaging scanner and handheld transducer and marked with a black dot in initial experiments. Animals in subsequent experiments had the black dot on the same abdomen location and positioned on a gel-containing platform so the black dot on the abdomen was directly on the center of the gel where the pFUS transducer was positioned. Next, a hypodermic needle was inserted in the IP space.

##### Craniotomy surgery and *in vivo* electrophysiological recordings

Animals were placed in a stereotactic frame and burr holes were made. Tungsten microelectrodes with an impedance of 300-500 kΩ or a 12-channel multi-array linear recording electrode (Microprobes, Gaithersburg, MD) were inserted in the right or left PVN^122^. Single-unit neuronal spiking activity was acquired and amplified 10,000x, sampled at 40 kHz and band-pass filtered between 300 and 6000 Hz with a multi-channel acquisition system (OmniPlex, Plexon, Dallas, TX) as before.^123–126^

PVN neurons with amplitudes of at least a 2:1 signal-to-noise ratio were identified and monitored for 60 minutes for a baseline recording. After, we injected 2 ml of 20% glucose or saline into the catheter to deliver one or the other solution IP. One hour of recordings later, we administered pFUS to the hepatic portal vein and another 60 minutes of single-unit recordings were obtained. As a control, PVN spiking activity was monitored for 60-minutes prior to an IP injection of glucose or saline with a subsequent post-injection monitoring period of 120 minutes.

##### Blood glucose monitoring

We obtained blood glucose samples from the tail vein every 20 minutes concomitant with PVN monitored neuronal activity. To do so, a small incision was made on the tail and the glucose level was noted using a small handheld device (Verio Onetouch).

##### Transcardiac perfusion with paraformaldehyde and nissl cresyl violet staining

After the experiment, animals were transcardiac-perfused with heparinized saline followed by 4% paraformaldehyde (PFA) in phosphate buffered saline (PBS). Brains were extracted and then placed in 30% sucrose to cryoprotect the brain. Sections were cut 60 μm thick (HM500 M cryostat, Leica Biosystems Inc. IL USA) and mounted on slides. Electrode placement was confirmed with cresyl violet (CV) for nissl substance as described elsewhere.^123,124,127–129^ CV stained slides were imaged using the PathScan Enabler IV (Electron Microscopy Sciences, Hatfield, PA) and co-registered onto a rat brain atlas^122^ to determine electrode location.

##### Chemicals and drugs

All chemicals were purchased from Sigma Aldrich (St. Louis, MO, USA) unless otherwise specified.

##### Data analysis and statistics

We identified PVN neuronal cells by their waveform and confirmed electrode placement in the PVN. For the former, offline cell sorting was performed using a Matlab Wave clust toolkit ^9^. Only single-units with amplitudes having at least a 2:1 signal-to-noise ratio were sorted based on wavelets and superparamagnetic clustering.^130^ Sorted spikes were processed using custom-made Matlab scripts (www.mathworks.com), which concatenated each 20 minute electrophysiological recording interval into a single continuous trace and plotted as 1 minute moving averages with standard error of means (SEMs) for the single-unit spiking activity across the entire recording. For between-group analyses, spike rates were normalized by the maximum spike rate in the 3 hour recording. To identify glucose-sensitive neurons, the spike rates 1 hour pre (“baseline period”) and post (“glucose period”) glucose injection were compared. Neurons after glucose administration with spike rates greater than the mean ± 1 S.D. of the baseline period were classified as GE neurons whereas those with spike rates less than the mean ± 1 S.D. of the baseline period were classified as GI neurons. Otherwise, PVN cells were classified as NRs to glucose.

Significance of changes in glucose levels and spike rates were calculated using non-parametric Wilcoxon’s rank sum statistical test.

#### Mechanical vs. Ultrasound Neuromodulation

##### Stimulation Systems and CAP Neuromodulation

Two stimulation systems were used for the animal study: a single-element High Intensity Focused Ultrasound (HIFU) system, and a mechanical vibrator (TIRA GmbH Model: Tira Vib S 502 shaker).

The pre-clinical ultrasound transducer system used was the same shown above and previously used for hepatic pFUS. The function generator (Agilent 33210A) was used to generate a tone burst, with a 1.1 MHz center frequency, 133 us pulse length, 500 us (or 200 ms) pulse repetition period and 200 mV peak-to-peak amplitude. The tone burst signal was again amplified by the power amplifier (E&I 350L). The output of the power amplifier was sent to the input of the HIFU matching network. The polyurethane coupling cone was used as the standoff, and the focus of the HIFU (full width at half maximum (FWHM); approximately 2 mm wide and 10 mm long with a measured peak negative pressure of 2.4 MPa) was targeted at the center of the spleen for this experimental activation of the cholinergic anti-inflammatory pathway. (ref) The stimulus was applied for 3 minutes in duration as reported previously. (ref)

The TIRA Vibration System (TV 50018) consists of Shaker S 502 and Amplifier BAA 60 (TIRA GmbH). A function generator (Agilent 32210A) was used to generate a sine wave at four specified frequencies (100, 500, 1000, 2000 Hz) with a pulse length of 133 us, a pulse repetition interval of 500 us and a peak-to-peak voltage of 200 mV. The stimulus frequency was swept across the four values throughout the 3-minute stimulation period. The mechanical piston was actuated at peak force (18 N) during the swept actuation (at frequencies 100 - 2000 Hz that are associated with the stimulation pulse frequency ranges associated with both Transient Shear Wave Elastography (ref) and previous reports of ultrasound-based neuromodulation (ref)). Both the ultrasound and TIRA systems were used to stimulate CAP for the same 3-minute treatment period. The focused ultrasound systems provide tissue displacement primarily near the ultrasound focus, while the TIRA vibrator was used to actuate tissue surface displacements that propagate to the focal spot. The TIRA generates displacements at the level of depth associated with the ultrasound focal points (i.e. 3-5 mm) that are of the same order of magnitude as the tissue displacement generated by the ultrasound system. ^(ref)^ The cholinergic anti-inflammatory activation was chosen as the nerve pathway (for comparison between ultrasound and mechanical actuator-based neuromodulation) as splenic norepinephrine release and corresponding TNF cytokine output enable rapid and quantitative assessment of both nerve pathway activation and down-stream neuromodulation effect. (ref) The LPS-induced inflammation model was utilized in experiments (see below), as this is a standard model to rapidly assess the level of cholinergic anti-inflammatory pathway activation based on pharmacologic and device-based intervention. (ref)

##### Preclinical induction of inflammation by lipopolysaccharide administration

Adult male Sprague–Dawley rats 8–12 weeks old (250–300 g; Charles River Laboratories) were housed at 25 °C on a 12-h light/dark cycle and acclimatized for 1 week, with handling, before experiments were conducted to minimize potential confounding measures due to stress response. Water and regular rodent chow were available ad libitum. Experiments were performed under protocols approved by the Institutional Animal Care and Use Committee of GE Global Research.

An LD75 dose of the endotoxin, lipopolysaccharide (LPS, from Escherichia coli, 0111: B4; Sigma) was administered by intraperitoneal (IP) injection to naive adult Sprague–Dawley rats, causing a significant elevation in concentrations of pro inflammatory cytokines. Elevation in cytokine concentrations peak in 4 h, but remain elevated as compared to control for up to 8 h post injection. After LPS administration, tissue and fluid samples were collected at 60 min post LPS administration and US stimulation.

Tissues were collected at the terminal time point as follows: an incision was made starting at the base of the peritoneal cavity extending up and through to the pleural cavity. Organs (eg spleen) were rapidly removed and homogenized in a solution of phosphate-buffered saline (PBS), containing phosphatase (0.2-mM phenylmethylsulfonyl fluoride, 5-µg/mL aprotinin, 1-mM benzamidine, 1-mM sodium orthovanadate, and 2-µM cantharidin) and protease (1-µL to 20 mg of tissue as per Roche Diagnostics) inhibitors. A targeted final concentration of 0.2-g tissue per mL PBS solution was applied in all samples.

Blood samples were stored with the anti-coagulant disodium (ethylenedinitrilo) tetraacetic acid (EDTA) to prevent coagulation of samples. Samples were then stored at −80 °C until analysis.

Cytokine concentration in tissue and fluid samples was determined by multi-analyze ELISA assay. A detailed protocol for ELISA Assays was provided by the respective supplier of the kits:

https://www.qiagen.com/us/products/discovery-and-translational-research/functional-and-cell-analysis/elisa-assays/multi-analyte-elisarray-kits/#orderinginformation

##### HPLC analyses of Circulating Neurotransmitter

Serum samples were injected directly into the machine with no pre-treatment. Tissue homogenates were initially homogenized with 0.1-M perchloric acid and centrifuged for 15 min, after which the supernatant was separated, and the sample injected into the HPLC with an inline ultraviolet detector. The test column used in this analysis was a Supelco Discovery C18 (15-cm × 4.6-mm inside diameter, 5-µm particle size). A biphasic mobile phase comprised of [A] acetonitrile: [B] 50 = mM KH2PO4, set to pH 3 (with phosphoric acid). The solution was then buffered with 100-mg/L EDTA and 200-mg/L 1-octane-sulfonic acid. Final concentration of mobile phase mixture was set to 5:95, A:B.

A flow rate of 1 mL/min was used to improve overall peak resolution while the column was held to a consistent 20 °C to minimize pressure compaction of the column resulting from the viscosity of the utilized mobile phase. The UV detector was maintained at a 254-nm wavelength, which is known to capture the absorption for catecholamines including norepinephrine.

##### In vitro TNF analysis in whole blood

In brief, terminal blood samples were collected into sterile heparin collection tubes (BD; #366480). Individual blood aliquots of 500uL were transferred into 12 polystyrene tubes containing either 0, 0.1, 1 or 10 ng/mL LPS. Concentration of LPS were tested in triplicate by ELISA (described below).

Each polystyrene tube was then mixed by gentle inversio before being secured on a lab rocker. Samples were then transferred to 37C for 4 hours with constant rocking. After 4 hours, samples were removed from the incubator and centrifuged at 6000 RPM for 5 minutes. The plasma supernatent was then removed and TNF concentrations analyzed by ELISA. A detailed protocol for ELISA Assay was provided by the supplier https://www.abcam.com/rat-tnf-alpha-elisa-kit-ab100785.html

#### In Vitro Ultrasound Neuromodulation and Chemical Ion Channel Blocking

##### Microfluidic device fabrication for production of hydrogel scaffolds

For 3D culture of peripheral neurons we use hydrogel particle scaffolds fabricated using previously reported techniques.^14,31,131^ Briefly, a flow focusing microfluidic device was used for manufacturing hydrogel spheres that were assembled in a porous scaffold for in vitro neuron growth. Microfluidic devices were fabricated using soft lithography techniques. Master molds were fabricated on mechanical grade silicon wafers (University wafer) using KMPR 1050 photoresist (Microchem). Devices were molded from the masters using poly(dimethyl)siloxane (PDMS) Sylgard 184 kit (Dow Corning). The base and crosslinker were mixed at a 10:1 mass ratio, poured over the mold, and degassed prior to curing overnight at 65 °C. Channels were sealed by treating the PDMS channels and a glass microscope slide (VWR) with oxygen plasma at 500 mTorr and 80 W for 30 s. The channels were functionalized by injecting 100 µL of Aquapel (Aquapel) and reacting for 30 s until washed by Novec 7500 (3M). The channels were dried by air suction and kept in the oven at 65 °C until used.

##### Synthesis and characterization of thiolated hyaluronic acid (HA)

Hydrogel spheres for neural growth used thiolated HA as a precursor to enhance neural cell growth. Sodium hyaluronate (700 kDa, LifeCore Biomedical) was dissolved at 10 mg/mL in distilled DI water. The pH of the HA solution was adjusted to 5.5 using 0.1M HCl. 1-Ethyl-3-[3-dimethylaminopropyl] carbodiimide (EDC) was dissolved in DI water at the appropriate molar ratio (0.25 × unless otherwise stated) immediately before addition to HA solution. Molar ratios in all cases are reported with respect to carboxyl groups on glucuronic acid moieties of HA. N-hydroxysuccinimide (NHS, Acros Organics) was then added at half of the molar ratio as EDC. The pH was then readjusted to 5.5 and the reaction was mixed at room temperature for 45 min.

Then, cystamine dihydrochloride (Sigma-Aldrich) was added (0.25 × molar ratio), pH was adjusted to 6.25 using 0.1 M NaOH, and the reaction continued while stirring at room temperature overnight. Dithiothreitol (DTT, Sigma-Aldrich) was added in excess (4 × greater than cystamine) at pH 8. The mixture was stirred for 1–2 h to cleave cystamine disulfides and yield thiolated HA (HA-SH). The reaction was quenched by adjusting the pH to 4. HA-SH was purified using dialysis against acidic (pH 4) DI water for 3 days in the dark. Purified, HA-SH was filtered through a 0.22 µm filter (EMD Millipore), frozen under liquid nitrogen, lyophilized, and stored at − 20 °C until use. HA thiolation was confirmed using the colorimetric Ellman’s test for free thiols.

##### Fabrication of microparticles for hydrogel scaffold

Hydrogel microparticles that assemble to provide a porous scaffold for in vitro culture of neurons were manufactured from PEG and HA using a microfluidic flow focusing droplet generator. HA-SH and 4-arm vinyl sulfone-terminated PEG (PEG-VS) (20kDa, Laysan Bio) were crosslinked via Michael-type addition between thiol and vinyl sulfone functional groups. HA-SH and PEG-VS were dissolved separately in PBS at pH 7.4. Cysteine-terminated RGD peptide (GCGYGRGDSPG, GenScript Biotech) was conjugated to PEG-VS by reaction at room temperature for 1 h prior to gel formation to provide additional sites for cell adhesion. These pre-gel solutions were sterile-filtered through a 0.2 µm polyethersulfone (PES) membrane in a leur-lok syringe filter, injected into the microfluidic device and pinched off by oil phase (0.1% Pico-Surf in Novec 7500, SF-000149, Sphere Fluidics). The flow rate for aqueous solutions was 10 µL/min and for oil solutions was 50 µL/min. Gels were collected from the device into a tube in oil phase, incubated overnight at room temperature in the dark. Microgels in the oil phase were vortexed with 20% 1H,1H,2H,2H-Perfluoro-1-octanol (PFO) (Sigma-Aldrich) in Novec 7500 for 10 s. Microgels were then mixed with 1:1 mixture of HEPES buffer (100 × 10 mM HEPES, 40 × 10 mM NaCl pH 7.4) and hexane followed by centrifugation at 10000 rpm to separate microgels from oil for five times. Microgels were incubated in sterile filtered 70% ethanol solution at 4 °C at least overnight for sterilization. Before experiments, microgels were washed with HEPES buffer five times.

##### Dissociation and encapsulation of dorsal root ganglion cells in the porous scaffold

Embryonic (Day 18) dorsal root ganglion (DRG) neuron cells from rats were dissociated from tissue (BrainBits) and incorporated into the microporous hydrogel scaffold. To dissociate DRG neurons we first prepared a 100 mg/ml stock solution of Collagenase/Dispase in DI water. The stock solution is diluted 1:100 with Hibernate E storage media without calcium and magnesium (BrainBits) for a final working solution of 1 mg/ml collagenase/dispase (0.1U/0.8U). The primary DRG tissue was placed in cell dissociation solution for 1 hour and incubated at 37 ℃ with gentle swirling every 5 min. Following incubation, tissue is allowed to settle and cell dissociation solution is removed leaving the tissue at the bottom in minimal solution. Immediately following this 2ml of Hibernate AB (Brainbits) is added. With a silanized pasteur pipette, the tissue was triturated ∼30 times until 90% dispersed. Supernatant containing the dispersed cells was transferred to a sterile tube and spun down at 1100 rpm for 3 min. The pellet of cells was re-suspended in 1ml of culture media (NbActiv4 with 25 ng/ml NGF, BrainBits).

Previously prepared HA-PEG microgels were re-suspended and 40 µL of the mixed solution was injected into a well-plate followed by 90 min of incubation at 37 °C to allow gels to anneal to form a microporous annealed particle scaffold.^132^ To incorporate the neurons, aliquots of neurons (4 µL, 50 000 cells) were injected into the porous scaffolds 45 min into gelation. Cell-laden hydrogel scaffolds were incubated for another 45 min at 37 °C until full annealing is achieved before addition of culture media (150 µL per well in a 96 well-plate).

##### Calcium dye incubation and ultrasound stimulation

Primary DRG neurons within porous hydrogel scaffolds were incubated with Fluo-4 Direct calcium assay kit with 250 × 10 mM stock solution of probenecid. Briefly, 5 mL of calcium assay buffer was mixed and vortexed with 100 µL of probenecid stock solution to create a 2× loading dye solution. The dye solution was then added to the cells with media in a 1:1 ratio and incubated for 1 h before imaging to allow sufficient diffusion of dye through the hydrogels. For experiments involving TTX (1 μM), ω-conotoxin GVIA (1 μM), GsMTx4 (10 μM), HC-030031 (10 μM), the chemical was added during calcium dye incubation (1 h). Incubated cells were observed under microscopy during pFUS stimulation. The ultrasound transducer, function generator, and power amplifier system utilized for pre-clinical hepatic stimulation (and described above) was coupled to each individual culture well using a custom plastic fitting on the culture well plate. The plastic fitting allowed for filling of the culture plate with additional growth media, and placement of the plastic standoff cone (described above) directly on top of this liquid layer for acoustic coupling. The height above the growth surface of the culture chamber was set to 12 mm to match the focal position of the transducer (and focal targeting used in the pre-clinical studies), and the ultrasound stimulus was centered within each neuron culture well using the 20 mm diameter hole in the center of the transducer (previously used to insert a small imaging transducer for targeting in the pre-clinical studies described above).

#### Metabolomics and Transcriptomic Measurements

##### Untargeted Metabolomics

Untargeted metabolomics from blood serum and liver were performed by Metabolon, Research Triangle Park, NC.^133^ All samples were prepared at Metabolon using their automated MicroLab STAR® system from Hamilton Company. Briefly, small molecules are extracted with methanol under vigorous shaking for 2 min (Glen Mills GenoGrinder 2000) followed by centrifugation. The extract is divided into five fractions: two for analysis by two separate reverse phase (RP)/UPLC-MS/MS methods using positive ion mode electrospray ionization (ESI), one for analysis by RP/UPLC-MS/MS using negative ion mode ESI, one for analysis by HILIC/UPLC-MS/MS using negative ion mode ESI, and one reserved for backup. All methods utilize a Waters ACQUITY ultra-performance liquid chromatography (UPLC) and a Thermo Scientific Q-Exactive high resolution/accurate mass spectrometer interfaced with a heated electrospray ionization (HESI-II) source and Orbitrap mass analyzer operated at 35,000 mass resolution. Quality control samples were introduced and spaced evenly with the randomized experimental samples across the platform run. The median relative standard deviation (RSD) for the internal standards spiked into each of the study samples prior to injection into the MS instrument was 3% for both the liver and blood serum runs. For the technical replicates created from the experimental samples, the RSD was 7% and 9% for the liver and serum respectively. A total of 905 (liver) and 891 (blood serum) compounds were quantified that were within Metabolon’s QC specifications. These compounds were then identified by comparison to Metabolons library (4500+) entries of purified standards or recurrent unknown entities that includes their retention time/index (RI), mass to charge ratio (m/z), and chromatographic data (including MS/MS spectral data). Biochemical identifications was based on: retention index within a narrow RI window of the proposed identification, accurate mass match to the library +/− 10 ppm, and the MS/MS forward and reverse scores computed from the ions present in the experimental spectrum relative to ions present in the library entry spectrum. Metabolon data analysts also used internally developed visualization and interpretation software to confirm and curate the data to ensure accurate identification of true chemical entities, and to remove those due to system artifacts, mis-assignments, redundancy, and background noise. For some identified metabolites, there is a symbol at the end of the metabolite name indicating a specific footnote: * indicates compounds that have not been officially confirmed based on a standard, but Metabolon stated they were confident in its identity; # indicates a compound that is a structural isomer of another compound in the Metabolon spectral library; ** indicates a compound for which a standard is not available, but Metabolon was reasonably confident in its identity or the information provided. A total of 816 (liver) and 792 (blood serum) metabolites were identified from the Metabolon’s library. The metabolite peaks were quantified as area-under-the-curve detector ion counts. For the liver samples, the quantified values were then normalized by total protein (Bradford assay) to correct for systematic variation in metabolite levels due to differences in starting amounts of tissue. All metabolite measures were then re-scaled to have median equal to 1. Missing values were then imputed with the minimum scaled value.

##### RNA Sequencing

RNA extraction from tissues and RNA sequencing were performed at the Roswell Park Cancer Institute, Genomics Shared Resource, Buffalo, New York.^134^ Extractions from tissues yield RNA amounts ranging from 11 to 256 (median 78) micrograms. The quality was assessed by the RNA integrity number (RIN) from a BioAnalyzer (Agilent Technologies). The RIN from all tissue samples ranged from 3 to 9.9 (median 7.4) The samples with the low RIN came from two tissue types, pancreas (median 4.0 RIN) and adipose (median 4.5 RIN). Sequencing RNA libraries were prepared using the TruSeq Stranded mRNA Sample Preparation Kit (Illumina, San Diego, CA) according to the manufacturer’s instructions for the RNA from all tissues except the pancreas and adipose. For these samples due to lower quality RIN scores, the TruSeq Stranded Total RNA Sample Preparation Kit was used. Sequencing was performed on the Illumina NovaSeq 6000 Sequencing platform that output 100-base pair pair-ended reads ranging from 14 to 55 (median 20) million reads per sample. The RNA-seq data was processed at GE Global Research using established bioinformatics software tools. Base quality control was checked and found to be excellent using Fast QC v0.10.1 from Babraham Bioinformatics.^135^ Sequencing reads were mapped (median 71%) to the annotated rat genome version, Rattus norvegicus Rnor 6.0.91, using STAR_2.5.3a aligner.^136^ Transcript abundance estimates were then generated using RSEM^137^ which outputs the expected count for each transcript.

##### Differential transcript expression analysis

Transcript count normalization and differential expression analysis was performed on all samples using the DESeq2 tool.^103^ The p-values attained by the Wald test are corrected for multiple testing using the Benjamini and Hochberg method. Transcripts with an adjusted p value < 0.1 were counted as being differentially expressed. Output from DESeq2 included the median ratio normalization (MRN) values for each transcript of each sample. These normalized values were used for gene set enrichment analysis.

##### Gene set enrichment analysis

Gene set enrichment analysis (GSEA) was performed using GSEA (version 3.0) tool^104^ to identify functional pathways with the gene set collection Gene Ontology (GO) biological processes (C5), Reactome and KEGG curated genes sets (C2) and the hallmark gene sets (H) available at the Molecular Signatures Database (MSigDB).^138^ Biological processes represented by specific gene sets includes: glucose metabolic process [GO:0006006], glucose catabolic process [GO:0006007], glycolysis [Broad Hallmark], gluconeogenesis [Reactome R-HSA-70263], response to insulin [GO:0032868], adipogenesis [Broad Hallmark], oxidative phosphorylation [Broad Hallmark], fatty acid catabolic process [GO:0009062], bile acid metabolism [Broad Hallmark], carbohydrate transport [GO:0008643], vitamin transport [GO:0051180], mitochondrial transport [GO:0006839], synapse organization [GO:0050808, voltage gated potassium channel activity [GO:0005244], mTORC1 signaling [Broad Hallmark], MYC targets v2 [Broad Hallmark], TNF-alpha signaling via NF-kB [Broad Hallmark]. Diabetic associated gene sets were assembled from public data sources including genes associated with type 2 diabetes mellitus from GWAS and other genetic association datasets.^139–141^ For GSEA, the DESeq2 MRN values were inputted into the GSEA tool for each gene in which its transcript had a log2 fold change with a p-value < 0.2. Gene sets identified by the GSEA tool to have a Familywise-error rate (FWER) p-value < 0.1 were considered significant. The FWER was used over the alternatively provided FDR statistics to minimize false positive findings.

## Statistical analysis

All statistical analyses were conducted by the R software (version 3.6). The Wilcoxon rank-sum test, a non-parametric statistical hypothesis test, was used to compare any two related samples unless otherwise stated.

## Data availability

RNA-seq sequencing data are available from the NCBI Gene Expression Omnibus (GEO) repository under accession number **GSE:** to be completed before publication

## Code availability

The analysis source code underlying the final version of the paper available upon request.

## Supplementary Figure and Captions

**Supplemental Figure 1.**
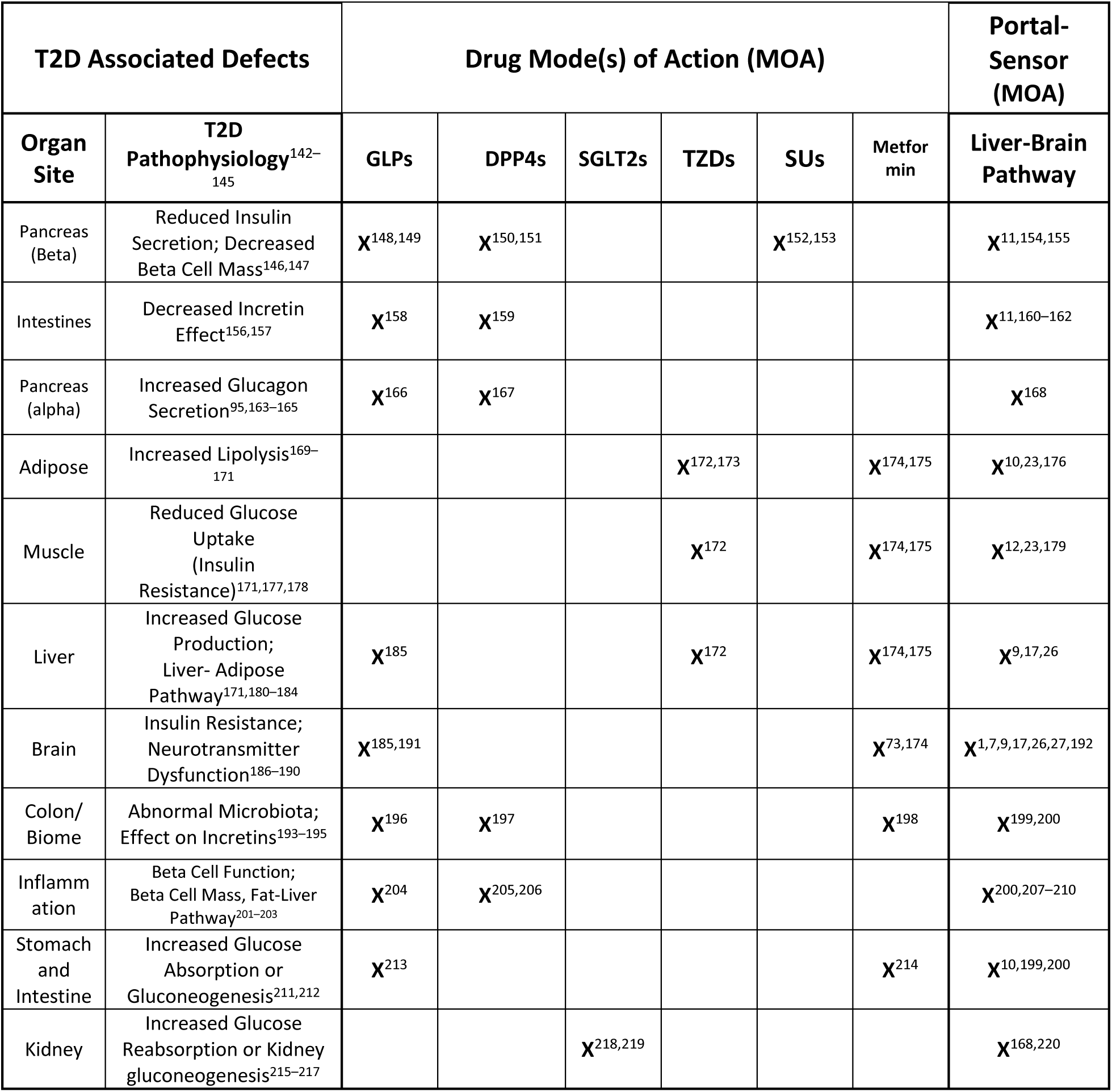
The tables contains descriptions and references on: 1) left columns: eleven of the major organ sites and organ-specific pathophysiological defects associated with Type II Diabetes (T2D), 2) middle columns: the known mechanisms of action of current T2D drugs and relation to known organ associated T2D defects, and 3) right columns: previously studied effects of the liver-brain neuropathway (including the hepatoportal glucose sensor) on metabolic function in each organ. The metabolic effects of the liver-brain neuropathway have more comprehensive coverage over the known T2D defects than any single T2D medication.

**Supplemental Figure 2.**
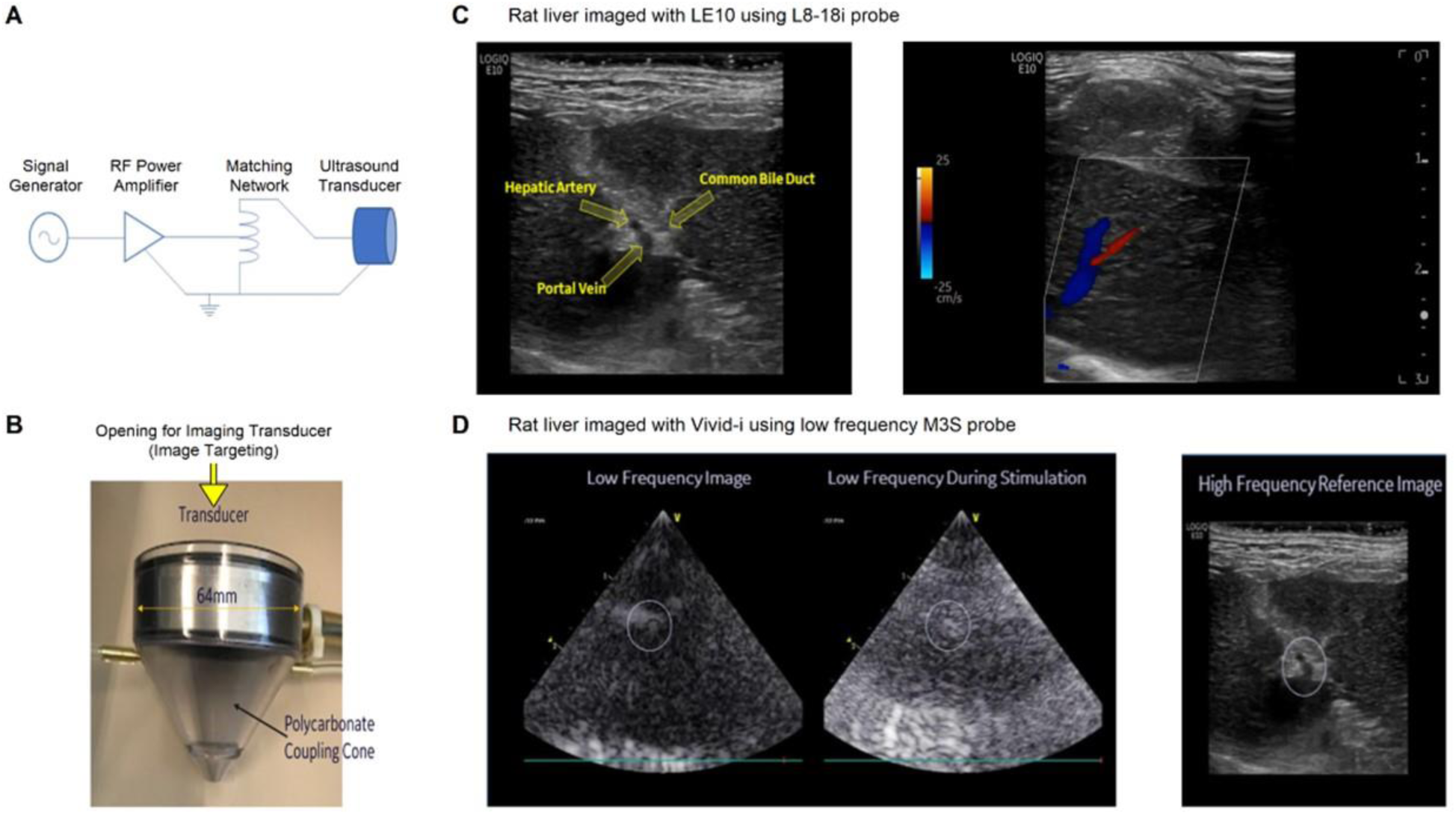
(A) Schematic of the 1.1 MHz focused ultrasound setup consisting of a signal generator (Model 33120A, Agilent Technologies Inc., Santa Clara, CA), a RF power amplifier (Model 350L, Electronics & Innovation Ltd., Rochester, NY) and a focused single-element ultrasound transducer (Model H102, Sonic Concepts Inc., Bothell, WA). The transducer is connected to the output of the power amplifier using an autotransformer matching network. (B) The 64 mm diameter ultrasound transducer has a 20 mm diameter hole in the center for insertion of an imaging probe. The transducer is acoustically coupled to the animal through a 6-cm tall plastic standoff cone filled with degassed water. (C) Ultrasound Images of rat liver using a high-frequency L8-18i linear probe (GE Healthcare, Milwaukee WI) used for identifying the targeted location. Left: B-mode image showing the outline of the liver and the dark region corresponding to the portal vein. Right: Color Doppler image showing the blood flow in the portal vein. (D) Ultrasound images of rat liver using a low frequency M3S phased-array probe (GE Healthcare, Milwaukee WI) as inserted into the opening of the focused ultrasound transducer.

**Supplemental Figure 3.**
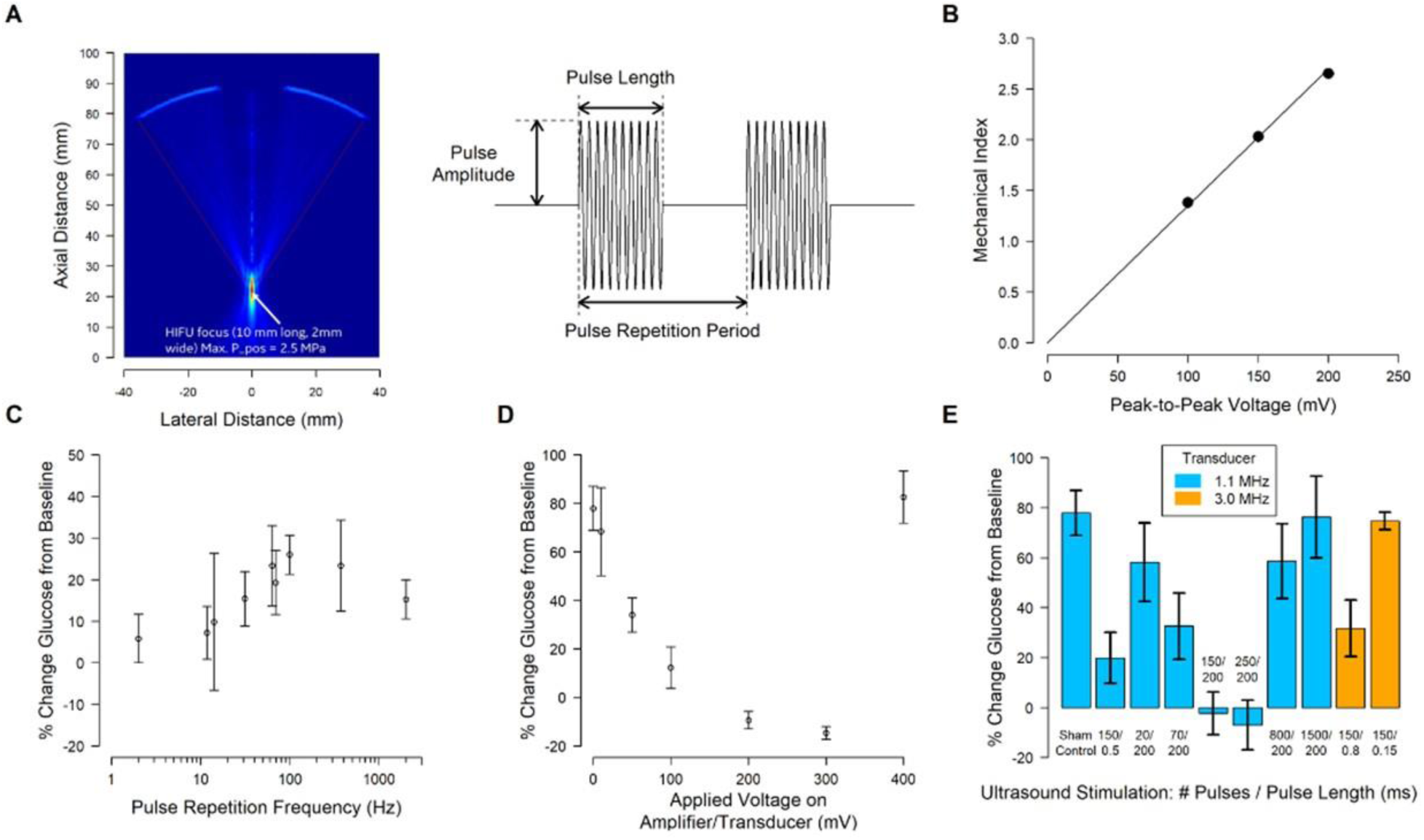
**A.** The focal pressure profile of the focused ultrasound transducer has a full-width half-maximum amplitude of 1.8 mm laterally and depth of 12 mm as simulated using Field II.^120,121^ **B.** The focused ultrasound is delivered in repeating bursts of the carrier. The waveform is governed by the selection of the Pulse Amplitude, Pulse Length and Pulse Repetition Period. The waveform is nominally delivered with a 135 mV pk-to-pk signal generator Pulse Amplitude, 150 usec Pulse Length and a 200 msec Pulse Repetition Period at the 1.1 MHz carrier frequency unless otherwise stated. The focused ultrasound transducer exhibits linearity of Mechanical Index (MI) over a range of input amplitudes up to 200 mV pk-to-pk as measured by an accredited laboratory (Acertara Acoustic Labs, Longmont, CO). **C**. Percent change from baseline in blood glucose in the LPS model of acute hyperglycemia after pFUS stimulation at various pulse repetition frequencies (Hz); at the standard pulse length, pulse amplitude, and carrier frequency used throughout the study. **D.** Percent change from baseline in blood glucose in the LPS model of acute hyperglycemia after pFUS stimulation at various applied pulse amplitudes, at the standard pulse repetition frequency, pulse length, and carrier frequency used throughout the study. **E.** Percent change from baseline in blood glucose in the LPS model of acute hyperglycemia after pFUS stimulation using various number of pulses (#) and pulse lengths (at both the standard carrier frequency (1.1. MHz) and 3.0 MHz).

**Supplemental Figure 4.**
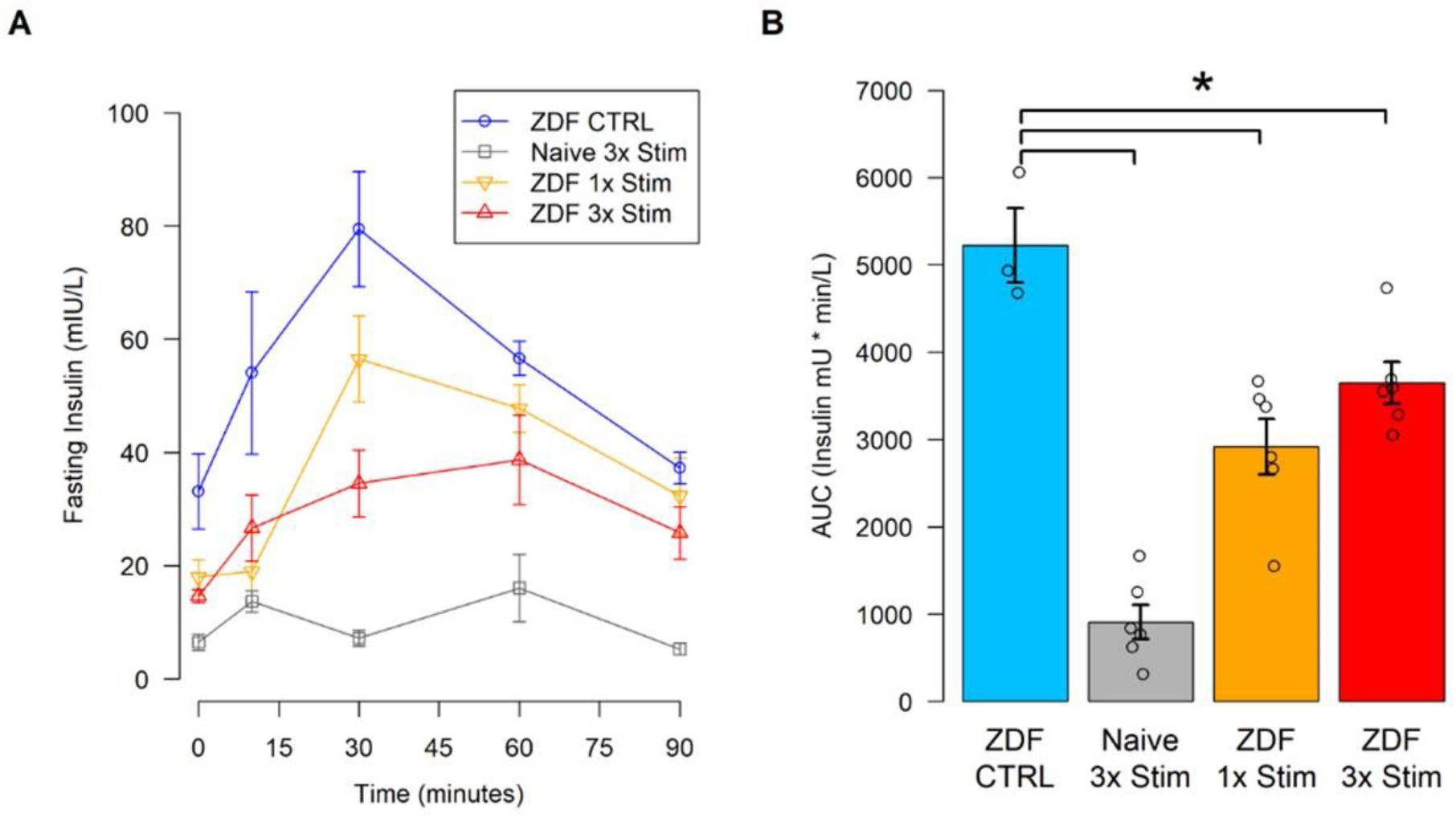
**A**. pFUS treatment resulted in a significant attenuation of hyperinsulinemia observed in ZDF rats. No significant change in effect was observed between a single application of pFUS (ZDF-1x) and three consecutive days of pFUS prior to OGTT (ZDF-3x). **B**. Calculation of incremental areas under the curve (AUC) for insulin showed a significant reduction in AUC as compared to Sham stimulated controls suggesting an increase in whole body insulin sensitivity as demonstrated by reduced hyperinsulinemia. (* p < 0.1; nonparametric Wilcoxon rank-sum test)

**Supplemental Figure 5.**
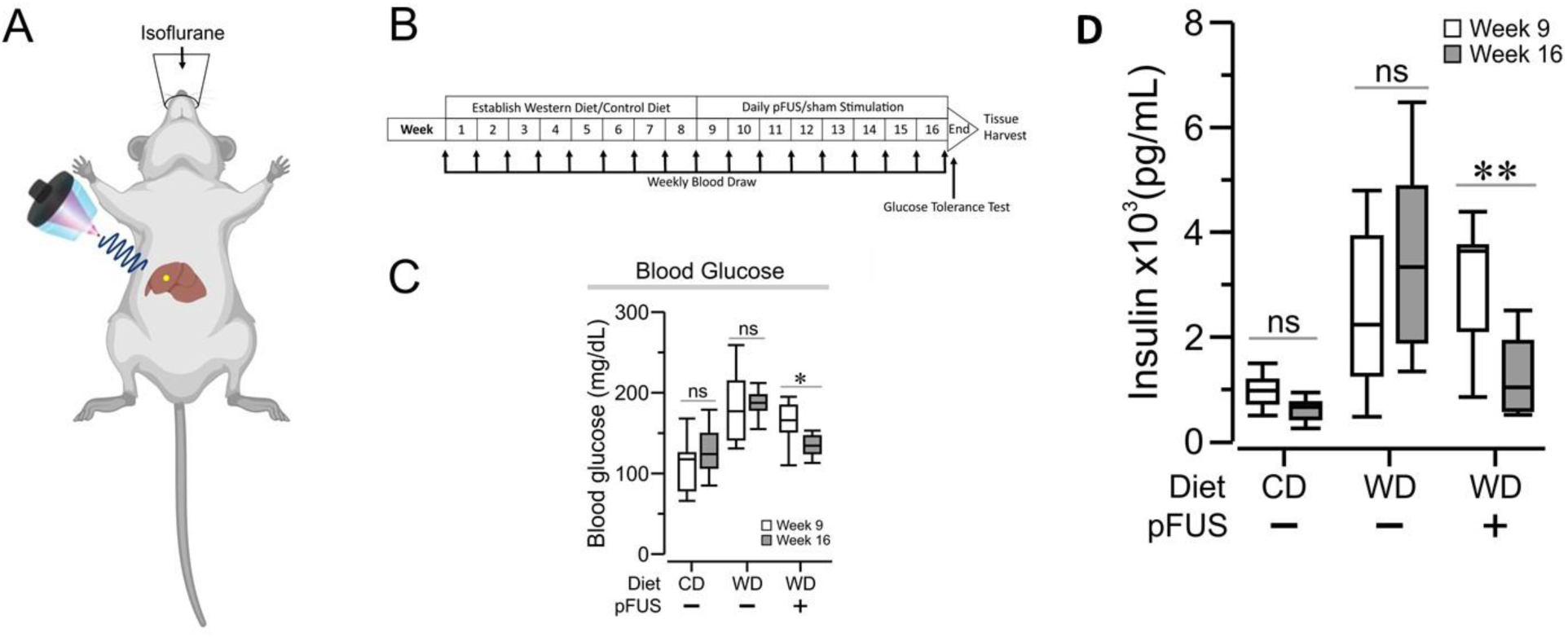
**A.** Schematic showing focused ultrasound targeting of the liver. Anesthetized mice are placed in the supine position with the conical focused ultrasound transducer aimed at the porta hepatis, represented by the yellow circle. **B.** Experimental timeline for daily stimulation. Mice are given 8 weeks to establish either the Western Diet (WD) or the control diet (CD), then are subjected to hepatic pFUS from weeks 9–16, with weekly blood draws. At the end of week 16, mice are subjected to a glucose tolerance test. **C.** Hepatic pFUS attenuates hyperglycemia in western diet-fed mice. Blood glucose levels were measured at week 9 (white plots), prior to the stimulation period and week 16 (grey plots), post-stimulation period. Daily stimulated western diet-fed mice had significantly reduced blood glucose levels across the stimulation period (WD–pFUS, week 9 vs week 16, * *P* < 0.05, 2-way ANOVA). **D.** Hepatic pFUS attenuates hyperinsulinemia in western diet-fed mice. Insulin levels were measured at week 9 (white plots), prior to the stimulation period, and at week 16 (grey plots), post-stimulation period. WD–pFUS mice had significantly reduced insulin levels across the stimulation period (WD–pFUS, week 9 vs week 16, * *P* < 0.05, 2-way ANOVA).

**Supplemental Figure 6.**
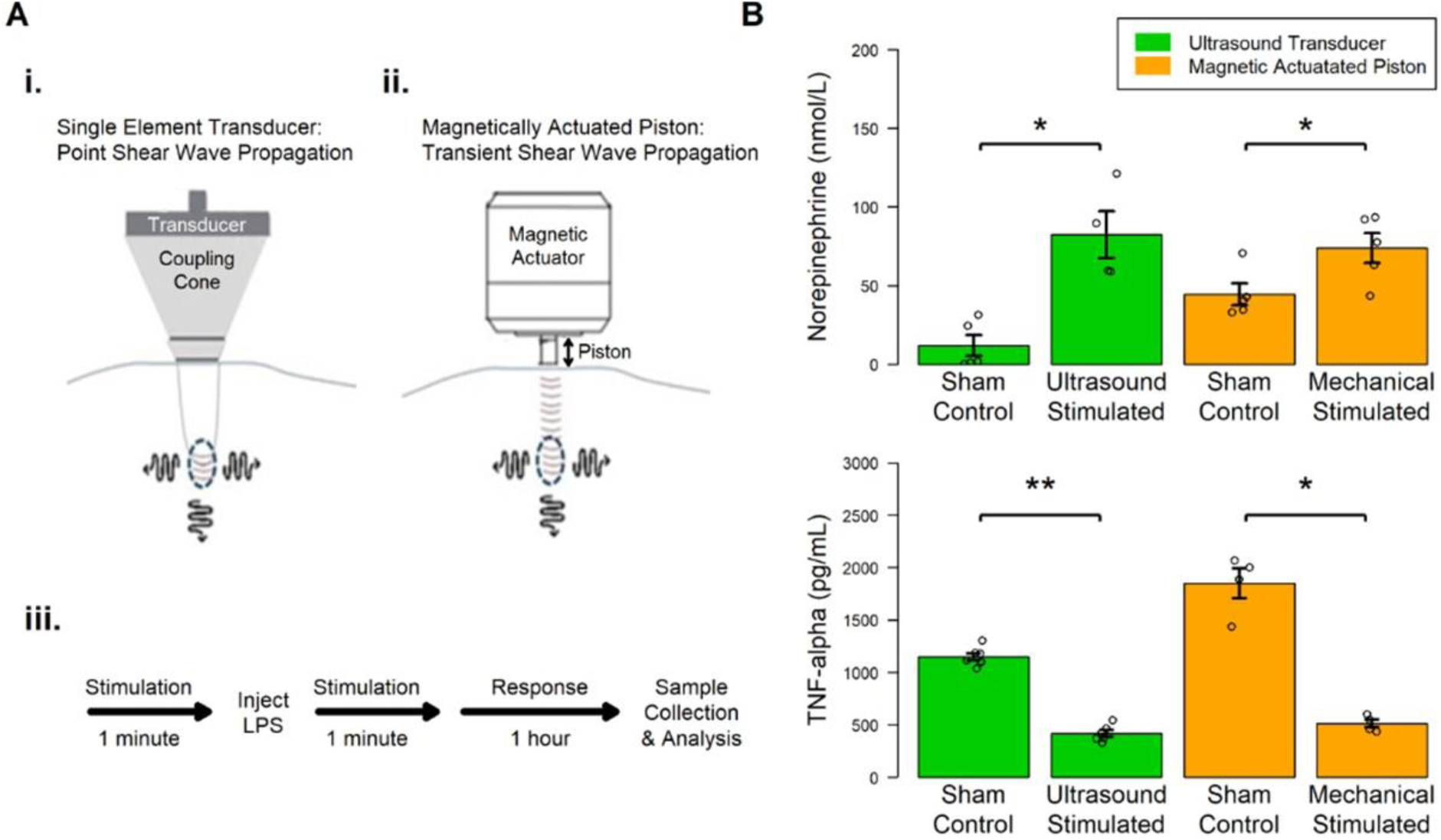
**A.** Activation and neuromodulation of the splenic cholinergic anti-inflammatory pathway (CAP)^1^ using both an ultrasound and a mechanical piston-based transducer. **i.** A schematic of the ultrasound transducer described previously^1^ (and its acoustic coupling to the animal using the water-filled coupling cone). As described above, the transducer provides an ultrasound-based stimulus with a mechanical component based on point shear wave propagation at the focal point of the ultrasound stimuli.^1,221,222^ **ii.** A schematic of the non-ultrasonic, magnetically actuated piston described above to provide a mechanical stimulus when directly coupled to the animal (based on 1D transient shear wave propagation from the piston-tissue interface).^222^ **iii.** The timeline of ultrasonic or mechanical neuromodulation experiments performed in the LPS-induced inflammation^1^ model. **B. (left)** Concentration of splenic norepinephrine (NE, top) and TNF alpha (bottom) with and without ultrasound stimulation in LPS-treated animals versus sham controls. **(bottom)** Concentration of splenic norepinephrine (NE, top) and TNF alpha (bottom) with and without mechanical-piston based stimulation in LPS-treated animals versus sham controls. In evaluating CAP response to the mechanical-piston stimulus to ultrasound-based stimulation (as measured by CAP-related neurotransmitter (NE) and cytokine (TNF alpha) concentrations)^1^, the mechanical piston-based stimulation resulted in nearly identical elevation of NE and reduction of splenic TNF in response to the piston versus ultrasound transducer-based stimulus. *(* p < 0.1; ** p<0.01; nonparametric Wilcoxon rank-sum test)*

**Fig. S7. A-B.**
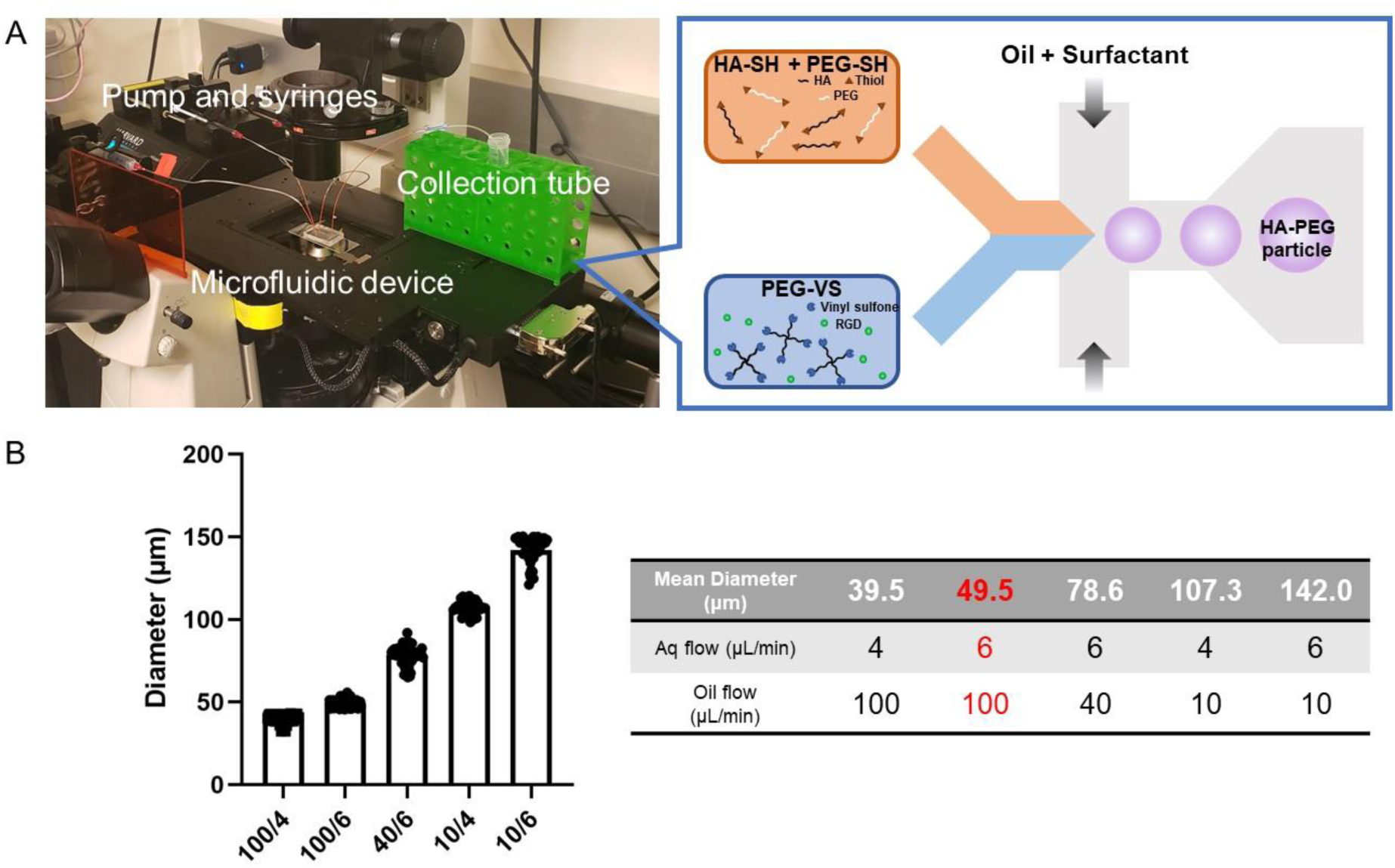
**A**. Photograph of the setup for microparticle fabrication and scheme illustrating microgel formation that is utilized to provide an optimal growth environment for three dimensional culture ^31^ of neurons (see figure 2 in the main text) using a microfluidic water-in-oil emulsion system. A pre-gel and crosslinker solution are segmented into monodisperse droplets followed by in-droplet mixing and crosslinking via Michael addition. **B**. The operational regime for microfluidic microgel generation has a large dynamic range, spanning almost an order of magnitude in size while maintaining tight control at each condition, with CVs<7% in all cases. We use 50-100 µm microparticles for experiment as this provided the best pore size for neuron culture.

**Fig. S7. C-F.**
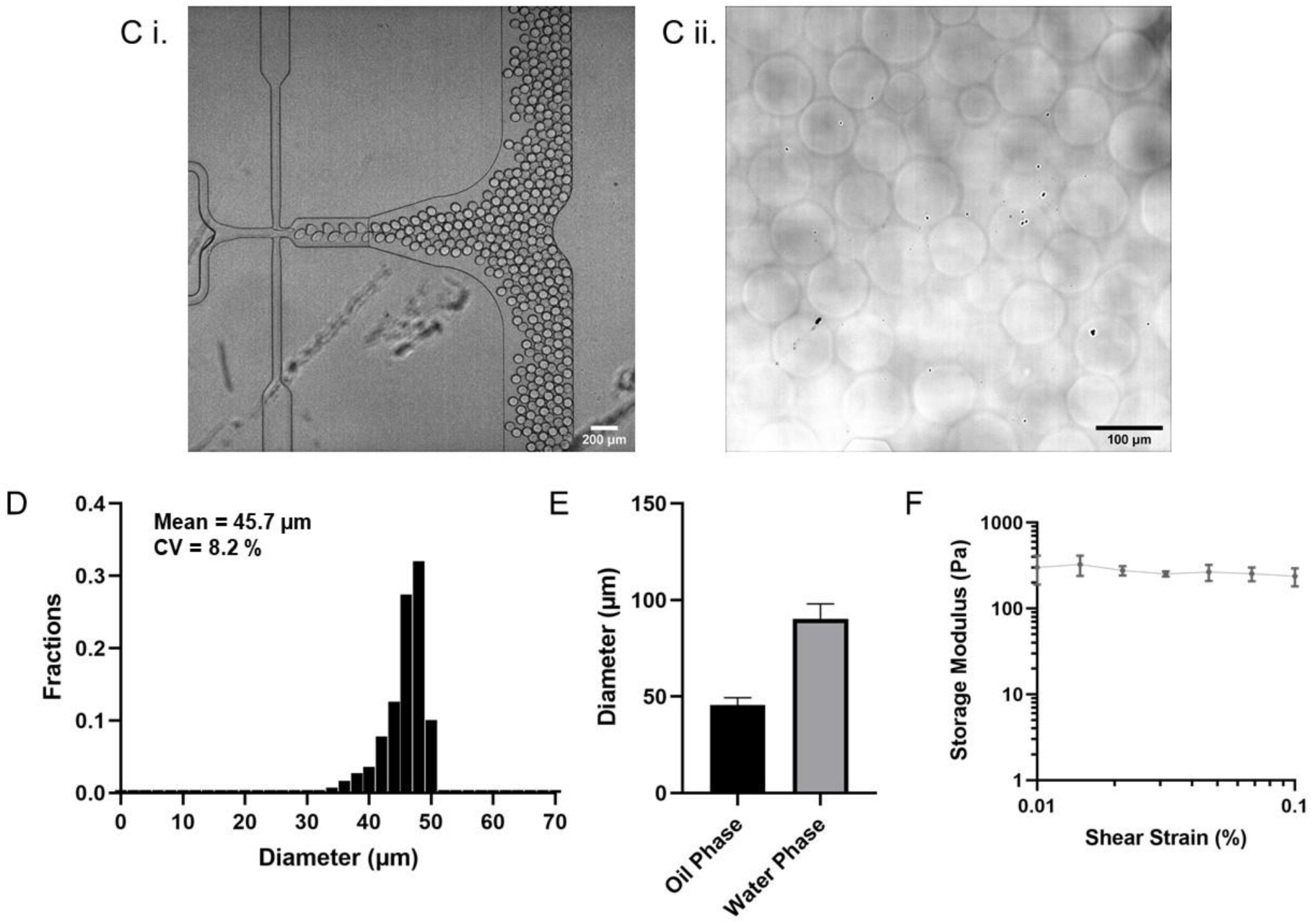
**C i.** Representative images of microgel droplets in flow after generation, which are the utilized to fabricate the three dimensional neuron culture^31^ (see figure 2 in the main text). **C ii.** Representative image of cross-linked microgels after phase transfer. **D.** Generation of HA microgel particles with highly defined sizes. **E.** HA microgel particles, made with a 50 µm diameter when surrounded by oil swell in buffer after aqueous extraction from the oil phase. **F.** Storage moduli of hydrogels. Data are shown as mean ± SD. HA microgel particles were manufactured with an ideal storage modulus tuned for neuron culture (< 500 Pa)

**Fig. S7. G-I.**
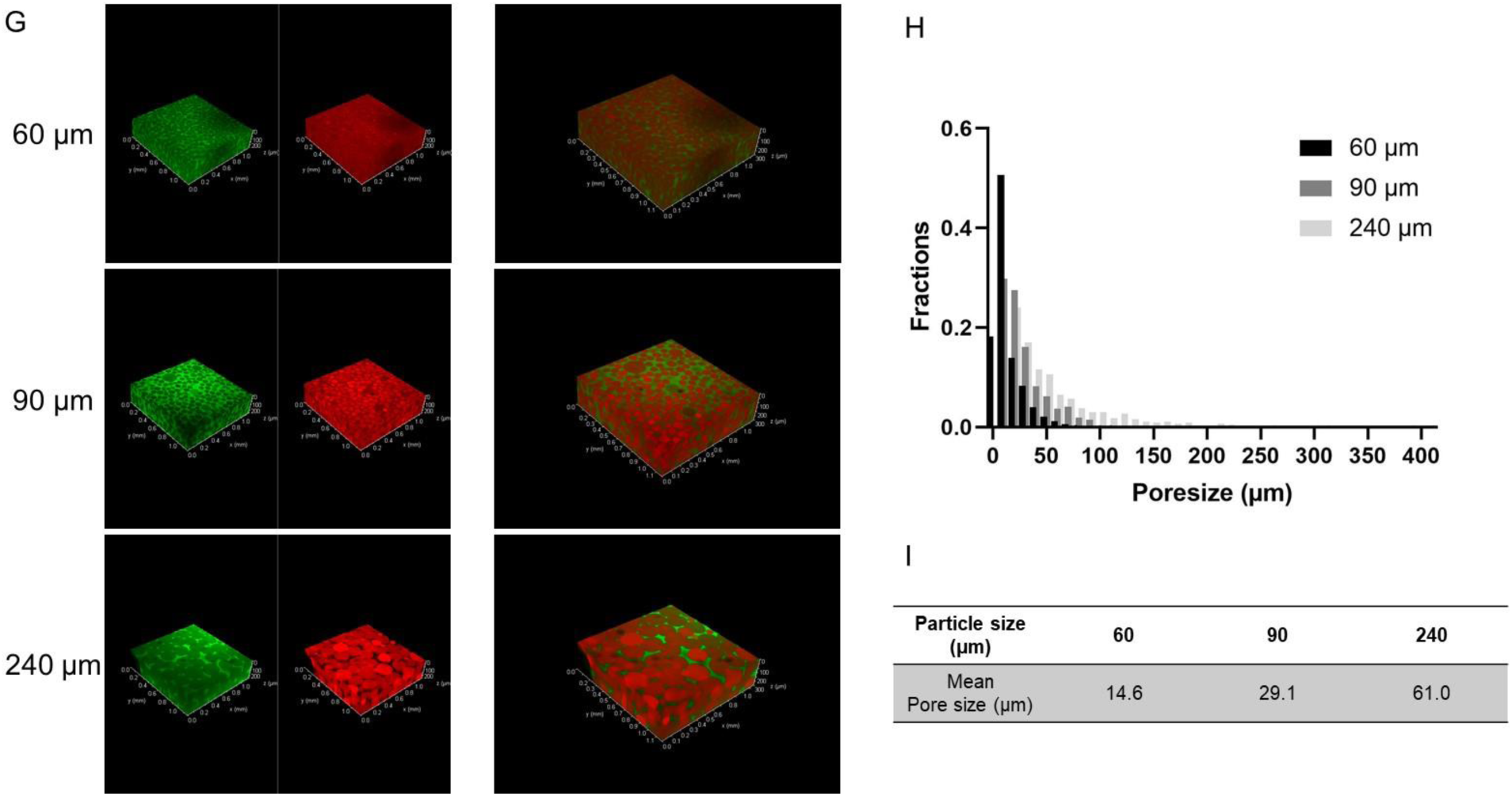
**G.** Confocal reconstructions of HA microgel-based neuron culture scaffolds^31^ after incubation with high molecular weight (Mw = 500 kDa) FITC-dextran indicates interconnectivity of the void spaces through the scaffold. In all cases, FITC dextran is represented in green and HA microparticles are represented in red. Different size of particles yield different void spaces. **H. and I.** Different building-block sizes allow deterministic control over resultant microporous network characteristics. 30 µm, 50 µm and 100 µm diameter microgel particles yield 15 µm, 25 µm, and 60 µm mean pore size respectively. Since dissociated neurons are around 10 ∼ 30 µm, we chose 50-100 µm microgel particles for further cell culture experiments.

**Fig. S7. J.**
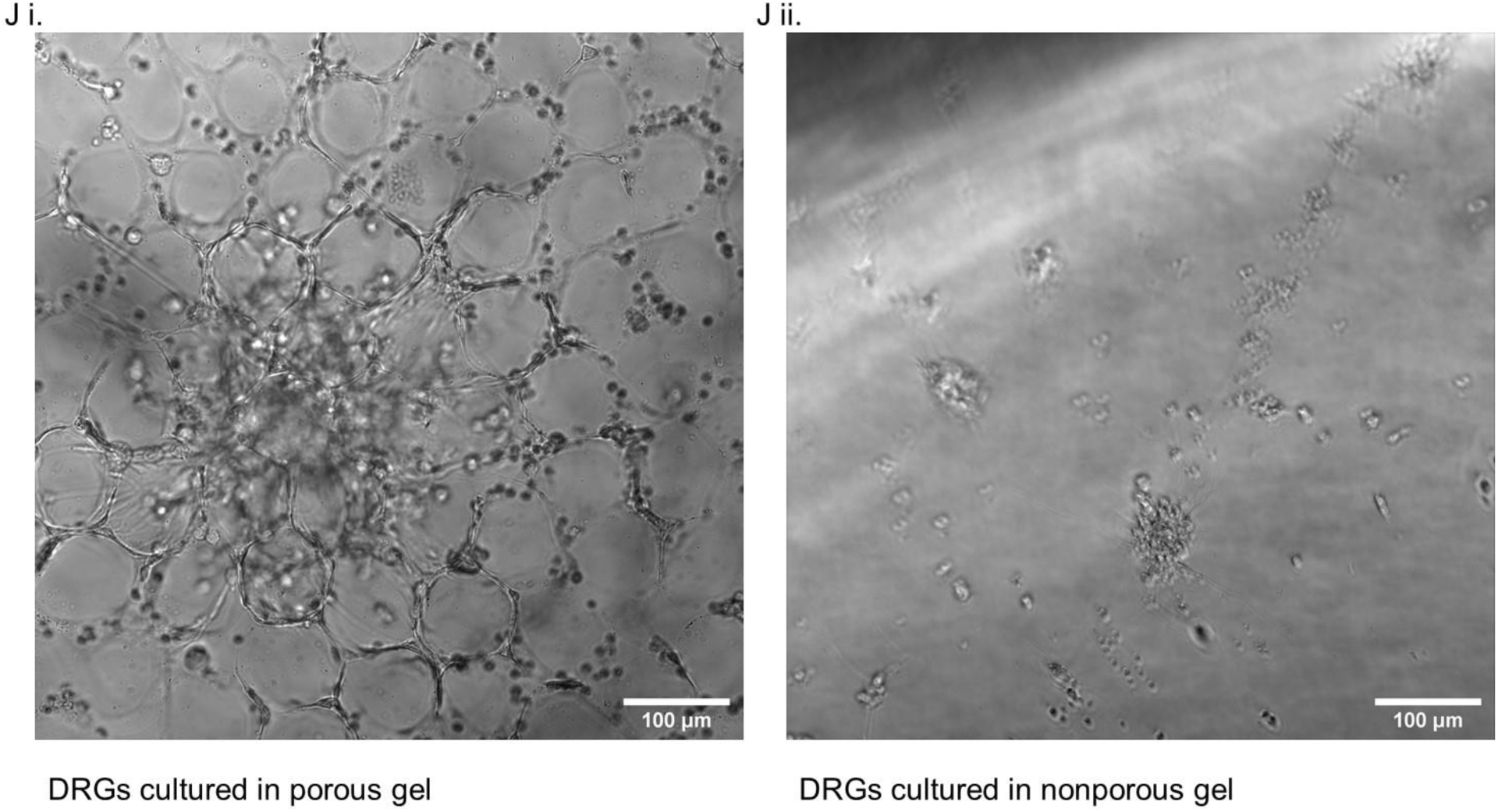
**J i.** Bright field image of 3D culture of neuron cells within an HA-based microporous gel^31^ (see figure 2 in the main text)**. J ii.** Bright field image of 3D culture of neuron cells within a HA-based nanoporous gel (i.e., cell directly embedded in the gel).

**Supplemental Figure 8.**
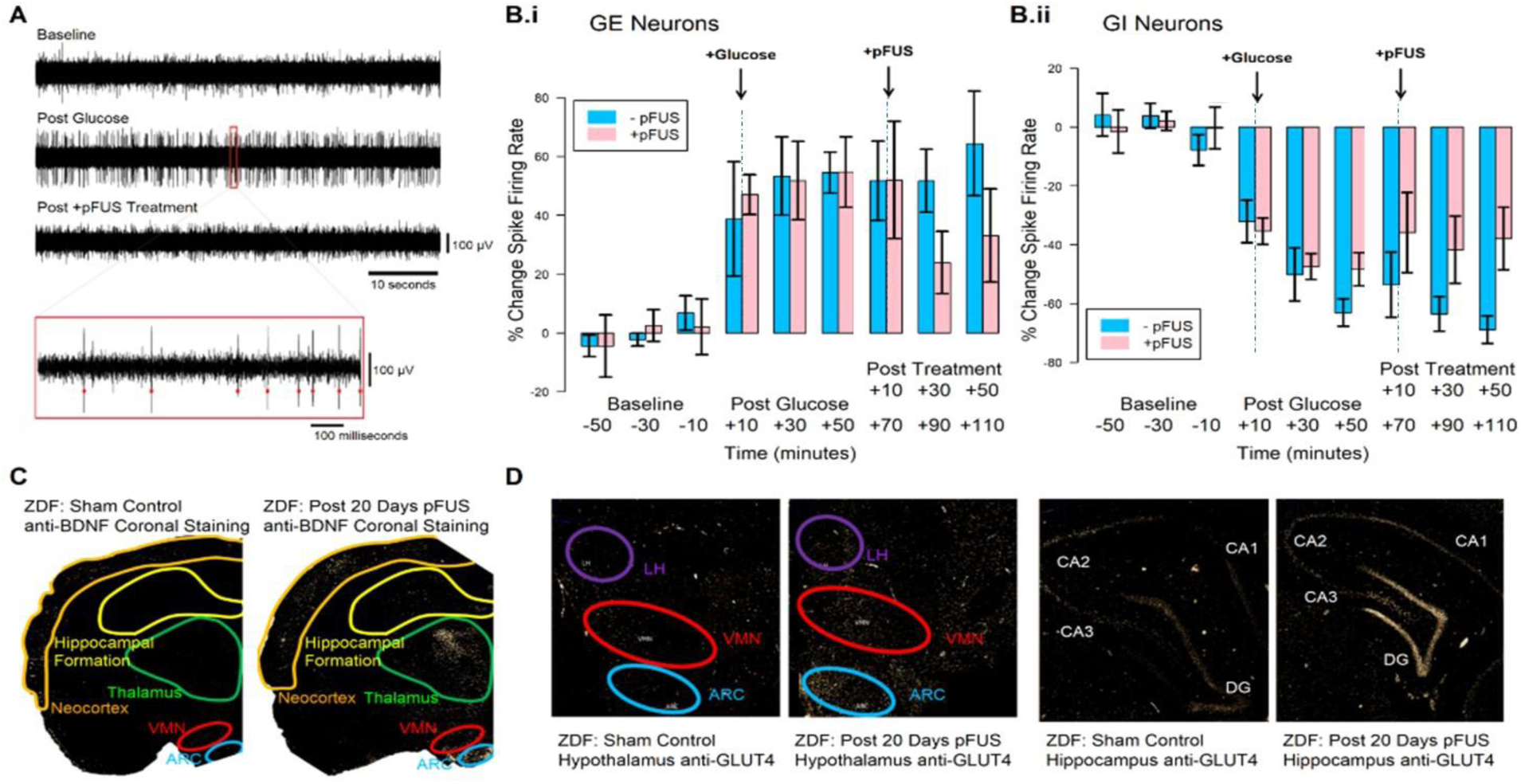
**A**. Sample electrode recordings in the PVN for data shown in Figure 3B ii. Sample traces show 60s of recording in Baseline, Glucose+ and pFUS+ periods. Zoomed trace shows 1s of recording in the Glucose+ period with detected spikes indicated by red dots. **B.** Firing rates for GE (**i.**) and GI (**ii.**) for all time points before (−50 – 0) glucose infusion, and before and after pFUS. Data in figure 3C represents firing rates for GE and GI neurons following with and without pFUS at the +30-minute post-pFUS timepoint. **C.** Histochemical analysis of paraffin-embedded rat brain tissue labeling BDNF antibody showing the unstimulated control (left) and pFUS stimulated animals (right). As observed a significant increase in BDNF staining was visible in the hypothalamus (with prominence in the arcuate and ventromedial hypothalamus), thalamic and hippocampal brain regions. Images are included as partial coronal sections to demonstrate total BDNF activation by pFUS across a number of brain regions. **D.** Histochemical analysis of paraffin embedded rat brain tissue labeling GLUT-4 receptor antibody showing hypothalamic (top) and hippocampal (bottom) staining patterns. As observed a significant increase in both hypothalamic and hippocampal GLUT-4 translocation occurred following hepatic pFUS (top and bottom right) as compared to unstimulated controls (top and bottom left).

**Supplemental Figure 9.**
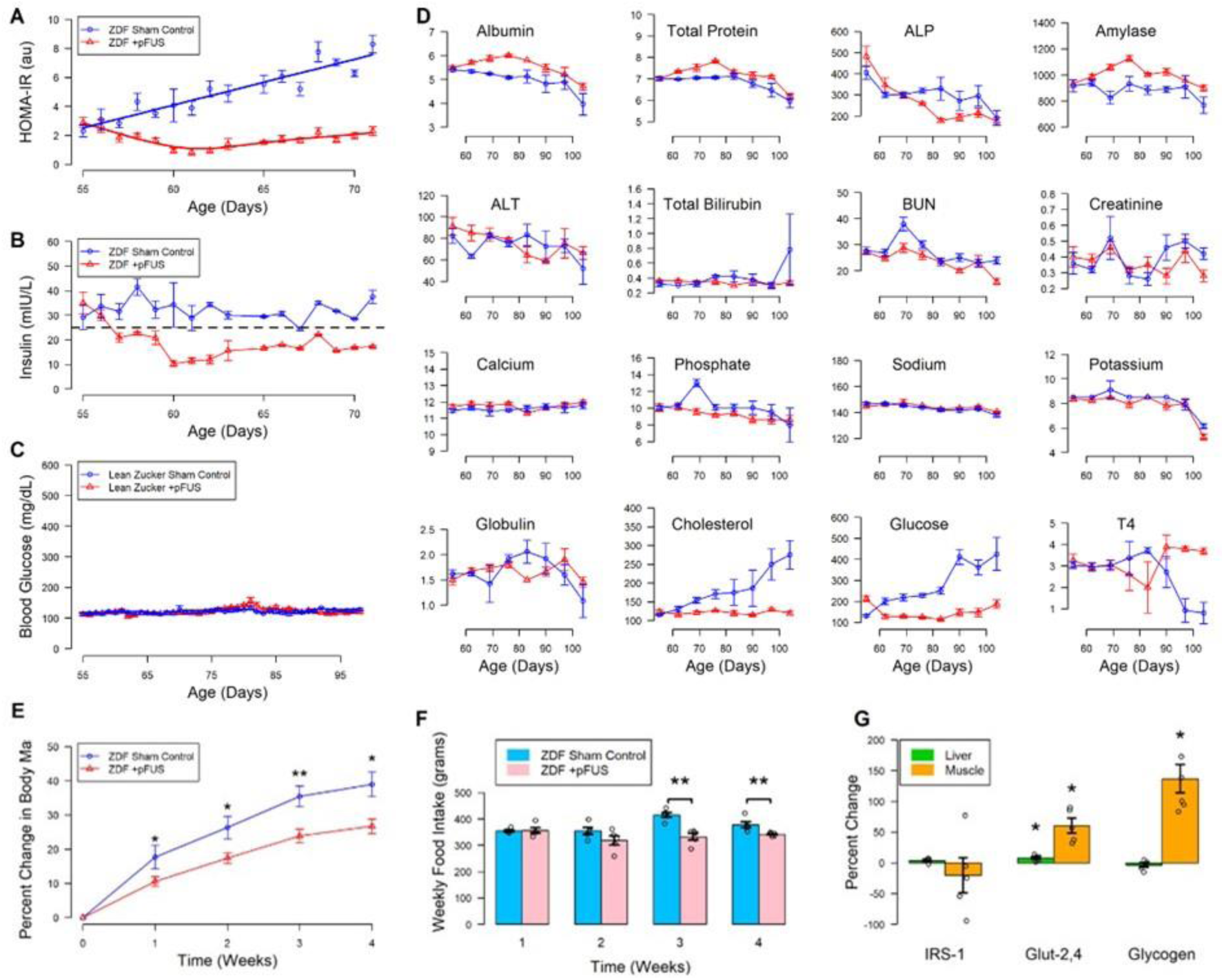
**A.** Chronic pFUS treatment applied beginning in the prediabetic stage (day 55) and continued for 2 weeks (through day 75) significantly improved insulin sensitivity as determined by HOMA-IR scoring in a ZDF rat model of T2DM. **B.** Chronic pFUS was associated with a rapid decrease in circulating insulin concentrations indicating improved insulin sensitivity following chronic pFUS treatment in the ZDF rat. **C.** pFUS treatment did not change circulating glucose levels as compared to Sham-CTRL in a naïve, non-diabetic animal model (SD rat). **D.** Comprehensive metabolic analysis of circulating blood markers in the ZDF rat model collected on a weekly basis for 7 weeks not only showed no significant change in markers associated with liver or renal disease but also failed to show a change in circulating electrolytes, demonstrating the lack of tissue damage following chronic pFUS. Changes in cholesterol, glucose and the inactive form of thyroid hormone support an improvement in obesity and T2DM related co-morbidities, including metabolic syndrome and hypercholesteremia. **E.** A significant slowing of the overall rate of weight gain in pFUS treated (orange line) ZDF rats as compared to their sham stimulated littermates (blue line) was observed. **F**. This decrease in both rate and total weight gain is unlikely the result of decreased food intake alone with no significant change in food intake occurring between weeks 1-2 of treatment. A small decrease in overall food intake was observed in Week 3 of treatment in those animals receiving daily pFUS treatment. **G.** Effects of daily pFUS stimulation on hepatic and skeletal muscle in the genetic model of T2D. The percent change in the concentration of IRS-1, GLUT2, GLUT4 and glycogen in hepatic and skeletal muscle from a ZDF rat following chronic stimulation between stimulated (n=5) and sham control (n=5) ZDF rats following chronic treatment (20days of treatment). (* p < 0.1, ** p < 0.01; nonparametric Wilcoxon rank-sum test)

**Supplemental Figure 10:**
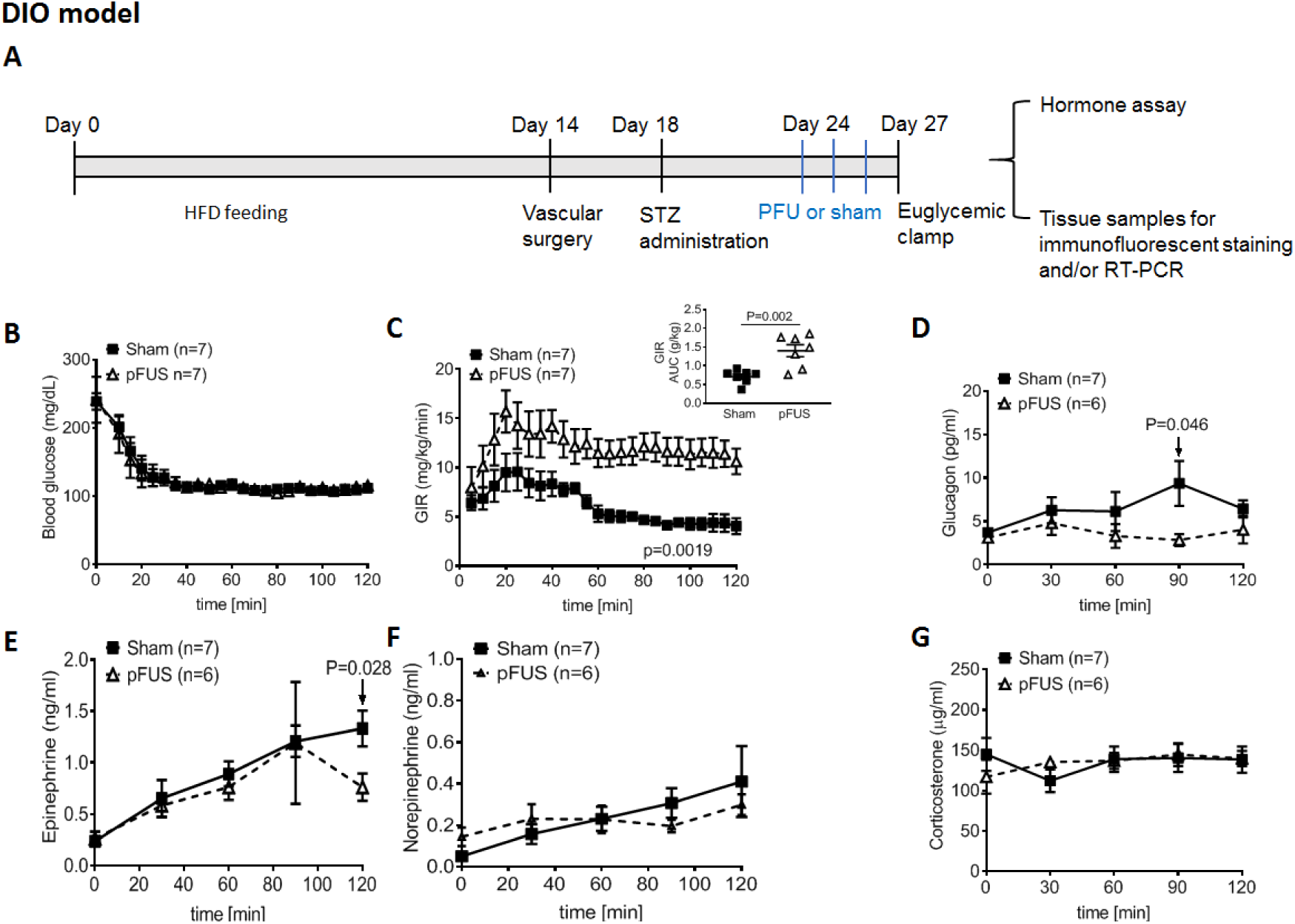
Effect of pulsed focused ultrasound (pFUS) of the porta hepatis on insulin sensitivity in STZ-induced diabetic diet-induced obesity (DIO) rats, as quantified through hyperinsulinemic-euglycemic clamp (HEC; see material and methods for experimental description). **A)** Timeline of experimental interventions. **B)** Glucose infusion rate (GIR) during standardized hyperinsulinemic clamp revealing higher steady state glucose infusion requirement after pFUS treatment and **C)** glucose infusion rate area under the curve (AUC) for entire study. Plasma hormone change during the clamp including **D)** glucagon **E)** epinephrine, **F)** norepinephrine and **G)** corticosterone. Values are mean ± SEM; 2-way ANOVA (GIR); multiple t-tests (hormones) student t-test (AUC)

**Supplemental Figure 11:**
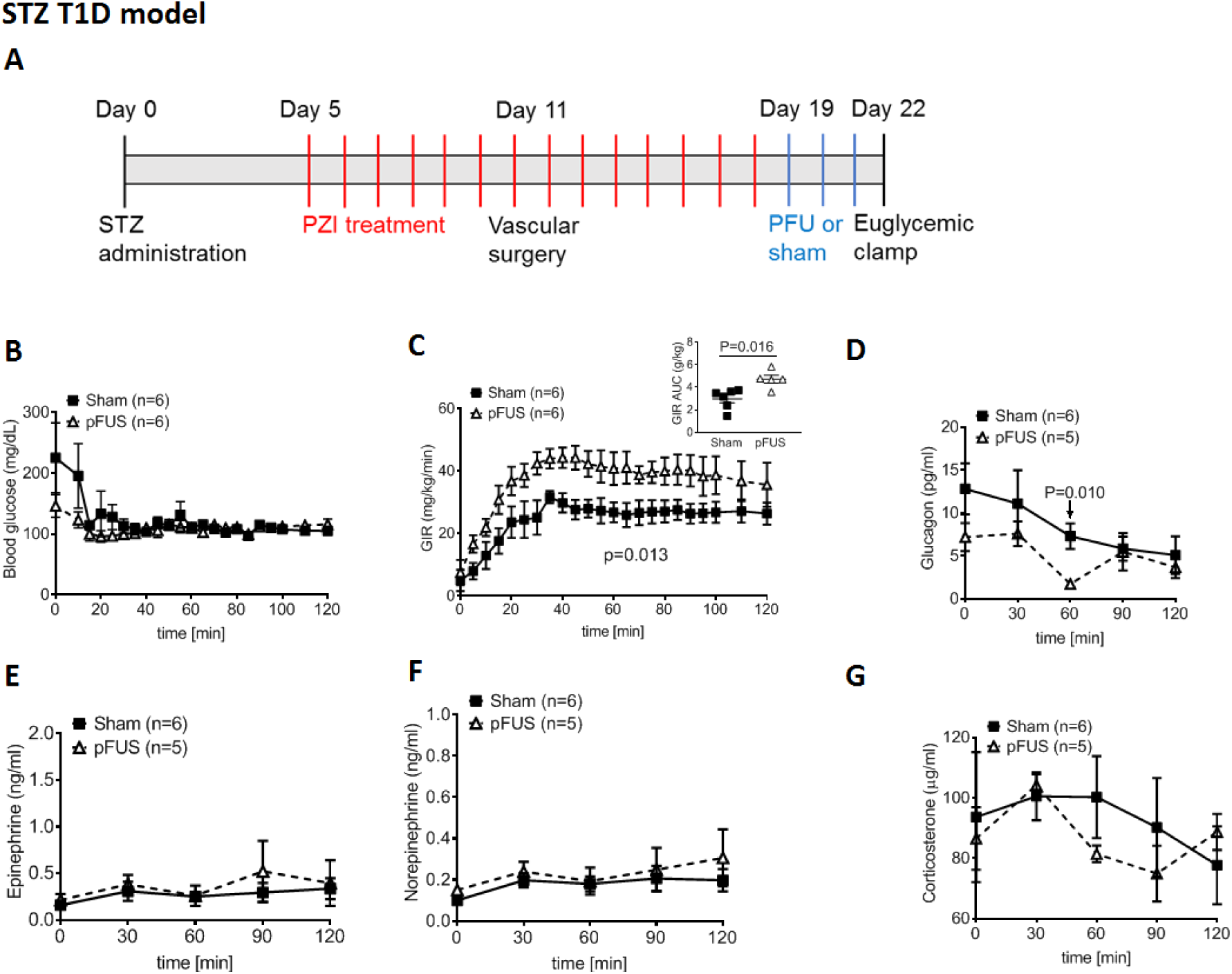
Effect of pulsed focused ultrasound (pFUS) of the porta hepatis on insulin sensitivity in STZ-induced T1D rats, as quantified through hyperinsulinemic-euglycmic clamp (HEC). **A**) Timeline of experimental interventions (see material and methods for experimental description). **B**) Glucose infusion rate (GIR) during standardized hyperinsulinemic clamp revealing higher steady state glucose infusion requirement after pFUS treatment and glucose infusion rate area under the curve (AUC) for entire study. Plasma hormone change during the clamp including **D**) glucagon, **E**) epinephrine, **F**) norepinephrine and **G**) corticosterone. Values are mean ± SEM.; Values are mean ± SEM; 2-way ANOVA (GIR); multiple t-tests (hormones) student t-test (AUC).

**Supplemental Figure 12.**
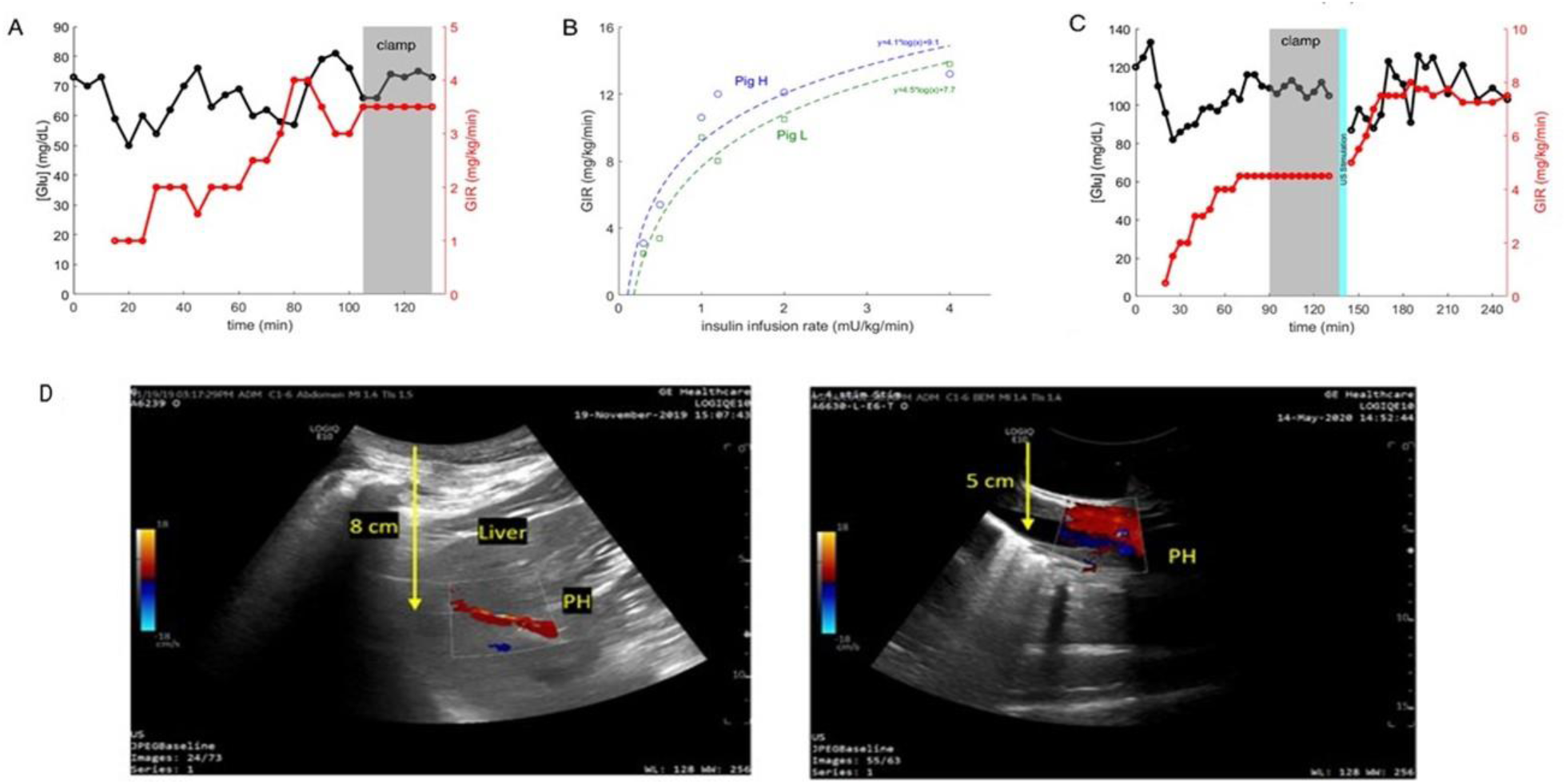
Short-term effect of focused ultrasound (pFUS) of the porta hepatis on insulin sensitivity in healthy swine, as quantified through hyperinsulinemic-euglycmic clamp (HEC; see material and methods for experimental description). **A**. Representative data from an HEC experiment. Under continuous, constant infusion of insulin at a rate of 0.5 mU/kg/min, glucose infusion rate (GIR) was adjusted every 5 minutes according to a formula^223^ to achieve euglycemia (defined as baseline glucose concentration +/−10%) and maintained for ≥ 30 min (grey shaded area). In this example, GIR at euglycemic equilibrium was 3.5 mg/kg/min. The quality of the HEC was assessed by calculating the normalized coefficient of variation (CV) of the glucose concentration and GIR for the duration of euglycemic equilibrium. In this experiment, CV for glucose was 6.92% and for GIR was 6.51%. **B**. In step-clamp experiments, euglycemic equilibrium GIR values were calculated for different insulin infusion rates (IIR) in 2 animals. Based on this data, we decided to conduct FUS experiments at IIR of 0.5 mU/kg/min, as the slope of the GIR curves, and therefore the sensitivity of the method to resolve changes in insulin sensitivity, decreased at higher IIRs. **C**. Example of a pFUS experiment, after establishing HEC (CV for glucose 2.85%, for GIR 0%). pFUS of the porta hepatis was applied for 4 minutes (blue shaded area), after which GIR was adjusted, just like before, to maintain euglycemia. Increase in GIR after pFUS reflects increased insulin sensitivity. **D.** Imaging of the porta hepatis in swine. Left: Noninvasive imaging (probe placed on skin). Right: Invasive imaging (probe placed on top of PH after it was surgically accessed).

**Supplemental Figure 13.**
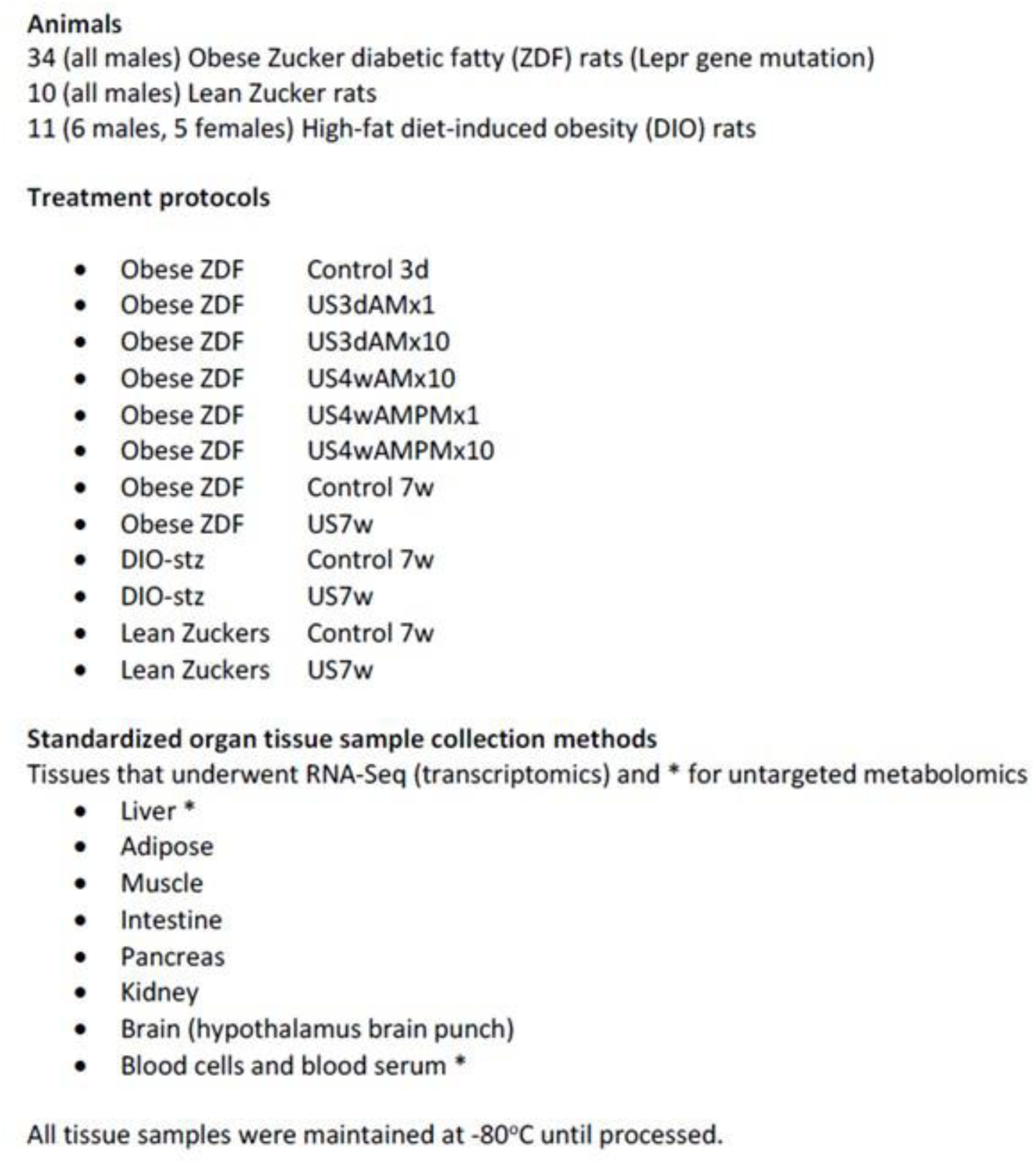
Sample list and description of metabolomic and transcriptomic samples; including controls.

**Supplemental Figure 14.**
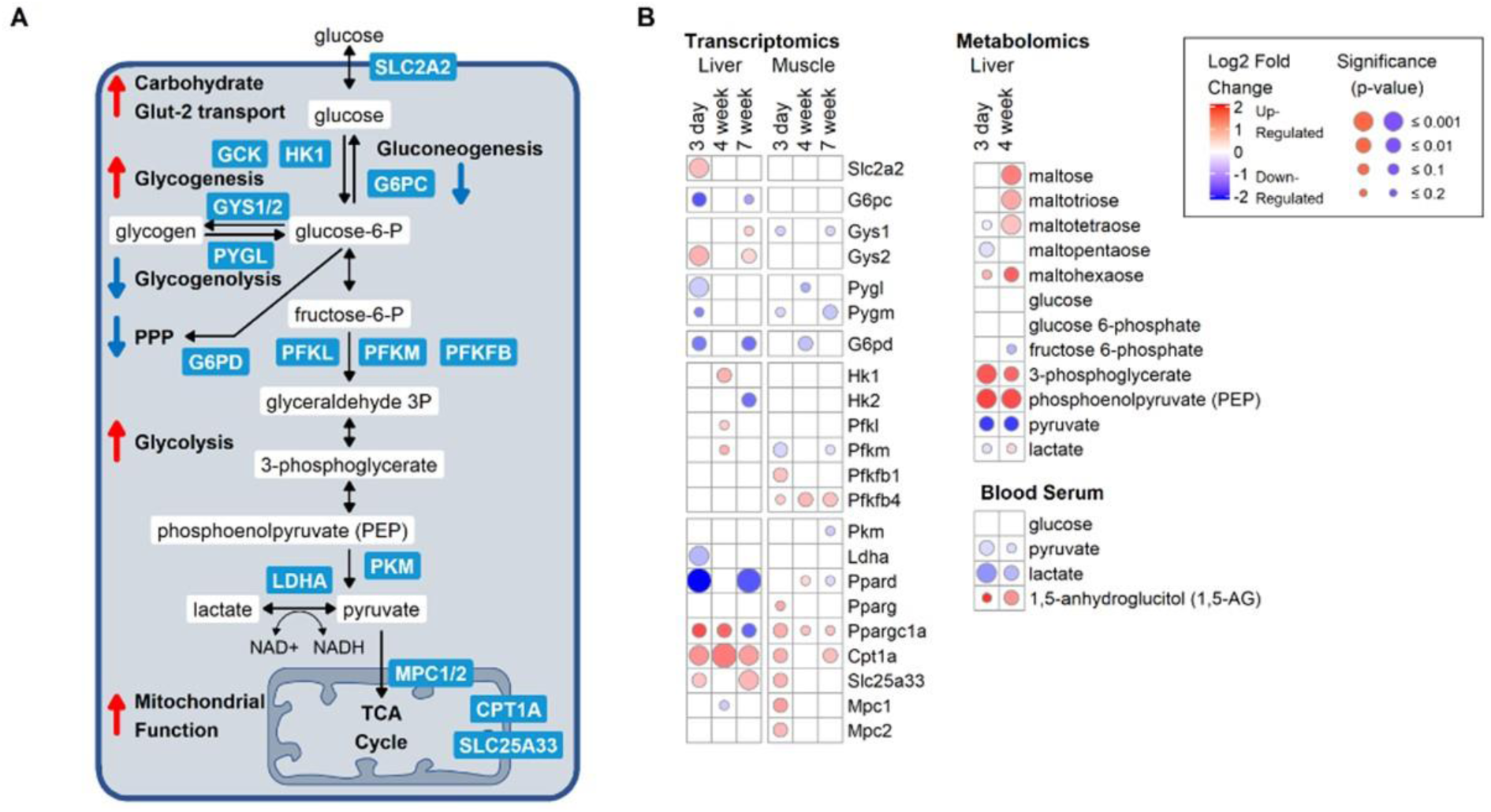
**A.** Schematic of key protein enzymes (blue rectangles), metabolites (white rectangles) and metabolic pathways (e.g. gluconeogenesis, fatty acid and bile acid metabolism, oxidative phosphorylation, mitochondrial transport) that are either upregulated (red upward arrows) or downregulated (blue downward arrows) within the liver-muscle axis. **B. (left)** Differential RNA transcripts (i.e. genes) from liver and muscle tissue of ZDF animals undergoing 3 days, 4 weeks, or 7 weeks of daily pFUS stimulation vs sham controls. The protein encoded by these differentially expressed genes is presented in the schematic diagram. **(right)** The changes in metabolites in liver tissue at 3 days and 4 weeks between ultrasound treated vs controls. The size of each circle represents the p-value of differentiation between the pFUS stimulated group (n=5) vs sham controls (n = 5). The color of the circle represents the degree of being upregulated (red) or downregulated (blue).

**Supplemental Figure 15.**
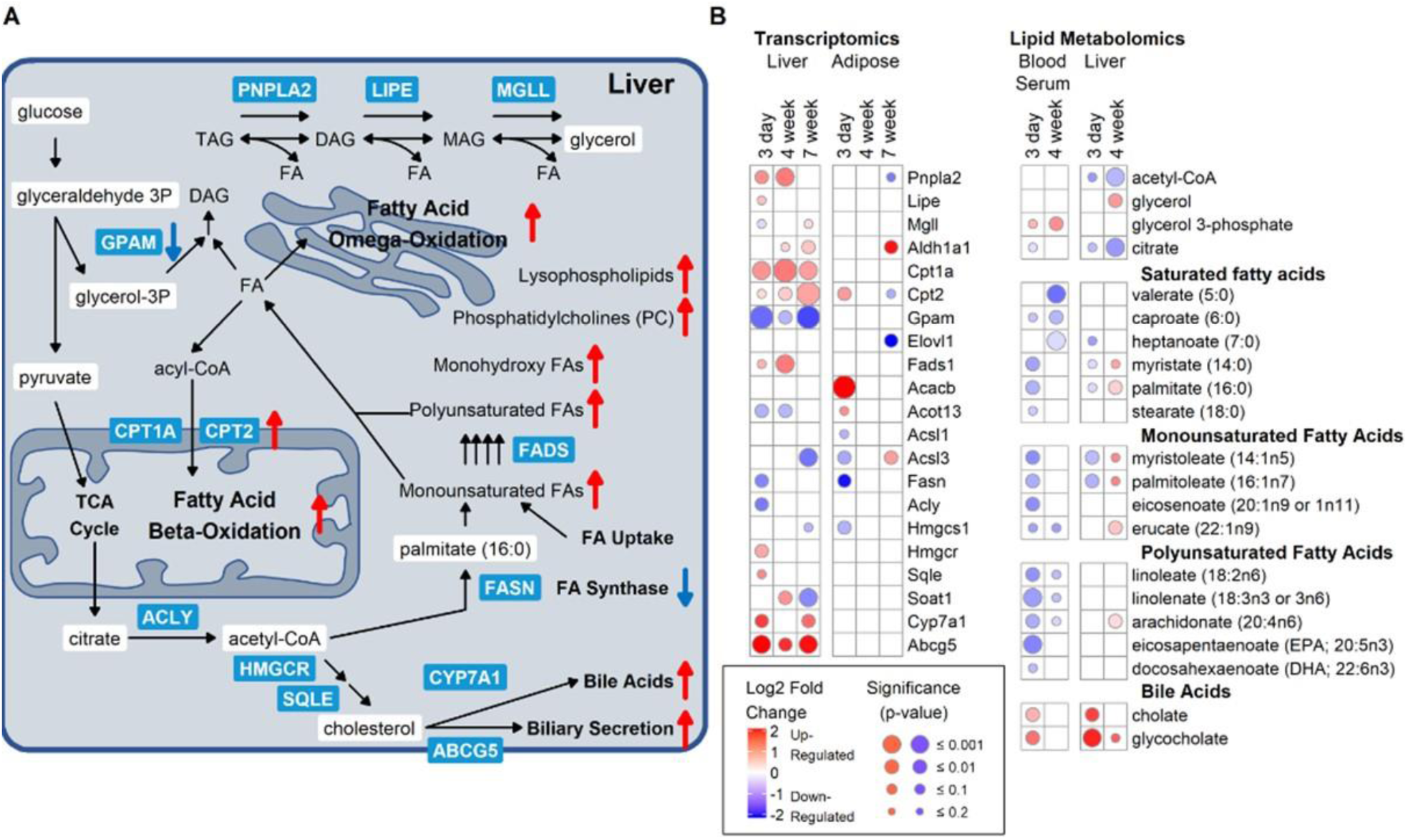
**A.** Schematic of key protein enzymes (blue rectangles), metabolites (white rectangles) and metabolic pathways (e.g. gluconeogenesis, fatty acid and bile acid metabolism, oxidative phosphorylation, mitochondrial transport) that are either upregulated (red upward arrows) or downregulated (blue downward arrows) within the liver-adipose axis. **B. (left)** Differential RNA transcripts (i.e. genes) from liver and adipose tissue of ZDF animals undergoing 3 days, 4 weeks, or 7 weeks of daily pFUS stimulation vs sham controls. The protein encoded by these differentially expressed genes is presented in the schematic diagram. **(right)** The changes in metabolites in liver tissue at 3 days and 4 weeks between ultrasound treated vs controls. The size of each circle represents the p-value of differentiation between the pFUS stimulated group (n=5) vs sham controls (n = 5). The color of the circle represents the degree of being upregulated (red) or downregulated (blue).

**Supplemental Figure 16.**
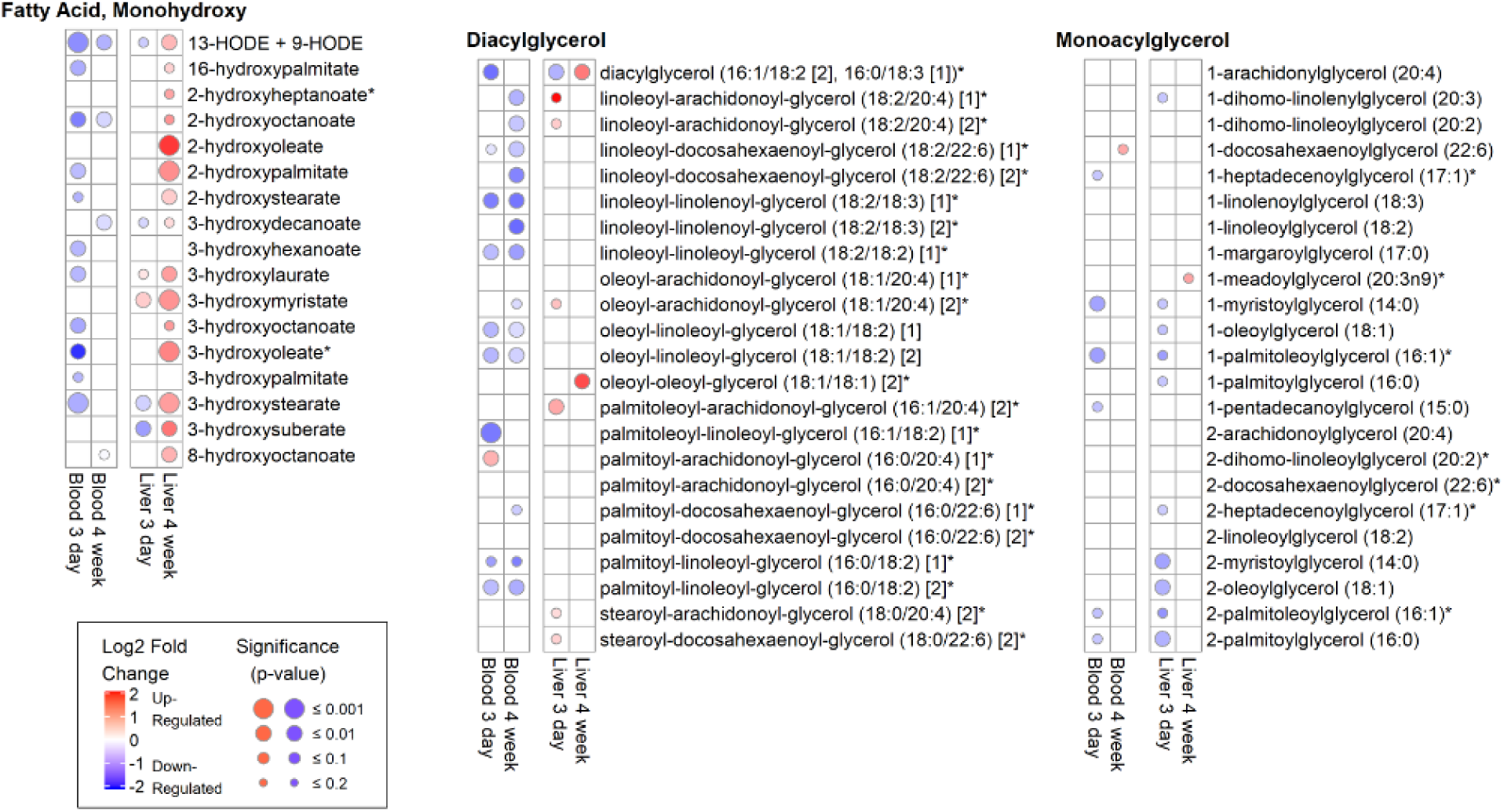
Changes in monohydroxy fatty acids, Diacylgycerols (DAGs), Monoacylglycerols (MAGs) in blood serum and liver tissue at 3 days and 4 weeks between ultrasound treated vs control ZDF animals.

**Supplemental Figure 17.**
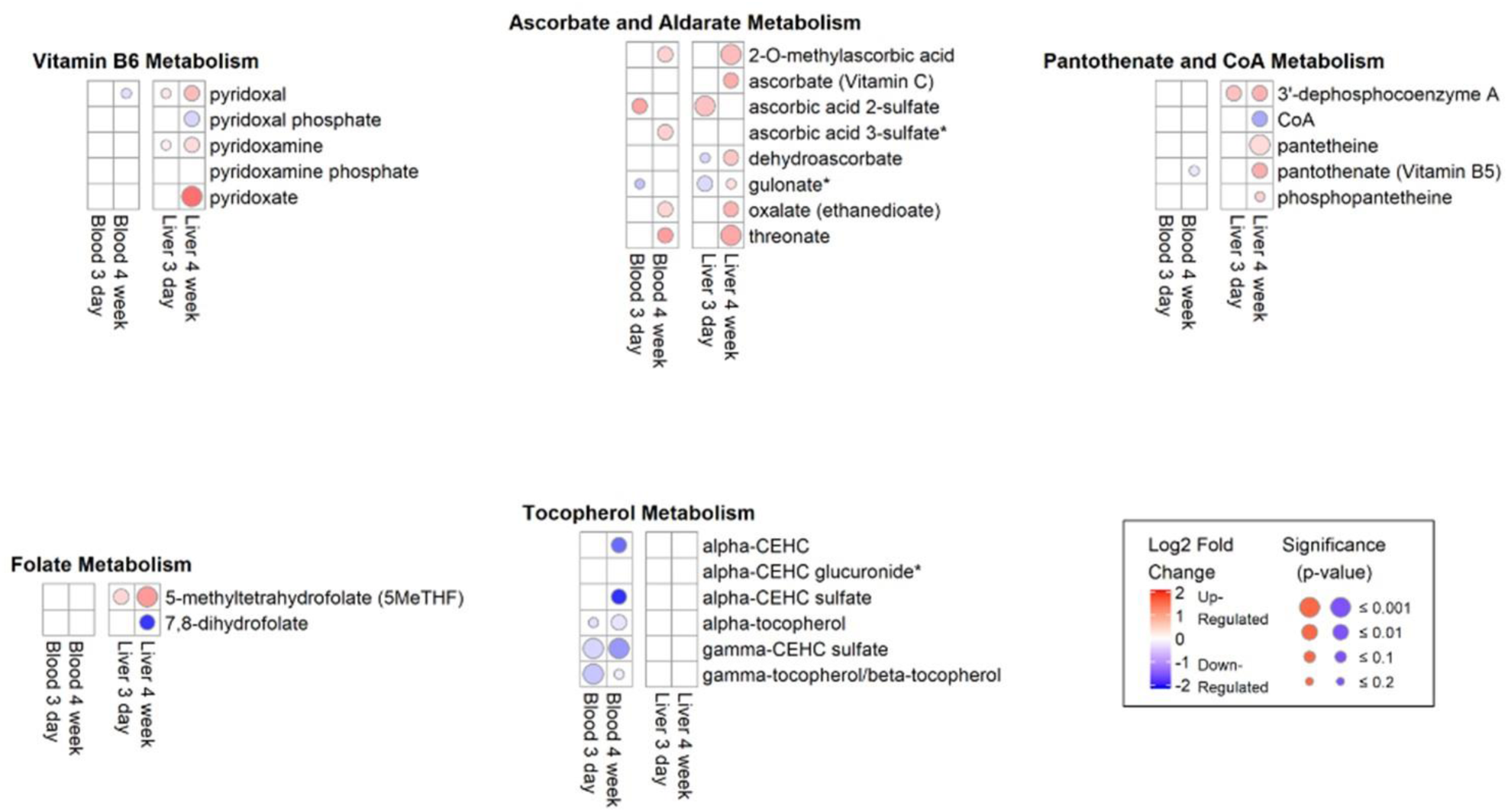
Changes in vitamin and cofactor metabolites in blood serum and liver tissue at 3 days and 4 weeks between ultrasound treated vs control ZDF animals.

**Supplemental Figure 18.**
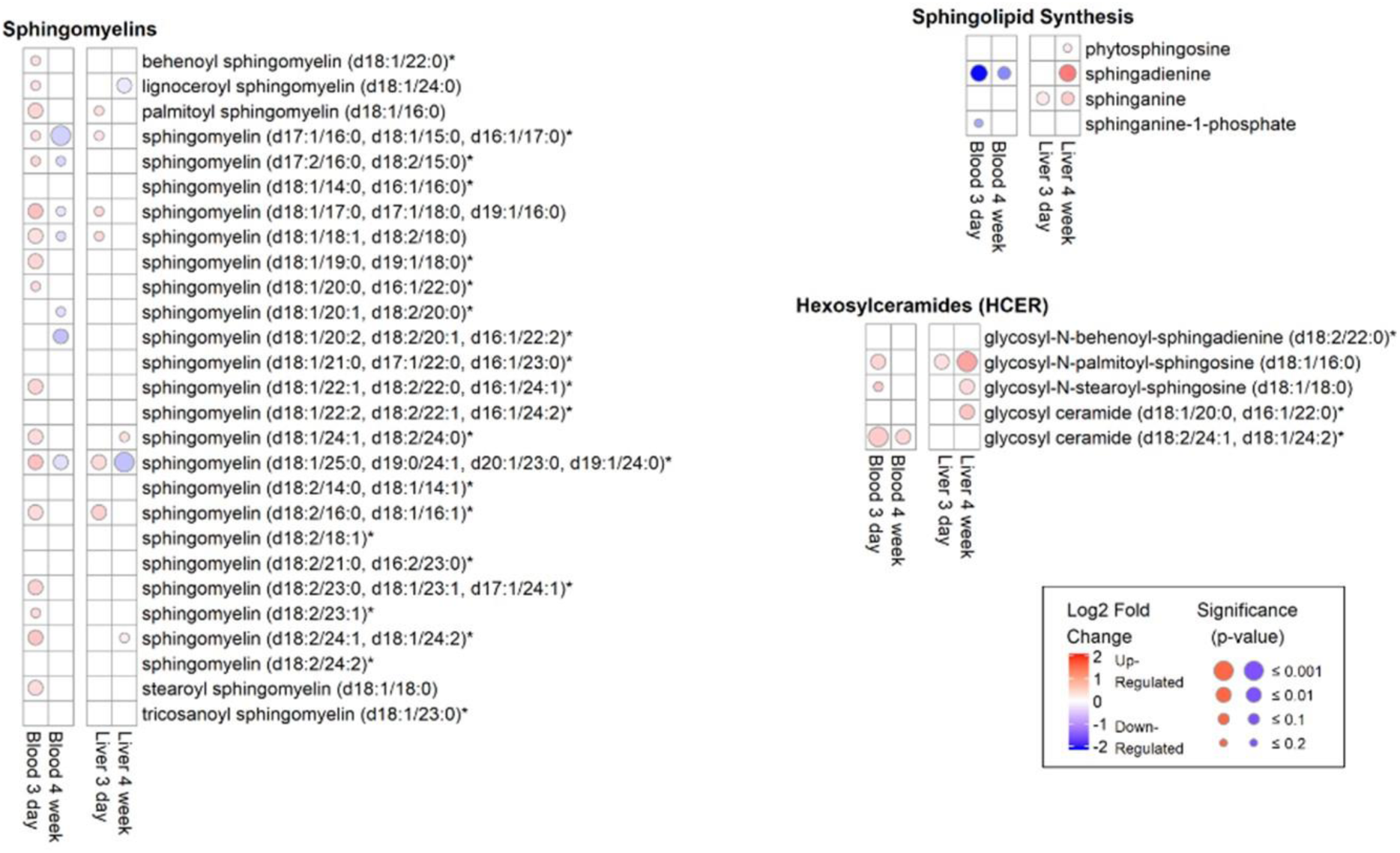
Changes in Sphingomyelins, hexosylceramides (HCER), and metabolites involved in sphingolipid synthesis in blood serum and liver tissue at 3 days and 4 weeks between ultrasound treated vs control ZDF animals.

**Supplemental Figure 19.**
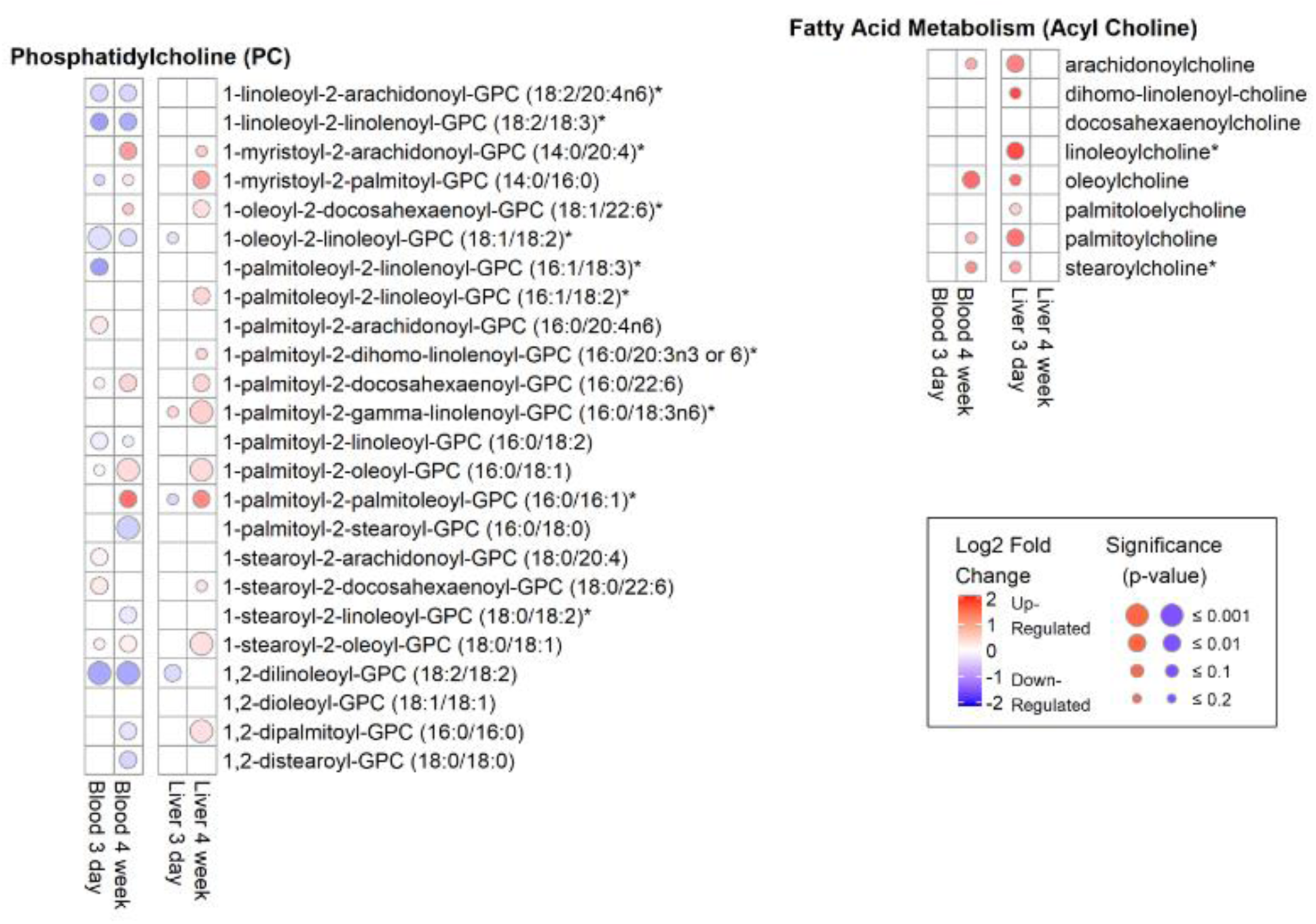
Changes in phosphatidylcholines (PC) and acyl cholines from fatty acid metabolism in blood serum and liver tissue at 3 days and 4 weeks between ultrasound treated vs control ZDF animals.

**Supplemental Figure 20.**
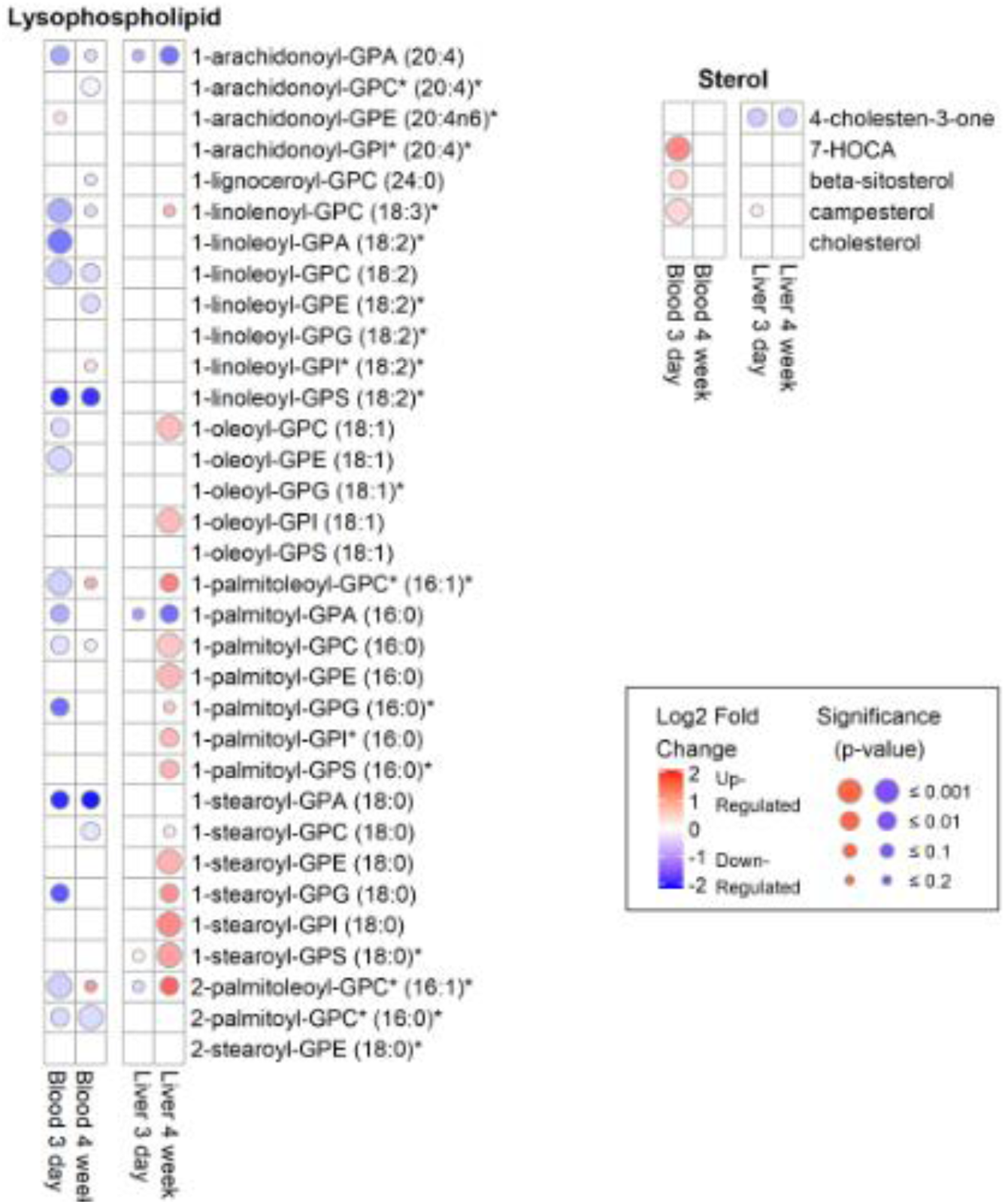
Changes in lysophospholipids and sterols in blood serum and liver tissue at 3 days and 4 weeks between ultrasound treated vs control ZDF animals.

